## Supplemental materials for "Mitochondria fragment and reassemble to initiate the formation and development of the nucleus"

**Fig. 4. Nutrient supplementation reversed nuclear formation of the organelles to mitochondrial recovery and re-establishment.** TEM was performed on five K562 cells at 5 min (**A-D**) and 15 min (**E**) time points. (**A**) High magnification image of the inset reveals that the aggregation of dense particles (opposite red, black and white arrowheads) induced by new medium was going to completely separate the nuclei (SN1 and SN2) (**A1**). (**A2-A4**) In contrast to the nutrient-limited condition, the assembling of dense particles blocked mitochondrial fragmentation, concomitantly inducing the return of dense particles for mitochondrial recovery (em1-em3) and re-establishment (such as em4-em10) (red arrowheads). Their return (opposite red arrowheads) discontinued the attachment of MN1 to either SN1 or MN2, concurrently obstructing nuclear growth. The reinstatement of em9 occurred via the recombination of the dense part (em9/1, small red arrow and black arrowhead) and that of lucent em9/2 (small red arrow and black arrowhead), whereas the return of dense particles from the surroundings (red arrowheads) re-established the electron-lucent em10. (**B**) The assembling of dense particles (small red arrowheads) induced by nutrient supplementation discontinued nuclear growth and resulted in the formation of bodies with mitochondrial morphology (NB1-NB4 and opposite red arrowheads) at the edge of the nucleus (**B1** and **B2**). (**C**) Upon challenge with new medium, nuclear fusions (SN1-SN3 and MN1-MN3) were obstructed, dense particles returned (red arrowheads, em4/1-em6/1) for mitochondrial renewal (em1-em3) and re-establishment (em4-em7), and aggregations of dense particles divided MN1 (em5/1 and em6/1; red arrow, small red and black arrowheads) into fragmented mitochondria that surrounded it (em1, em2, em4/2-em6/2). SMLB (black and white arrowheads) formation and mitochondrial recombination (em4/1-em6/1 and em4/2-em6/2) benefited the regeneration of completely fragmented mitochondria (**C1** and **C2**). (**D**) When the nucleus was at its early stage of formation, nutrient supplementation reversed the fragmentation of the organelles to promote mitochondrial recovery (em1-em4) and re-establishment (such as em5-em9 and opposite red arrowheads) (**D1-D4**). (**E**) The nucleus in this cell was small and oval shaped (NN), and new medium led to the aggregation of dense particles within it and at its edge. The dense particles concurrently aggregated to recover em1 (red arrowheads). At the nuclear edge and within em1, VLGs (small red arrowheads) appeared, which benefited mitochondrial re-establishment in the cytoplasm, similar to ERLs (white arrows) and SMLBs (black and white arrowheads). em2 was dense and elongated, and the return of dense particles transiently assembled into LMM (red arrows), which divided into em1. The internal assembling of dense particles began to separate these organelles (em2/1-em2/6) and provided dense particles (red arrowheads) for the regeneration of additional mitochondria and for the recombination of em2/3 and em2/4, dispersing the aggregates in lucent em2/5 (red arrow and black arrowhead) and reassembling ink masses (large red arrow) for mitochondrial development. (**F**) Schematic mechanism of mitochondrial assembling of dense particles initiated the nuclear formation and development.

**Figs. S1 to S42**

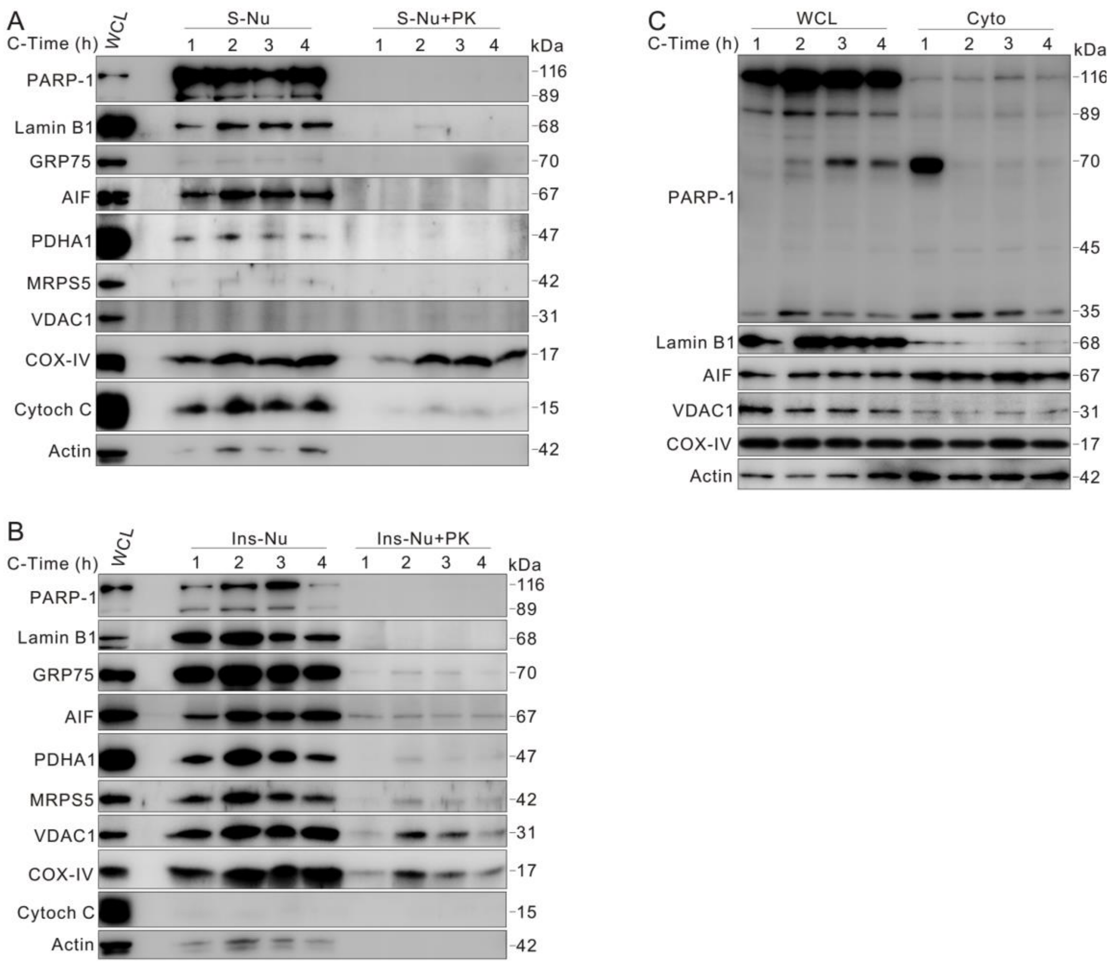


**Fig. S1. Mitochondrial resident proteins were present in the nuclear fraction.** Subcellular fractionation was performed after incubating K562 cells for 4 h, and the nuclear fractions were treated with or without proteinase K (PK, 200 ng/ml) for 30 min at room temperature and immunoblotted with the indicated antibodies. S-Nu: soluble nuclear fraction; Ins-Nu: insoluble nuclear fraction; WCL: whole cell lysate; Cyto: cytoplasmic fraction.

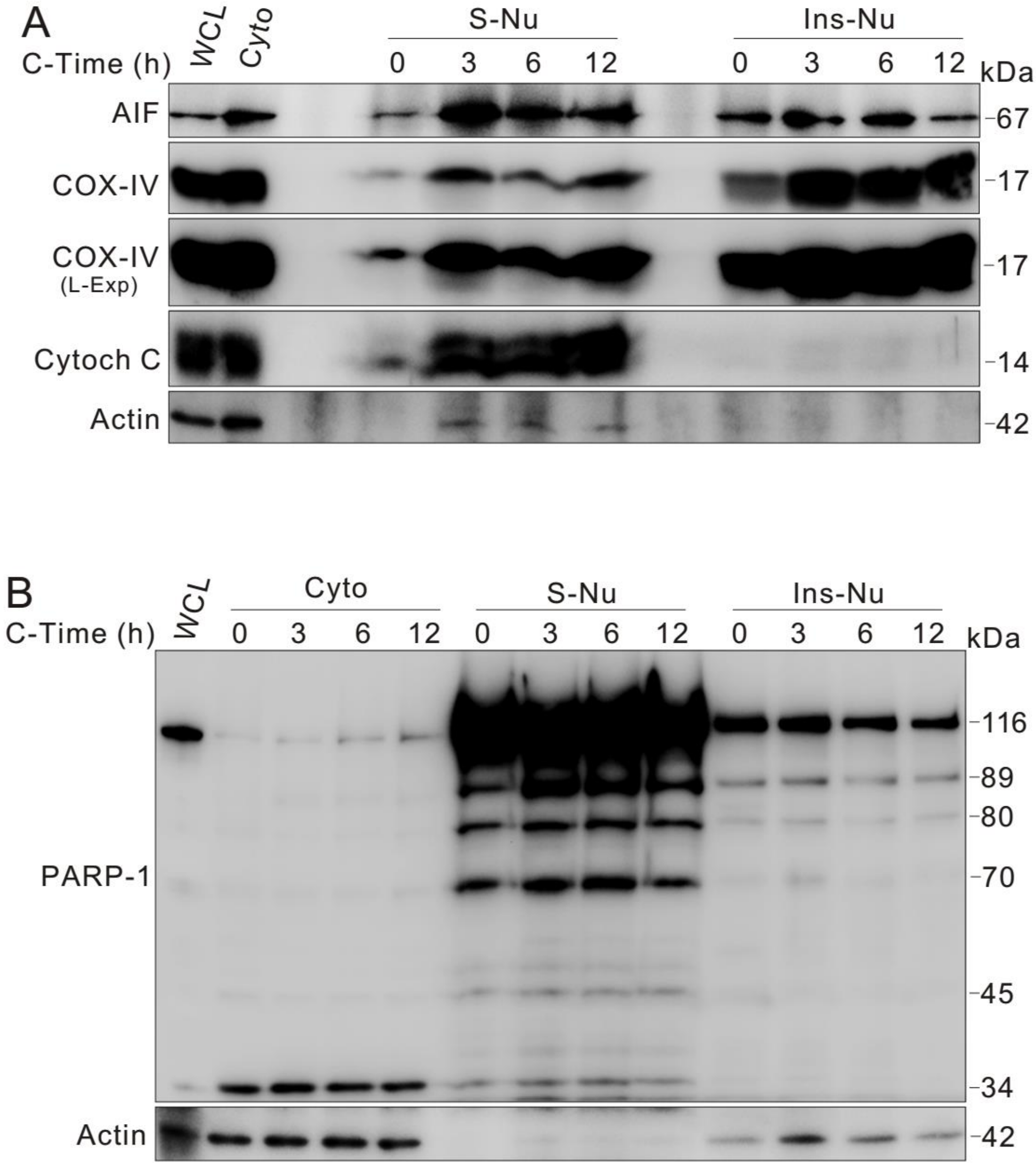


**Fig. S2. Cytochrome C, cytochrome C oxidase subunit IV (COX-IV) and apoptosis-inducing factor (AIF) are localized in the nucleus.** Nuclear fractions were extracted from K562 cells following incubation for 12 h and were analyzed by immunoblotting with the indicated antibodies. L-Exp: long exposure.

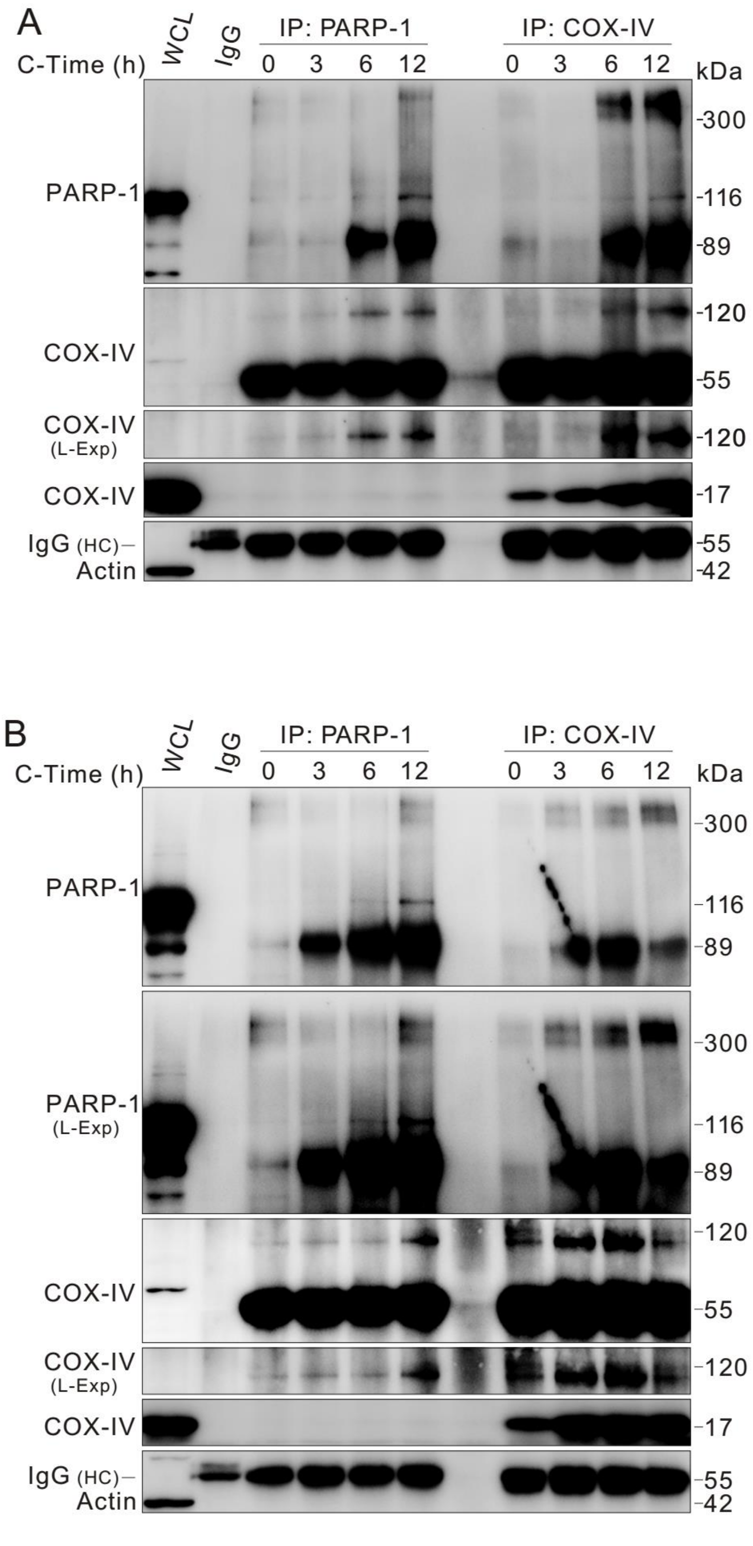


**Fig. S3. COX-IV interacts with PARP-1 in K562 cells.** Both soluble (**A**) and insoluble (**B**) nuclear fractions were isolated after incubating K562 cells for 12 h, and these fractions were lysed and subjected to immunoprecipitation using antibodies against either COX-IV or PARP-1. The immunoprecipitates were resolved by electrophoresis and probed by immunoblotting with the indicated antibodies. The results revealed that COX-IV and PARP-1 were able to form a high molecular weight complex (> 300 kDa), and the antibody against PARP-1 was unable to pull down the monomer of COX-IV. Mouse IgG2b was used as a negative control. HC: heavy chain.

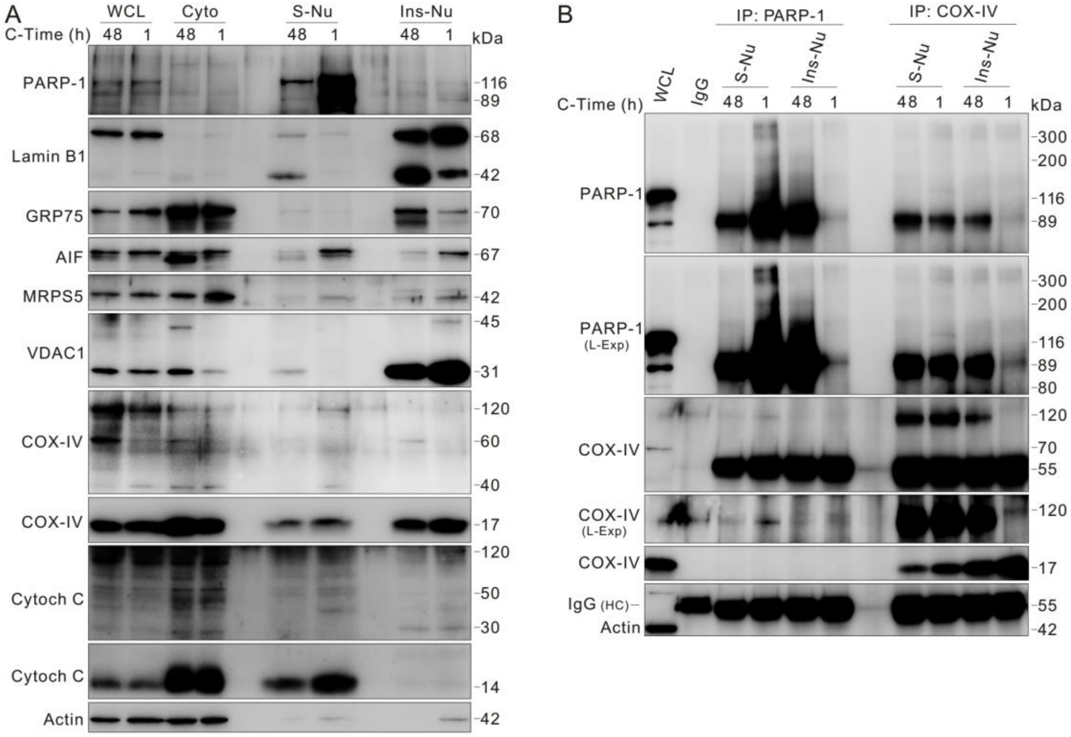

**Fig. S4. COX-IV interacts with PARP-1 in HeLa cells.** (**A**) HeLa cells were cultured for 48 h without changing the medium, and then the cells were split, moved to new medium and incubated for 1 h. Subcellular fractionation was performed, and the lysates of the WCL, Cyto, and nuclear fractions were subjected to immunoblotting with the indicated antibodies. (**B**) Reciprocal immunoprecipitation was carried out by using antibodies against either COX-IV or PARP-1. The immunoprecipitates were resolved by electrophoresis and probed by immunoblotting with the indicated antibodies. PARP-1 was not able to pull down the monomer of COX-IV, and PARP-1 easily formed a high molecular weight complex (> 300 kDa) with COX-IV at the 1 h time point in the soluble nuclear fraction.

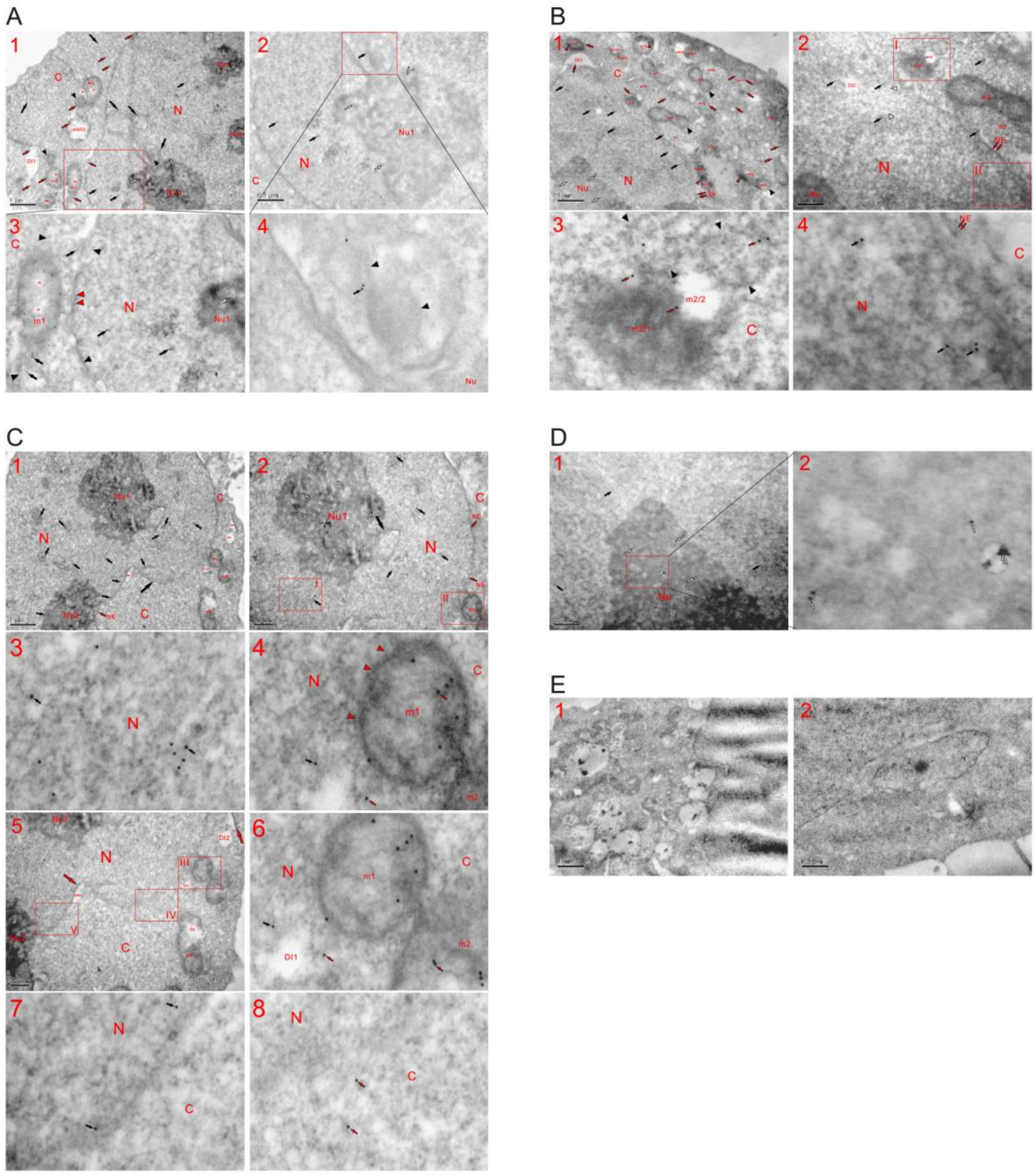


**Fig. S5. The mitochondrial protein glucose-regulated protein 75 (GRP75) labeled with gold particles (gold/GRP75) stained the nucleus and nucleolus.** Immunotransmission electron microscopy (TEM) was performed to visualize four K562 cells (12 h time point), and the micrographs of sections showed that GRP75 labeled with gold particles stained the nucleus (small black arrows), nucleolus (Nu) (small white arrows) and cytoplasmic mitochondria (small red arrows). In the cytoplasm and nucleus, the heavily or completely fragmented organelles possessed much less gold/GRP75 (large black arrows) than normal mitochondria (**A-D**). (**A**) In the cytoplasm, both relatively normal and fragmented mitochondria were stained by gold/GRP75 (small red arrows), and electron-dense mitochondria (m1 and m5) were dispersed in the nucleus. Mitochondrial fissions (m2 and m3) led to the formation of small mitochondrion-like bodies (SMLBs) (black arrowheads), and the heavily fragmented m4 showed little gold/GRP75 staining. The assembling of gold/GRP75-containing dense materials or particles at the periphery caused the matrix of em5 to become electron transparent and caused the mitochondrion to change into a lucent oval-shaped structure (DI1) and the disappearance of gold/GRP75 (small red arrow), and similar dilute phases (DI) appeared in other the organelles (m1-m3, DI). At the nuclear edge, completely fragmented mitochondria lost their morphology and dispersed into the nucleus (opposite black arrowheads and black arrows). Much fewer gold particles were present in the nucleus than in the cytoplasm, and gold/GRP75 stained heterochromatin (small black arrows) and Nu (small white arrows). Next to Nu1, fragmented organelles were observed (black arrows and arrowheads), which were stained with gold/GRP75 (**A1-A4**). (**A3)** and (**A4**): high magnification images of the insets in (**A1**) and (**A2**), respectively. (**B**) Mitochondrial fissions caused the organelles to diffuse into the nucleus (such as m1-m3 and DI1) and to be separated (such as m4-m8 and black arrowheads) while dispersing into each other (such as em4 and em1). m5 separated into dense m5/1 and electron-transparent m5/2. The aggregation of gold/GRP75 (small red arrowheads) appeared to occur within mitochondria (m3, m6/2, m9 and m10), and the morphology of completely fragmented mitochondria was lost (DI1, m1, m2, m3/2, m5, m6/2, m11-m16 and m17/2). m2 dispersed into the electron-opaque m2/1 and electron-lucent m2/2, and similar electron-transparent areas or spots existed in the nucleus (DI2 and white arrowheads). Adjacent to m2, gold/GRP75 (small red arrows) stained a completely fragmented mitochondrion (opposite black arrowheads), and the gold particles (small black arrows) appeared below the NE (double small red arrows); NE: nuclear envelope (**B1-B4**). (**B3** and **B4**): high magnification images of the inset I and II in (**B2**), respectively. (**C**) Next to Nu1 and at the nuclear edge, fragmented mitochondria (m1, DI1, DI2 and large black arrows) and dense parts of the organelles dispersed into the nucleus, in which an area enriched with gold/GRP75 was observed (**C3** and small black arrows), and fusion between the nucleus and m1 was exhibited (**C4** and red arrowheads). Mitochondrial fission caused m2 to disperse into m1 and caused gold/GRP75 appeared on either side of DI1 (nuclear and cytoplasmic side) (**C6**, small black and red arrows). The gold particles also stained the nucleus below NE (small black arrows) and the cytoplasm at the nuclear edge (small red arrows) (**C7** and **C8**). (**C3** and **C4**): high magnification images of insets I and II in (**C2**), respectively. (**C6-C8**): high magnification images of insets I-III in (**C5**), respectively. (**D**) High magnification image of the inset reveals the gold-GRP75 staining in Nu (small white arrows); two small white arrows: a possible aggregate of gold particles (D1 and D2). (**E**) Two negative control cells, which were incubated without the first antibody (E1 and E2). N: nucleus; Nu: nucleolus; NE: nuclear envelope; C: cytoplasm; m:

mitochondrion; DI: dilute phase.

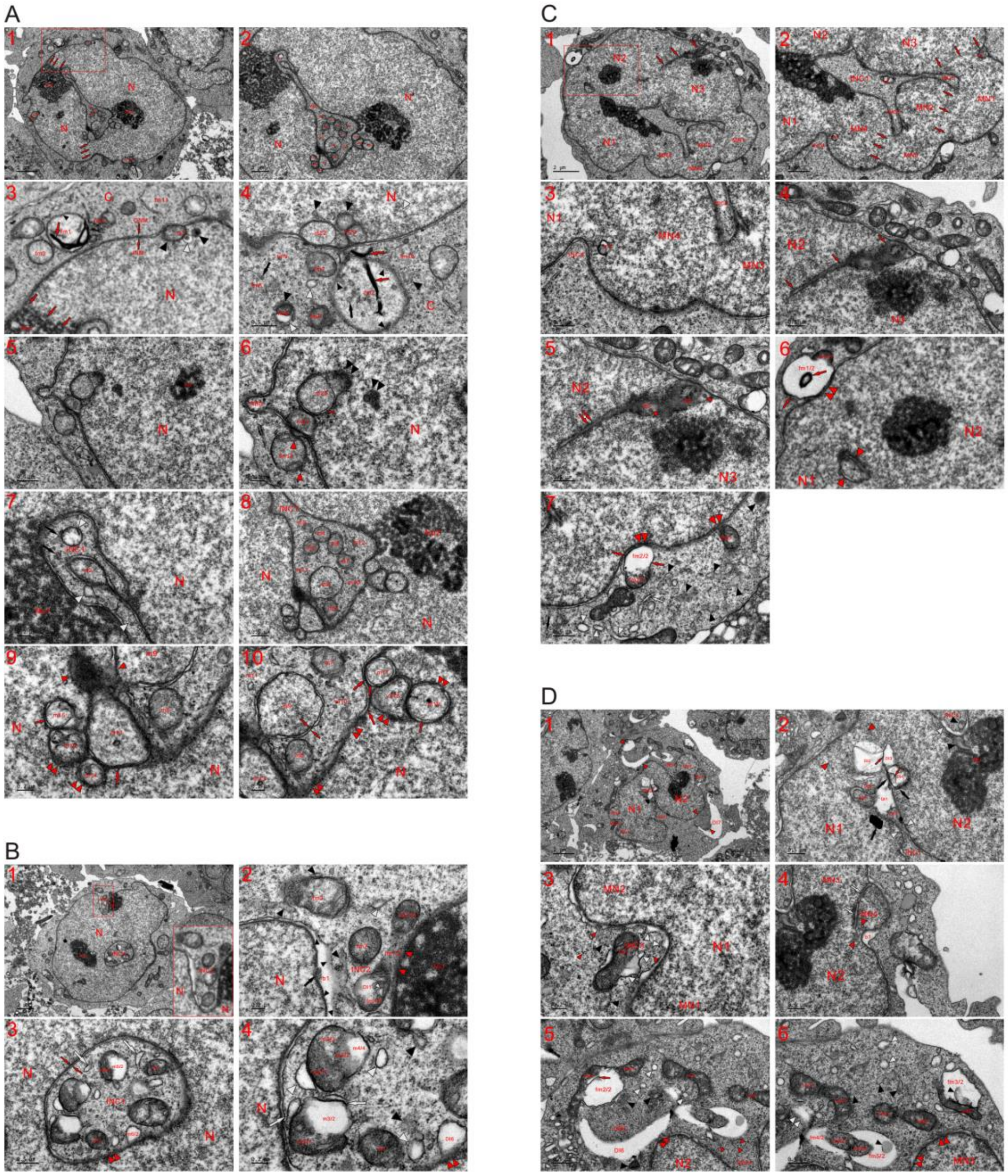

**Fig. S6. Mitochondria are located in the nucleus and within intranuclear inclusions (INCs).** TEM was performed on four K562 cells at the 4 h time point (incubating the cells with new medium for 4 h), and the micrographs reveal that both cytoplasmic mitochondria and nuclear-localized mitochondria underwent fragmentation and separation (**A-D**). (**A**) The same cell shown in **Figure 1A**. In the nucleus, a large intranuclear inclusion (INC), which contained mitochondria, and trace of nuclear fusion were observed (three abreast red arrows); in the cytoplasm (**C**), mitochondria were partially (such as fm4-fm7) or completely (such as fm8-fm11) fragmented, and the aggregation of dense particles to the periphery (small black arrowheads) with or without internal assembling (large red and black arrows) caused the organelles to become electron-transparent and to be dispersed (such as fm1-fm3). Below the NE (opposite small red arrows), mitochondrial fission occurred (m1, m22 and m23; white and black arrowheads), and the internal and external (their aggregation at the mitochondrial periphery) assembly of the particles caused m22 to become electron-lucent and to concurrently include itself in the nucleus as a nuclear mitochondrion before its dispersion to achieve nuclear conversion. Similar to m22, both m1 and m23 included themselves in the nucleus along with completing nuclear transition of the neighboring mitochondria. Small dense bodies (SDBs) (black arrowheads) and small dilute bodies (SDIBs) (white arrowheads) were derived during mitochondrial fissions, some of which looked like small mitochondria-like bodies (SMLBs) (**A1-A4**). (**A3**): a high magnification image of the inset in (**A1**). (**A5** and **A6**) At the nuclear edge, dense particles reassembled to form a nuclear bud (NBD), and fm12 fragmented into dense particles that diffused into the nucleus and the surroundings (red arrowheads). Both m20 and m21 became nuclear localized in a similar manner as m22, and the aggregation of dense particles at the periphery temporarily blackened the limiting membrane (LMM) of m22 (red arrowheads). Part of m21 had achieved nuclear transition; vestiges of mitochondrial fissions were observed within the nucleus (double black arrowheads), and the particles assembled into a Nu-like structure (Nu3). (**A7**) Traces of mitochondrial fragmentations are shown (black arrows and white arrowheads), and the density of m2 was similar to that of the nucleus. (**A8-A10**) Within INC1, partially (m3-m8) or completely (m10-m12) fragmented mitochondria were present, and m9 possessed a tubule-NE (opposite red arrows), looked like an MN and attached to the nucleus by the dispersed organelle (opposite red arrowheads). Heavily or completely fragmented mitochondria lost their morphologies (m3 and m10-m12), and the aggregation of dense particles caused the organelles to diffuse into the nucleus (double red arrowheads). Before complete fragmentations, the enclosed mitochondria (m13-m19) blackened the LMM (small red arrows) by assembling the particles to their peripheries, and similar assemblies simultaneously occurred in neighboring mitochondria to form dark strands (large red arrows); these strands were transient and dispersed into the nucleus. ONM: outer nuclear membrane; INM: inner nuclear membrane. (**B**) In this nucleus, there appeared the closed INC1 and opened INC2, whose formation was due to three-sided nuclear expansion (high magnification image of the inset) (**B1**). (**B2**) Within INC2, the aggregation of dense particles (black arrow and arrowheads) to the periphery caused the organelle to temporarily become a bubble (b1) at the nuclear edge and created dilute spots in fm1/1 (DI1 and small white arrowhead), and their assembling dispersed (fm1/2) or was going to disperse mitochondria (fm1/1, fm1/3, fm2, fm3, and black and white arrowheads) into dense particles, which entered Nu1 (red arrowheads). (**B3** and **B4**) During mitochondrial fissions within INC1, endoplasmic reticulum (ER)-like structures (large white arrowhead), SDBs (black arrowheads) and SDIBs (white arrowheads) temporarily formed, and some SDIBs changed into vesicles by assembling dense particles at the periphery. The aggregation of the particles first condensed mitochondria (m1 and m2) and then caused the organelles to separate into dense (m3/1-m5/1, m4/2) and electron-lucent (m3/2, m4/4 and m5/2) or electron-dilute (m4/3) parts; m1 could evolve into the appearance of DI6 to fuse with the nucleus (double red arrowheads) through the aggregation of dense particles to the periphery. Opposite red arrows: aggregation of dense particles from neighboring mitochondria transiently formed a tubule-NE fragment. (**C**) Traces of nuclear fusions (N1-N3 and MN1-MN4) are shown (m1, m2, INC2-INC4 and three abreast red arrows). Within INC1, almost all mitochondria were completely fragmented, and only part of m3 was observed (**C1-C4**). (**C5**) Between nuclei (N2 and N3), incomplete nuclear merging was demonstrated (double red arrows), and mitochondrial fissions occurred due to the assembly of dense particles (m4, m5 and small red arrowheads). (**C6** and **C7**) Incomplete nuclear transition of a mitochondrion appeared between N1 and N2 (opposite red arrowheads). In the cytoplasm, the aggregation of dense particles caused the organelles to separate into electron-opaque (fm1/1 and fm2/1) and electron-transparent (fm1/2 and fm2/2) parts, and their internal assembly occurred in fm1/2 (large red arrow). The aggregation of the dense particles to the periphery transiently blackened the LMM (small red arrows) and caused the transparent parts to fuse with the nucleus (double red arrowheads). The condensed fm3 assembled the particles into the nucleus and fused with it (double red arrowheads). Large white arrow: ERL; black arrow and arrowheads: SDBs; white arrowheads: SDIBs. (**C6**): a high magnification image of the inset in (**C1**). (**D**) Partial nuclear fusion (opposite red arrowheads) between N1 and N2 partitioned the cytoplasm to form a large INC, which further divided into INC1 and INC2 by merging in the middle of the nuclei (opposite white arrowheads). Within INC3, the aggregation of dense particles caused mitochondrial fissions to concurrently form dense structures (DE1-DE3, red and black arrows) and electron-lucent DIs (such as DI1-DI3). Within INC4 and adjacent to Nu, traces of mitochondrial fragmentation are shown (black and white arrowheads) (**D1** and **D2**). (**D3** and **D4**) Three-sided nuclear expansion (N1, MN1 and MN2) by mitochondria formed the small and shallow INC2 at the edge of N1, and within it, condensed (fm1) or fragmented mitochondria (black and white arrowheads) changed into nuclei via the rearrangement of dense particles (small red arrowheads). Mitochondrial assembling of the particles (opposite red arrowheads) to link MN4 and N2 caused part of the organelle to become nuclear bubble (b1). The aggregation of dense particles enlarged both micronuclei (MN3 and MN4) to cause their fusion, and incomplete nuclear transition of mitochondria was observed between them (white arrows). (**D5** and **D6**) Mitochondria assembled dense particles to condense (fm2/1-fm5/1 and fm2/3) and separate concomitantly occurring the internal aggregation (red arrow and black arrowheads) and forming electron-transparent structure (fm2/2-fm5/2). These mitochondria dispersed through the continuous aggregation of the particles to periphery and disappeared via the consequent intermixing of dense particles in the cytoplasm (double white arrowheads). Dilute parts of fm7 and fm8, both of which were being dispersed by further assembling dense particles; a large lucent DI6 should have formed from more than one mitochondria as observed with fm4 and fm5. In addition to fusing with the nucleus (double red arrowheads), mitochondrial fissions dispersed the organelles into each other (DI6, fm2, fm7 and fm4) to encircle heavily or completely fragmented mitochondria (DE6, black and white arrowheads) or to enrich the dense particles at the opening and inside INC4 (small red arrowheads). By doing this, the organelles promoted the formation of new MNs (DI7 and opposite red arrowheads in **D1**) and caused the INC to disappear. Black arrowheads: SDBs; white arrowheads: SDIBs.

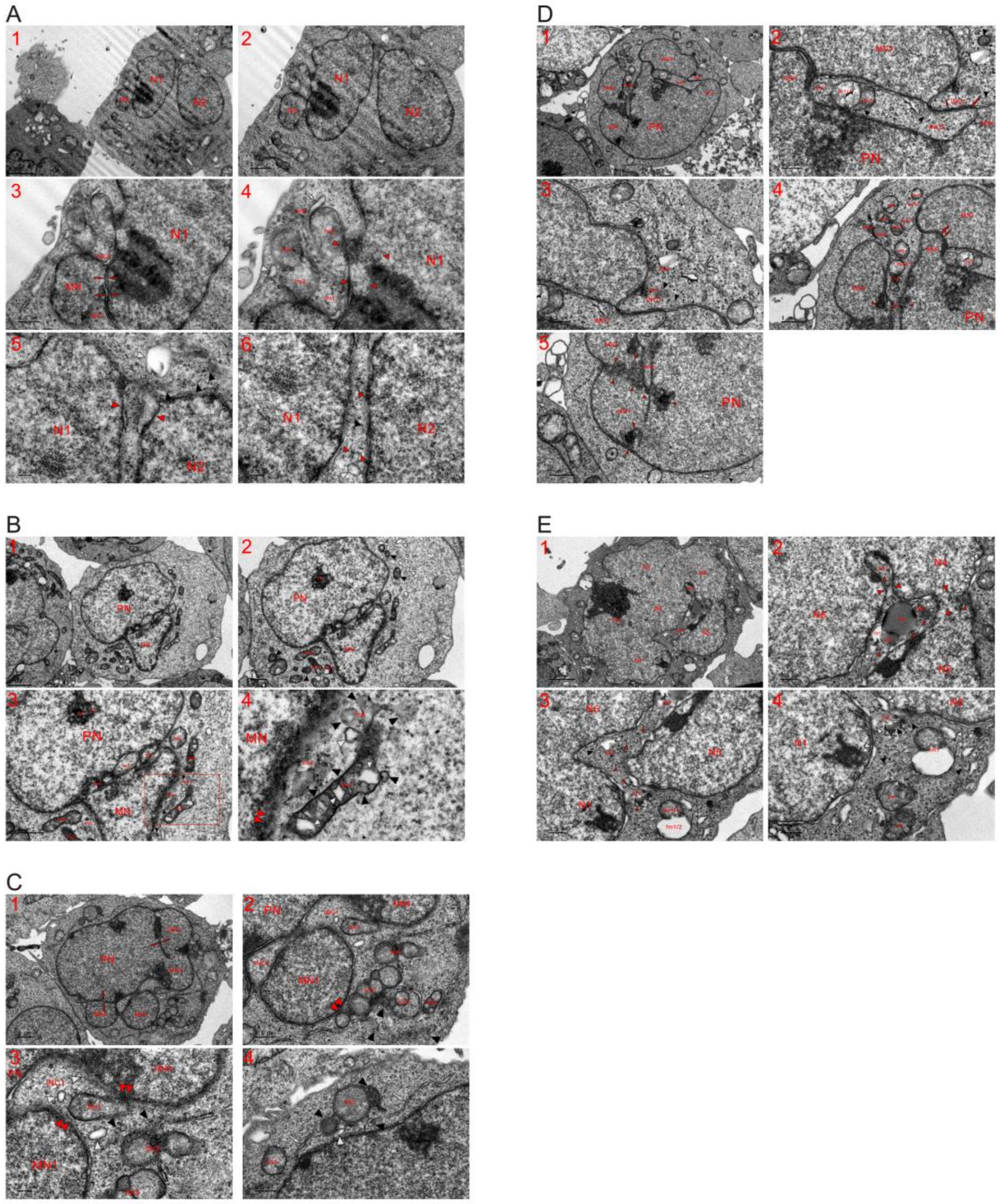


**Fig. S7. Mitochondria fragmented for nuclear transition to connect a MN with the PN.** TEM was performed on five K562 cells at the 2 (**A**) and 4 (**B-E**) h time points, and mitochondria achieved nuclear transition to join together the separately-formed MN with the PN by first turning into dense particles, and attachment of the MN partitioned cytoplasm to shape the INC. (**A**). This cell possessed two large nuclei (N1 and N2) and an MN, all of which were separately established and then merged (**A1** and **A2**). (**A3** and **A4**) Along with the nuclear growth induced by mitochondrial fragmentation, cytoplasmic space between N1 and the MN narrowed, nuclear fusion occurred, and the aggregation of dense particles of neighboring mitochondria formed nuclear tubule-like structures (opposite red arrows). Partial nuclear fusion partitioned the cytoplasm to shape INC1 and INC2, in which fragmented mitochondria were dispersed into dense particles for nuclear transition (such as fm1-fm5), and at the edge of N1 and Nu, traces of mitochondrial fragmentation or mitochondrial assembling of dense particles for nuclear development were observed (opposite red arrowheads). White arrow: ERL; white arrowheads: SDIB. (**A5** and **A6**) Between nuclei (N1 and N2) and at their edges, mitochondria heavily or completely fragmented into dense particles (small and opposite red arrowheads), SDBs and SDIBs (black and white arrowheads), and both SDBs and SDIBs eventually became particles. (**B**) Partially (fm1 and fm2) or completely (fm3) fragmented mitochondria (such as fm1-fm3 and fm9) existed between the MN and PN, and outside of the nucleus, the organelles fragmented into dense particles that assembled for nuclear enlargement and growth (fm4-fm11, black and white arrowheads). The internal assembling of the particles formed electron-lucent spots in fm7 (small white arrowheads), which fused with the MN (double red arrowheads) and diffused into the neighboring organelles (fm4 and fm5) to provide dense particles. During mitochondrial fissions, SDBs (black arrowheads), SDIBs (large white arrowheads) and ERLs (white arrows) were intermediately generated (**B1-B4**). Neighboring the Nu, incomplete nuclear transition of a mitochondrion was observed (m1). (**B4**): a high magnification image of the inset in (**B3**). (**C**) Nuclear fusions resulted in the combination of four MNs (MN1-MN4), and in the cytoplasm, almost all mitochondria became electron opaque with a density similar to that of the nucleus (fm1-fm5). The attachments of MNs to the PN and the fusion of MNs (MN1 and MN2) partitioned the cytoplasm to form INC1 and INC2, and the organelles dispersed into nuclei (double red arrowheads) and into each other (fm1-fm3, black and white arrowheads) to close INC1. Fragmented mitochondria further assembled dense particles to intermediately form ERLs (large white arrows), SDBs (black arrowheads) and SDIBs (white arrowheads), and all the structures eventually dispersed into dense particles in the cytoplasm. (**D**) In addition to the PN, four MNs were observed; these MNs were individually formed and either attached (MN4) or partially fused with the PN (MN1-MN3). The formation of nuclei decreased the space in the cytoplasm and compartmentalized the cytoplasm to form INCs (INC1-INC3). The formation of an NPB and dark strand by dense particles divided the cytoplasm into INC1 and INC2; INC1 was closed, and within both INCs, partially (m1/1 and m1/2) or completely (m2, fm1, fm2, and black and white arrowheads) fragmented mitochondria were observed (**D1-D3**). (**D4** and **D5**) Mitochondria completely fragmented (m3, opposite red arrowheads) to join together individually-formed nuclei (MN1 and MN2; MN1 and PN; MN2 and PN), and m3 dispersed into dense particles to partition INC3 (INC3/1 and INC3/2), which contained the less fragmented m4 and m5. At the opening of INC3/1, mitochondrial fissions caused the organelles to diffuse into each other (fm3-fm7) and into nuclei (MN2 and MN3). Between MN1 and PN, the simultaneous assembly of dense particles from neighboring mitochondria formed nuclear tubules (white arrows) that indicated incomplete nuclear merging. (**E**) Nuclei (N1-N6) that should have been separately formed were joined together to build a large nucleus, and partial nuclear fusion occurred between N4 and N5 (opposite red arrowheads). Through mitochondrial fissions driven by the assembly of dense particles (small red arrowheads), mitochondrion further divided the enclosed cytoplasm (INC1 and INC2) (opposite red arrowheads) with causing itself to become the appearance of an INC (INC1). Within INC2, mitochondria (such as m2-m4) fragmented to intermediately become dense bodies (DE1, DE2 and black arrowheads) and electron-lucent structures (DI1 and white arrowheads) before totally dispersed into dense particles (small red arrowheads). At its opening, mitochondrion was dispersed into particles (m1) for nuclear development. In the cytoplasm, the aggregation of dense particles separated fm1 into dense fm1/1 and electron-transparent fm1/2, and condensed mitochondria (fm2 and fm3), SDBs (black arrowheads) and SDIBs (white arrowheads) were going to disperse into dense particles. Formation of SDIBs was driven by the assembling of particles to the peripheries. After assembly at edge of nucleus, SDIBs looked like nuclear bubbles (double white arrowheads).

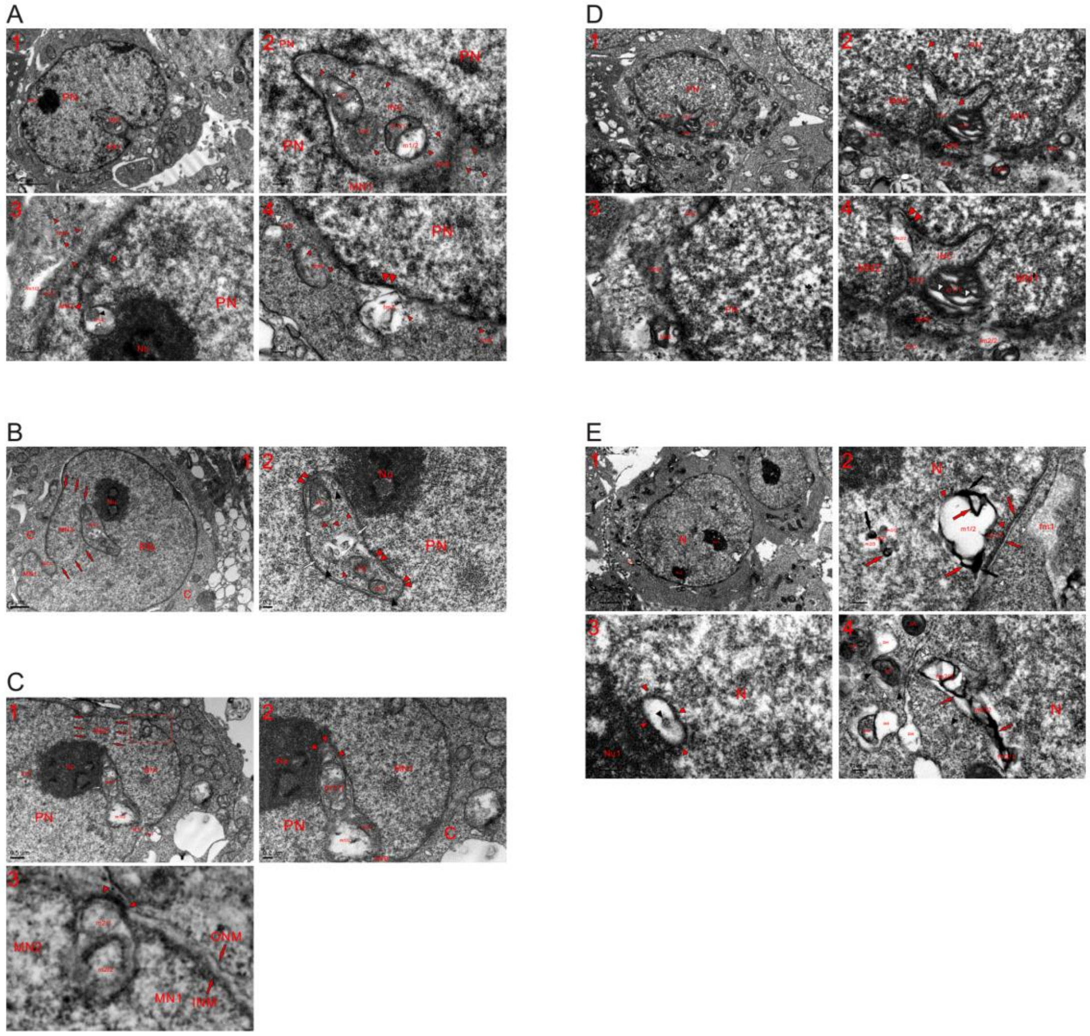


**Fig. S8. Nuclear localization of mitochondria in HEK293T cells.** TEM was performed on five HEK293T cells at different time points (**A**: 1 h; **B** and **C**: 3 h; D: 12 h; **E**: 48 h), and the micrographs reveal that mitochondria localized within the nucleus and in INCs (**A-E**). (**A**) Three-sided nuclear expansion (PN and MN1) compartmentalized the cytoplasm into the INC, which was subsequently sealed by the formation of a nucleoplasmic bridge (NPB) through the mitochondrial assembling of dense particles (small red arrowheads). Within the INC, completely fragmented mitochondria had dispersed into the particles (small red arrowheads) and less fragmented ones were observed (m1/1 and m1/2; m2; m3 and small white arrowheads)

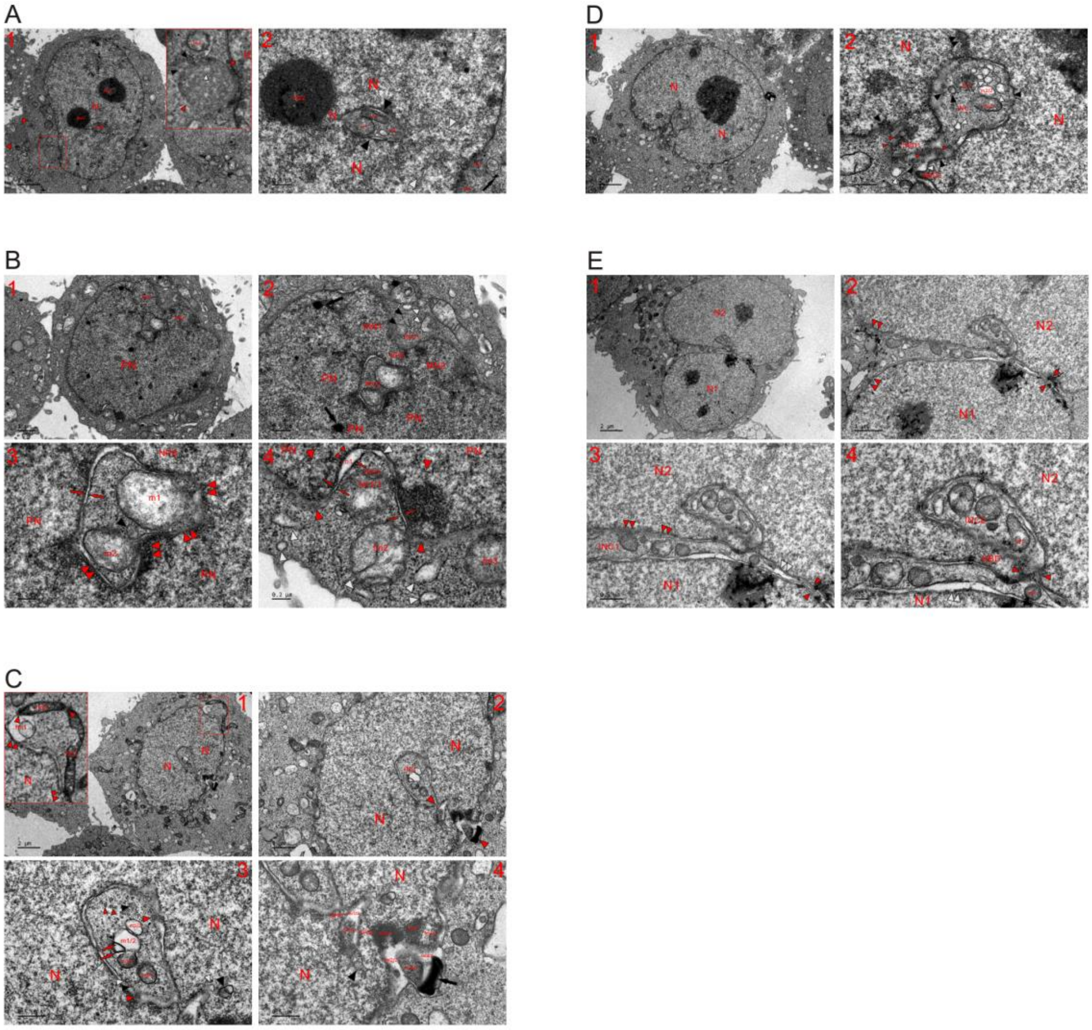

**Fig. S9. Nuclear localization of mitochondria in HeLa cells.** TEM was performed on five HeLa cells at 1 and 6 h time points (**A** and **B**: 1 h; **C-E**: 6 h), and the micrographs reveal that mitochondria were only localized in INCs (**A-E**). (**A**) High magnification image of the inset reveals that fragmented mitochondria (fm3, black and white arrowheads) grouped into a large spherical body (opposite red arrowheads), which tended to change into an MN by assembling dense particles. Four-sided nuclear formation or growth (N, Nu2 and opposite white arrowheads) via mitochondrial assembling of dense particles (such as m1-m3, fm1 and fm2) reduced the space of enclosed cytoplasm and formed to a closed INC (opposite black arrowheads), whose formation was merely due to the incomplete nuclear transition of the organelles (m1-m3). (**B**) Three-sided nuclear formation (PN, MN1 and MN2) partitioned the cytoplasm, which was further divided by the newly formed NPB to form an opened INC1 and the closed INC2, concomitantly causing the organelles to become nuclear localized (such as m1 and m2). Within INCs, mitochondria fragmented (m1, m2, black and white arrowheads) to disperse into the nucleus in the form of dense particles (double red arrowheads). The assembly of particles from neighboring fragmented mitochondria shaped the tubule-NE fragment (opposite small red arrows). Large black arrows: ink dots, which formed by the aggregation and compaction of dense particles and represented the incomplete mixture of fragmented mitochondria and the nucleus (**B1-B3**). (**B4**) At the nuclear edge, a small and shallow INC3 was observed, whose formation was merely due to incomplete nuclear transition of a mitochondrion (fm1/1 and b1) and its neighboring counterparts having been accomplished nuclear evolvement (opposite red arrowheads). Between internal (fm1/1) and external (small red arrowheads) aggregation of dense particles, lucent structures were formed (b1 and white arrowhead). The transient formation of tubule-NE fragments (opposite red arrows), which formed by the simultaneous assembly of dense particles from the neighboring organelles to the peripheries or assembling of the particles from fm1 itself, falsely caused the formation of INC3 to look like an invagination of the cytoplasm. The aggregation of dense particles led to mitochondrial fissions or fragmentations (fm2, fm3 and white arrowheads), which consequently and eventually dispersed the organelles into the particles for nuclear development. (**C**) High magnification image of the inset reveals that the aggregation of dense particles diluted fm1 and condensed mitochondria (fm2 and fm3), which dispersed to contact each other (red arrowheads) and to fuse with the nucleus (double red arrowheads), and by doing so, the enclosed cytoplasm accelerated for nuclear development (**C1**). (**C2-C4**) The INC was going to be closed (opposite red arrowheads) by fragmented mitochondria (such as fm1/1-fm4/1, fm1/2-fm4/2, black arrow, black and white arrowheads), and from both sides, mitochondria-derived dense particles first assembled into two NPBs (NPB1 and NPB2), both of which enlarged and disappeared during the nuclear transition of the neighboring mitochondria. Within the INC, the mitochondrial aggregation of dense particles (small red arrowheads) led to the separation of the organelles (m1/1, m1/2, m2/2, m3/1 and red arrows) and to the formation of virus-like particles (large red arrowheads), all of which eventually dispersed into particles for nuclear evolvement. Within the nucleus and adjacent to the INC, incomplete mitochondrion-to-nucleus transition was observed (black and white arrowheads). (**D**) The INC was being closed by newly built NBDs (NBD1 and NBD2) and mitochondrial assembling of dense particles for nuclear development (small red, black and white arrowheads), and following the nuclear transition of the fragmented organelles, the NBDs turned into the attached micronuclei and joined together. Within the INC, mitochondria fragmented to temporarily form SDBs (black arrowheads) and SDIBs (white arrowheads), some of which changed into vesicles by the assembly of dense particles to the peripheries. The internal assembly of the particles caused m1 to look like an MVB, and their aggregation at its periphery dispersed the organelle. m2 assembled dense particles to separate (m2/1 and m2/2). (**E**) Partial nuclear fusion occurred (opposite red arrowheads) to N1 and N2, concurrently partitioning the cytoplasm to form INC1, and at its opening and within it, fragmented mitochondria dispersed into nuclei (double red arrowheads) and into each other (**E1-E3**). (**E4**) After the transition of the NBD into an NPB by fragmented mitochondria (m1, m2 and opposite red arrowheads), INC2 was closed. The formation of INC2 was not related to herniation or invagination of cytoplasm but was merely due to the nuclear transition of the mitochondria that surrounded it. At the edges of both N1 and N2, fragmented mitochondria continuously assembled dense particles to their peripheries, temporarily forming nuclear bubbles (double white arrowheads).

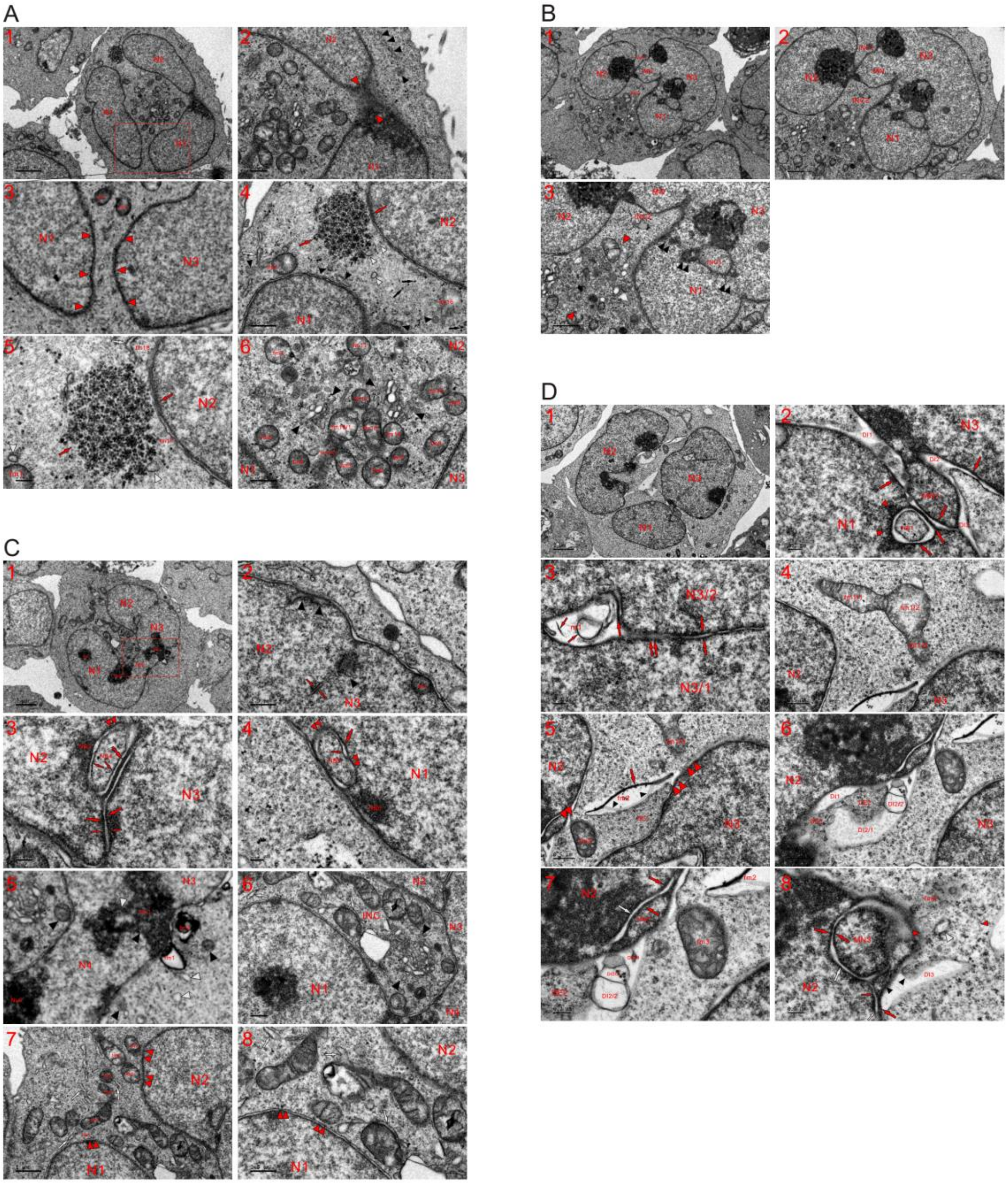


**Fig. S10. Separately-formed nuclei joined together to partition the cytoplasm into a large INC.** TEM was performed on four K562 cells at the 4 (**A-C**) and 8 (**D**) h time points, and the micrographs showed that more than one nucleus was individually and simultaneously built in a single cell, and their combination partitioned the cytoplasm to form a large INC (**A-D**). (**A**) In this cell, three individually formed nuclei (N1-N3) were being joined together by fragmented mitochondria (DE1, DE2 and opposite red arrowheads), and the aggregation of dense particles condensed the organelles (DE1 and DE2), which had lost the LMM and were dispersed into particles via their further assembly (**A1-A3**). (**A3**): a high magnification image of the inset in (**A1**). (**A4-A6**) Mitochondrial-derived dense particles aggregated into glycogen-like granules (black arrows), which grouped between N1 and N2 (opposite red arrows), and partially (such as fm1-fm14 and fm15/1) or completely (fm15/2 and fm16-fm18) fragmented mitochondria existed between or among nuclei. During mitochondrial fissions, ERLs (white arrows), SDBs (black arrowheads) and SDIBs (white arrowheads) were transiently formed, and fissions of the organelles led to the formation of Golgi complex-like structures (opposite black arrowheads). (**B**) Formation of an MN linked the large nuclei (N1 and N3) concurrently compartmentalizing the cytoplasm to form INC1 and INC2, and traces of nuclear fusion or incomplete nuclear transition of mitochondria were observed between N1 and N3 (INC3 and double black arrowheads). At the opening of INC2, fragmented mitochondria formed a group, which linked nuclei (N1 and N2) and benefited the nuclear transition of the organelles (opposite red arrowheads) (**B1-B3**). (**C**) Between and within nuclei, the vestiges of incomplete nuclear conversion of mitochondria were observed (NB2-NB6; large, small and opposite red arrows; black arrowheads). The aggregation of dense particles at the periphery (the external assembling) from two neighboring mitochondria formed nuclear tubules (opposite red arrows), which usually consisted of dark strands (large red arrows). During external assembly, the internal aggregation and linearization of dense particles in one of the organelles shaped dark threads (small red arrows), which in turn created a structure of double nuclear tubules adjacent to NB4. The further assembly of dense particles occurred to make both dark strands and nuclear tubules disappear, leading the fragmented organelles to disperse into the nucleus (double red arrowheads). NB: nuclear body. (**C1-C4**). (**C5**) A high magnification image of the inset in (**A1**), showing that mitochondria-derived dense particles assembled to form Nu1 (such as fm1 and fm2; black and white arrowheads), which consequently or concurrently joined together N3 and N4. (**C6-C8**) Three-sided nuclear formation partitioned the cytoplasm to form a large and opened INC, which filled with fragmented mitochondria, and at its opening site, mitochondria assembled dense particles to disperse into each other and to fuse with the nucleus (double red arrowheads). Opposite black arrowheads: a Golgi complex-like structure, which was formed by the combined fragmentations of the organelles. (**D**) Three nuclei (N1-N3) were joined together to build a large one, the formation of MN1 linked the nuclei (N1 and N3) and the aggregation of dense particles formed dark strands (large red arrows) and created electron-transparent structures (DI1-DI3). Through the internal aggregation of dense particles, the organelle formed NB1 and a lucent ring-like structure, and their assembly at the periphery allowed it to disperse into N1 in the form of particles (red arrowheads) and transiently aggregate into dark strands (**D1** and **D2**). (**D3**) Nuclear fusion (N3/1 and N3/2) within N3 by mitochondrial aggregation of dense particles was observed (m1, opposite and double red arrows), and m1 became electron-transparent through the internal (large and small red arrows) and external assembly of dense particles. (**D4-D8**)

Aggregation of dense particles enlarged fm1, causing it to separate (fm1/1-fm1/3) and disperse into the particles (fm1/1 and double red arrowheads); fm2 lost its LMM while replenishing dense particles for DE1 via the aggregation of dense particles at the periphery (double red arrowheads), and its electron-lucent part was further divided by their internal assembly (red arrow and small black arrowheads). Condensed fm3 was diffused into nuclei (double red arrows), fm2 and DE1, which accelerated nuclear transition with the supplemented dense particles from both mitochondria (fm2 and fm3). At the edge of N2, the aggregation of particles from more than one mitochondria formed DEs (DE2 and DE3), concurrently creating electron-lucent DIs (DI1 and DI2), and their internal assembly further divided DI2 (DI2/1-DI2/4). Either MN2 or MN3 looked like a mitochondrion, within which internal and external assembly of dense particles occurred. The aggregation of the particles at mitochondrial peripheries formed dark strands (red arrows), and two strands from neighboring organelles combined to shape nuclear tubules (white arrowheads). At the cytoplasmic edge of MN3, fragmented mitochondria either directly dispersed into it (fm4, opposite red and white arrowheads) or aggregated dense particles to it (DI3 and black arrowheads). The internal assembly of dense particles in the organelle (DI3) formed a dark strand (large red arrow), which combined with aggregating dense particles at the periphery (small red arrow) of its neighboring counterpart in N2 to shape the tubule (small white arrow). Their further assembly caused dark strands and electron-transparent lumens to disappear.

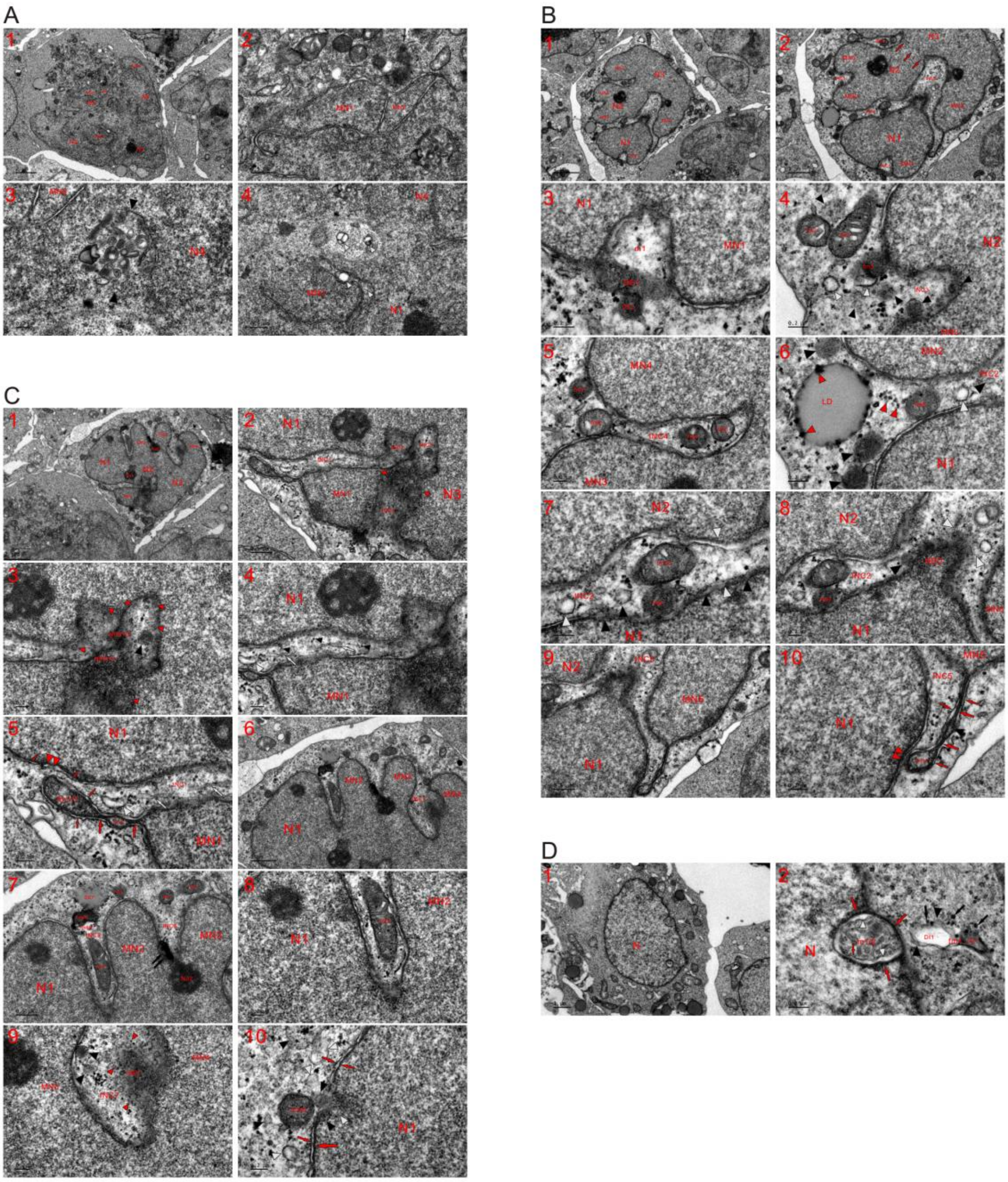


**Fig. S11. Incomplete nuclear transition caused mitochondria to become nuclear localized**. TEM was performed on four K562 cells at the 4 (**A-C**) and 8 (**D**) h time points, and the micrographs reveal that the nuclear localization of mitochondria and the formation of an INC was due to the incomplete nuclear transition of the organelles. (**A**) Nuclear formation and fusion occurred concomitantly (N1-N4 and MN1-MN4), and within nuclei or among them, partially fragmented mitochondria existed individually (white arrowheads) or as a group (opposite black arrowheads). (**B**) The combination of separately built nuclei (N1-N3 and MN1-MN5) partitioned the cytoplasm to form INCs (INC1-INC5) (**B1** and **B2**). (**B3**) Internal and external assembly of dense particles caused m1 to become electron lucent, disperse into nuclei (N1 and MN1) and look like a small and shallow INC at the nuclear edge (INC1). The simultaneous assembly of the particles with the mitochondria neighboring m1 in the cytoplasm formed electron-dense bodies (DE1 and DE2). (**B4**) A dispersed mitochondrion formed INC3 (black arrowheads), and the condensed organelles (fm1-fm3 and black arrowheads) were diffused through further aggregation of the particles. The assembly of the particles to the peripheries led to the formation of SDIBs (white arrowheads). (**B5**) Internal assembly of dense particles in large mitochondria formed small dense bodies, which looked like small mitochondria and dispersed into nuclei via further aggregation of the particles. Condensed mitochondria appeared in the cytoplasm outside of the nucleus (fm6 and fm7) as well as in the partitioned cytoplasm (INC4) (fm4 and fm5). (**B6-B8**) Formation of an NBD by fragmented mitochondria (black and white arrowheads) or mitochondrial aggregation of dense particles linked N1 and N2, concurrently three-sided partitioning the cytoplasm to form INC2. From the outside and inside of INC2, mitochondria fragmented to seal it and make it disappear (fm8-fm10, black, white and small red arrowheads). Mitochondrial aggregation of dense particles led to the formation of a lipid droplet (LD), which was assembled into the particles (small red arrowheads). Through their further assembly, condensed mitochondria (such as fm9 and fm10) and aggregates of dense particles (black arrowheads) provided the dense particles to intermix with the electron-lucent cytoplasm for nuclear development (opposite white arrowheads). (**B9** and **B10**) Internal aggregation and compaction of dense particles in the organelle formed dense bodies (fm11 and black arrowhead) and an elongated dark strand (large red arrows), which linked MN5 and N1 through fm8 and closed INC5. Adjacent to the strand, the congregation and linearization of the particles from its own and neighboring fragmented mitochondria shaped dark threads (small red arrows). The further assembly of dense particles caused fm11 to disperse into N1 (double red arrowheads) and the surroundings, and dense bodies, dark strands and threads disappeared by turning into the particles via a similar assembly mechanism. (**C**) Nuclear growth by mitochondrial aggregation of dense particles formed large (N1-N3) and small nuclei (MN1-MN4), and incomplete nuclear transition of the dispersed mitochondria and the internal and external assembly of dense particles, which reassembled into nucleus, led to the partitioning of the cytoplasm and to the simultaneous formation of INCs. Newly formed nuclei (NPB1/1, NPB1/2 and opposite red arrowheads) further divided the partitioned cytoplasm (INC1-INC4), and NPB1/2 was formed by the assembly of dense particles to the peripheries from neighboring mitochondria. Incomplete nuclear transition of the organelles or one completely fragmented mitochondrion shaped these small INCs (INC2-INC4), within which traces of mitochondrial fragmentations were observed (black and small red arrowheads). In narrow INC1 and before totally turning into dense particles (small black arrowheads), mitochondrial fissions led to the transient formation of ERLs (white arrows) and SDIB (white arrowheads) (**C1-C4**). (**C5**) Internal aggregation of dense particles condensed the organelle (such as fm1/1 and fm1/2) and formed dark strands to link MN1 with N1 while concurrently sealing INC1, and fm1/1 further assembled dense particles to diffuse into N1 (double red arrowheads) and the surroundings. The congregation and linearization of the particles at its periphery shaped dark threads (small red arrows); this also happened to the organelles at the edge of N1. (**C6-C9**) Formation of MNs (MN2-MN4) compartmentalized the cytoplasm to form INCs (INC5-INC7), and the assembly of dense particles in the organelles led to the mitochondrial condensations (fm2-fm5), separation of fm6 (fm6/1 and fm6/2), formation of DE7 and compaction into an ink mass (double black arrows). Through the further assembly of dense particles, these mitochondria-derived dense structures eventually turned into particles, and during the dispersion of electron-lucent fm6/2, its dense particles appeared in the cytoplasm. Nu2 was enlarged by dispersing of the ink mass into dense particles. Within INC7, an NBD was formed by mitochondrial assembling of dense particles and looked like a dispersed condensed mitochondrion, and fragmented mitochondria (black arrowheads) aggregated dense particles to it (small red arrowheads). (**C10**) During its fission through the aggregation of dense particles, part of fm8 enclosed into N1 (black and white arrowheads), and condensed fm8/2 was diffused via further assembly of the particles. Their assembly at the peripheries of neighboring mitochondria (or at periphery of a mitochondrion) formed a dark strand (large red arrow) and thread (small red arrow), both of which combined to form a tubule-NE fragment (opposite red arrows). (**D**) Aggregation of dense particles to the peripheries of m1 and its neighboring mitochondria formed dark strands (red arrows), which in turn enclosed the mitochondrion to become nuclear localized. Their internal assembly formed m1/2, which looked like an MVB. Between internal (m1/2) and external (red arrows) aggregation of dense particles, electron-lucent intervals (white arrowheads) created, and further congregation and linearization of dense particles at the edge of m1/2 shaped dark thread (small red arrow). In the cytoplasm and adjacent to m1, the aggregation of dense particles separated fm1 (such as DI1, DE1 and black arrowheads), and mitochondria-derived particles formed large glycogen-like granules (black arrows).

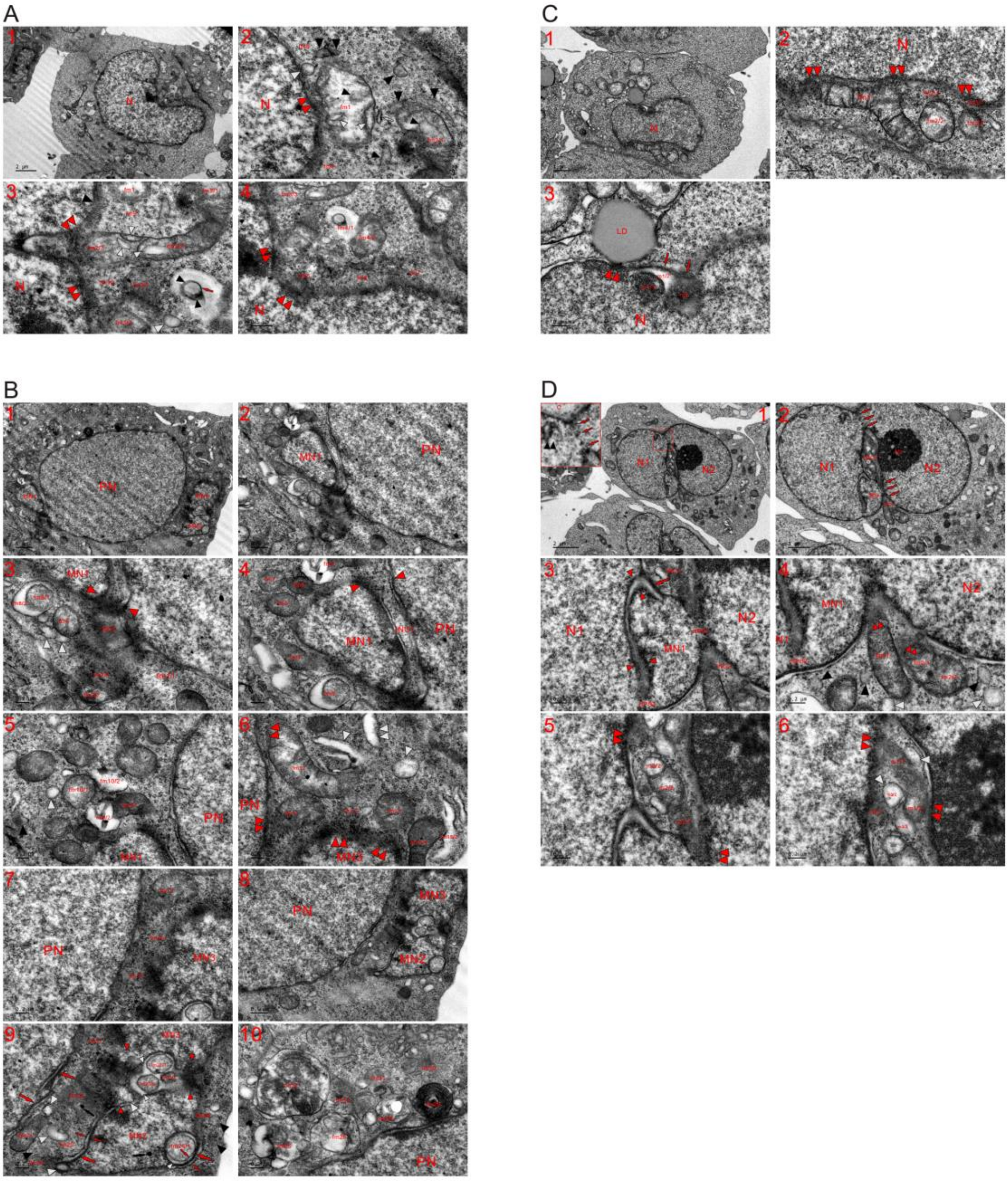


**Fig. S12. Fragmented mitochondria are directly dispersed into the nucleus to promote nuclear growth and fusion.** TEM was performed on four K562 cells at the 2 (**A** and **B**), 4 (**C**) and 8 (**D**) h time points, and the micrographs demonstrated that cytoplasmic mitochondria and the enclosed mitochondria dispersed into the nucleus to achieve nuclear transition through complete fragmentation into dense particles. (**A**) At the nuclear edge, mitochondria partially (fm1-fm4) or completely (fm5-fm10) fragmented by assembling dense particles that dispersed into the nucleus (double red arrowheads). Through the internal aggregation of the particles, a dark thread (red arrow), SDBs (small black arrowheads) and SDIBs (small white arrowheads) appeared within the organelles. During mitochondrial fissions (such as fm2/1-fm2/3; fm3/1 and fm3/2 and fm4/1 and fm4/2), the ERL (white arrow), SDBs (large black arrowheads) and SDIBs (large white arrowheads) transiently formed in the cytoplasm before eventually turning into dense particles, which reassembled and rearranged to achieve nuclear transition (**A1-A4**). (**B**) In addition to the PN, three MNs were separately formed in this cell, and mitochondria fragmented into dense particles to join together MNs (MN1-MN3) and the PN. Due to the enrichment of dense particles at each end of the nuclear interval, one or two NPBs (opposite red arrowheads) could form to achieve partial fusion between the PN and MN1 while concurrently partitioning the cytoplasm to form an INC (INC1). fm1 and part of fm2 (fm2/1) dispersed into dense particles, and their internal aggregation led to mitochondrial condensations (such as fm4-fm7), separated the organelles (fm8/1-fm10/1 and fm8/2-fm10/2) and caused the formation of SMLBs (black and white arrowheads) (**B1-B5**). (**B6-B9**) Mitochondria fragmented to diffuse into each other (such as fm11-fm15) as well as into the nucleus (double red arrowheads), and during mitochondrial fissions, spherical (white arrowhead) and rod-shaped SDIBs (fm15/2 and double white arrowheads) were formed. Between the PN and MNs (MN2 and MN3), condensed mitochondria (fm11 and fm21) further assembled dense particles to disperse, resulting in the appearance of completely fragmented mitochondria (fm16-fm18). In addition to the mitochondrial fragmentations, dense particles assembled to transiently form dark strands (large red arrows), threads (small red arrows), tubule-NE fragment (opposite small red arrows), ink dots (black arrows), SDIBs (white arrowheads) and SDBs (black arrowheads) (such as fm20 and fm22). Between MN2 and MN3, dense particles were internally assembled to form fm23 (fm23/1-fm23/3 and small white arrowheads), which cooperated with neighboring organelles (opposite red arrowheads) to merge two MNs following mitochondria-to-nucleus transition. At edge of MN2, internal (fm24/1 and small white arrowhead) and external (large red arrow) aggregation of the particles happened in fm24. The simultaneous aggregation of dense particles at mitochondrial peripheries (external assembly) from fm24 and the neighboring mitochondrion first formed a thick dark strand (large red arrow), while their external assembly led to the formation of dark threads (small red arrows) at edges of both fm24/1 and the fragmented organelle. (**B10**) At the nuclear edge (PN), the internal aggregation of dense particles caused organelles to temporally become MVBs (such as fm25-fm27) or a Nu-like structure (fm28). Once they were heavily or completely fragmented, mitochondria dispersed in the cytoplasm (such as fm29-fm32). (**C**) At the nuclear edge, mitochondria (such as fm1-fm3) dispersed in the form of dense particles into the nucleus (double red arrowheads) and the surroundings, and parts of these organelles were becoming (such as fm2/1) or had become (fm3/1) nuclear appearance. The aggregation of dense particles within the LD allowed it to fuse with the nucleus (double red arrowheads), and the assembly of the particles caused mitochondrial separation (m1/1 and m1/2) or fragmentation (m2) and included the organelles themselves (m1 and m2) in the nucleus (red arrows). (**D**) The same cell shown in **Figure 1E**. Traces of partial nuclear fusion between N1 and N2 are demonstrated (three red arrows and high magnification image of the inset in **D1**), and the nuclear fusion partitioned the cytoplasm to form the intranuclear inclusion. From both sides, mitochondria-derived dense particles assembled into NPB1, which linked nuclei (N1 and N2) and further divided the partitioned cytoplasm to shape the closed INC1 and small INC2. Due to the enrichment of dense particles at the cytoplasmic side of nuclei (MN1 and N1), NPB2 first formed by fragmented mitochondria to link the separately built MN1 with N1, and a dark strand and aggregates of the particles further assembled to complete nuclear merging (opposite red arrowheads) (**D1-D4**). (**D5** and **D6**) Both cytoplasmic mitochondria and enclosed mitochondria aggregated dense particles to disperse into nuclei (double red arrowheads) and the surroundings, and mitochondrial separations (such as m1/1 and m1/2; m2/1-m2/3; m3/1 and m3/2) and SMLB formation (black and white arrowheads) occurred concurrently.

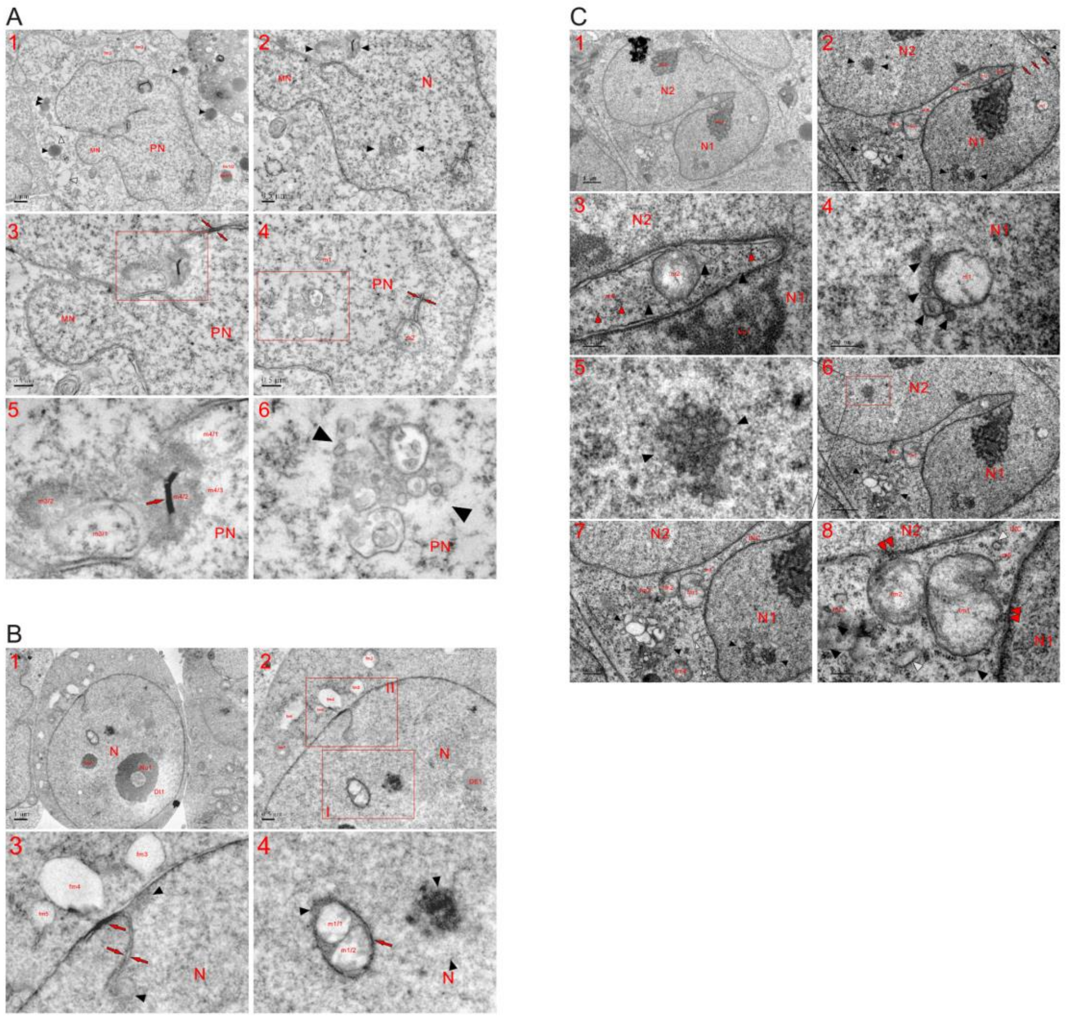


**Fig. S13. Nuclear-localized mitochondria were observed following the prolonged incubation of K562 cells.** TEM was performed on three K562 cells at the 48 h time point, and the micrographs reveal that mitochondria without enclosed NE appeared in the nucleus, and the occurrence of nuclear mitochondria was merely due to incomplete conversion into the nucleus of the organelles. (**A**) In the cytoplasm, the aggregation of dense particles separated mitochondria while concurrently forming electron-dense (fm1/1 and black arrowheads) and electron-lucent bodies (fm1/2 and white arrowheads). The assembly of dense particles to the periphery caused the organelles to become electron transparent (such as fm2 and fm3). Separately formed MN joined together with the PN, and external and internal aggregation of dense particles happened in both m1 and m2. Within the PN, mitochondrial separations (m3/1 and m3/2; m4/1-m4/3 and red arrow) and a group of fragmented mitochondria (opposite black arrowheads) were observed. Opposite red arrows: nuclear tubules (**A1-A6**). (**A5** and **A6**): high magnification images of insets in (**A3**) and (**A4**), respectively. (**B**) Aggregation of dense particles formed nucleoli (Nu1 and Nu2) concomitantly creating DI1 and condensed fm1, which was dispersed by further particle assembly. Their assembly at the periphery caused the organelles to become electron-lucent DIs (such as fm2-fm6), and both the internal (m1/1 and m1/2) and external (red arrow and black arrowhead) aggregation of dense particles occurred in nuclear-localized m1 (m1/1 and m1/2). In other parts of the nucleus, mitochondrial aggregation of the particles was observed (DE1; red and opposite red arrows; black and opposite black arrowheads) representing incomplete mitochondria-to-nucleus transition (**B1-B4**). (**B3** and **B4**): high magnification images of insets I and II in (**B2**), respectively. (**C**) Partial nuclear fusion was achieved between N1 and N2, which concomitantly compartmentalized the cytoplasm to form a large and narrow INC. m1, which did not contain an enclosed tubule-NE, became electron-transparent via the assembly of dense particles at the periphery. Adjacent to m1, fragmented mitochondria appeared (black arrowheads). Within the INC (m2) and at its opening (fm1 and fm2), less fragmented mitochondria were observed. Heavily or completely fragmented mitochondria (such as m2-m5; fm3 and fm4) formed SDBs (black arrowheads) and SDIB (white arrowhead) or totally turned into dense particles (small red arrowheads) within the INC and in the cytoplasm. In both the nuclei and cytoplasm, a group of fragmented mitochondria were observed (opposite black arrowheads). The aggregation of dense particles allowed the organelles to disperse into each other (fm1-fm3, black and white arrowheads) and nuclei (double red arrowheads) (**C1-C8**). (**C5**): a high magnification image of the inset in (**C6**).

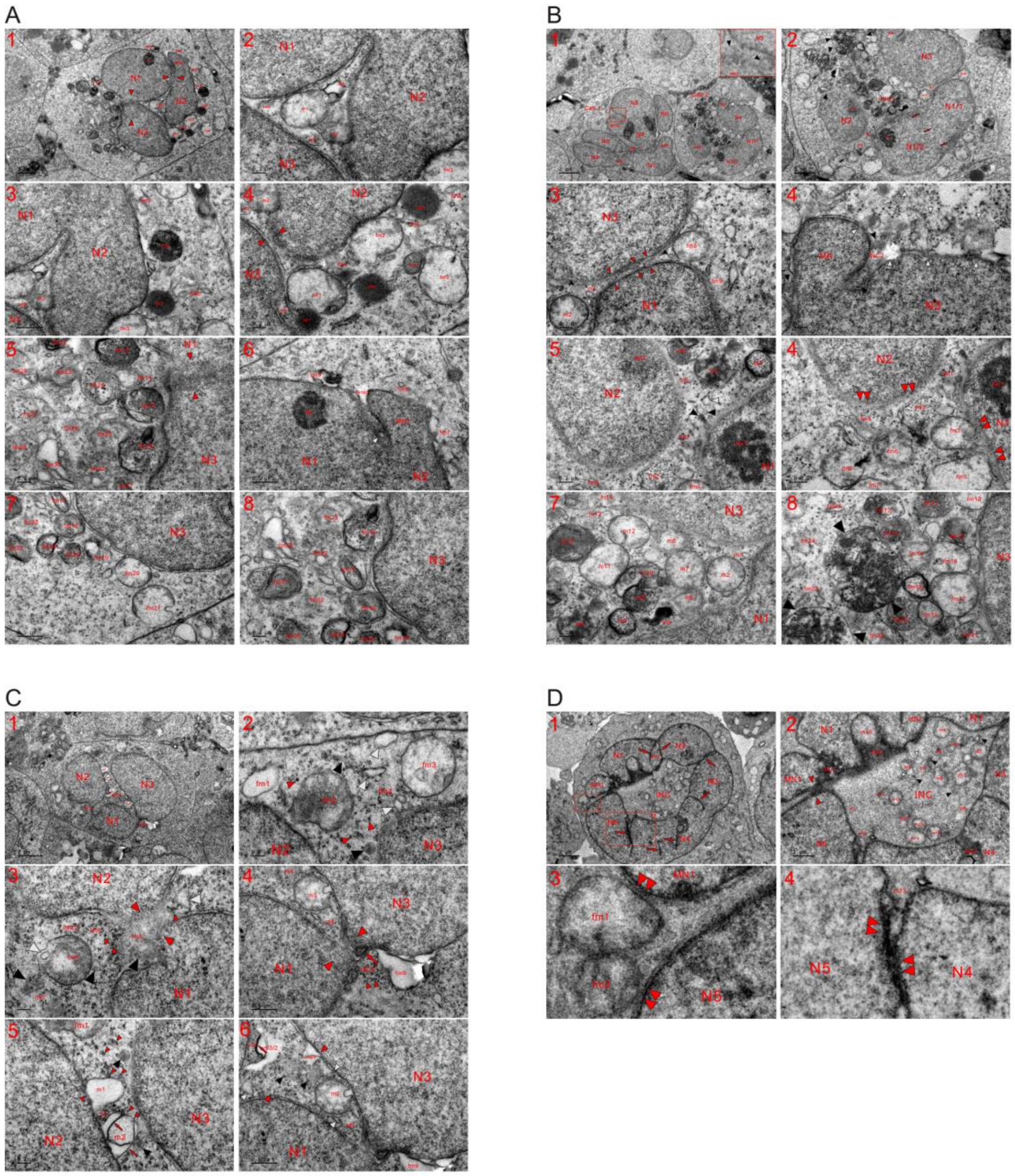


**Fig. S14. Joining together of separately formed nuclei partitioned the cytoplasm to form INCs at the 48 h time point.** TEM was performed on four K562 cells at the 48 h time point, and the micrographs demonstrate that more than three nuclei simultaneously and individually formed in a single cell, and nuclear fusions mediated by fragmented mitochondria partitioned the cytoplasm to form a large INC. (**A**) The same cell shown in **Figure 2A.** Micrographs of whole cells reveal that separately formed nuclei (N1-N3) were joined together (opposite red arrowheads), which concurrently partitioned the cytoplasm to shape an INC. Through the assembly of dense particles, the enclosed organelles (m1-m5) underwent mitochondrial fissions and fragmentation to disperse into each other and nuclei (**A1** and **A2**). (**A3** and **A4**) Mitochondrial-derived dense particles (m2, m3, fm1 and opposite red arrowheads) reassembled to seal the gap between nuclei (N2 and N3), and m6 further assembled the particles to diffuse in N3. The aggregation of dense particles caused the organelles to become electron lucent (such as fm1-fm3), caused the dispersions of the organelles (fm4-fm7), formed electron-dense bodies (DE1-DE4) and blackened the organelle to look like a Nu (DE5). (**A5-A8**) The gap between N1 and N3 was recently sealed (opposite red arrowheads), and completely fragmented mitochondria lost morphology (such as fm7 and fm8). fm9 was separated and dispersed via internal and external assembly of dense particles. External aggregation of the particles at the periphery allowed part of fm10 to become electron lucent (fm10/2) and fragmented mitochondria (fm10 and white arrowhead) to merge nuclei (MN1 and N1). In the cytoplasm, mitochondria internally and externally assembled dense particles to diffuse into each other and nuclei (fm12-fm35), and fm21 was dispersed into the cytoplasm via the assembly of the particles at its edge. The internal aggregation of dense particles formed glycogen-like granules in fm16, which dispersed into N3 via external assembly. fm31 internally assembled dense particles to form a structure of membrane whorls. (**B**) In cell No. 1 (Cell 1), several nuclei were merged to build a large one (N1-N6, MN1 and MN2) and the formation of NPBs (NPB1 and NPB2) linked separately formed nuclei (N1 and N2; N3 and N4). High magnification image of the inset reveals that mitochondria fragmented (black arrowheads) to join together MN2 and N5 (**B1**). In cell No. 2 (Cell 2), three-sided nuclear formations (N1-N3) partitioned cytoplasm to form a large and opened INC (INC1), small nuclei (N1/1 and N1/2) merged to build N1 (three abreast red arrows), and the interval between N1 and N3 was almost sealed (opposite red arrowheads). At the nuclear edge, partially (fm2 and m8) and heavily (such as m1, black and white arrowheads; fm9 and opposite white arrowheads) fragmented mitochondria were observed, and the morphology of the completely fragmented mitochondria was lost as they turned into dense particles. Within INC1 and at its opening, mitochondria fragmented into each other or grouped together (opposite black arrowheads) to promote nuclear development. Between MN and N3, INC2 looked like a dispersed mitochondrion (black and white arrowheads) (**B1-B4**). (**B5** and **B6**) Between N1 and N3, mitochondria fragmented (such as fm1-fm6, black and white arrowheads) to join together two nuclei, and the aggregation of dense particles caused the organelles to disperse into nuclei (double red arrowheads) as well as into each other (fm1-fm8). (**B7** and **B8**) Through the internal and external assembly of dense particles, cytoplasmic mitochondria that localized at the opening of INC1 (fm10-fm25) and the enclosed mitochondria (such as m1-m15) fragmented to diffuse into each other, and some of the organelles directly dispersed into nuclei. By grouping together (opposite black arrowheads), fragmented mitochondria tended to change into an MN. (**C**) Joining together of separately formed nuclei (N1-N3) compartmentalized the cytoplasm to shape INCs (INC1-INC3), and at the opening of INC1, mitochondria were partially (such as fm1-fm3), heavily or completely (fm4; black, white and small arrowheads) fragmented. To promote nuclear development, the organelles first turned into dense particles (small arrowheads) (**C1** and **C2**). (**C3** and **C4**) Nuclear fusion (opposite red arrowheads) between N1 and N2 occurred via fragmented mitochondria (fm5; black, white and small red arrowheads), and partial nuclear merging was already achieved between N1 and N3 (opposite red arrowheads) with trace mitochondrial fission (red arrow). In addition to nuclear fusions, small and shallow INCs (INC2 and INC3) were formed by partitioning the cytoplasm, and the INCs disappeared, while fragmented mitochondria (such as fm6-fm9; black, white and small red arrowheads) accomplished nuclear transition. (**C5** and **C6**) Within INC1, m1 became an electron-lucent or electron-dilute phase (DI) via the assembly of dense particles at its periphery (DE1 and small red arrowheads) and the internal (red arrow) and external (DE1, red arrow, black and small red arrowheads) aggregation of the particles occurred in m2. Their assembly separated the organelles (m3/1, m3/2 and red arrow; m4/2), leading m4/2 to fuse with N3 and causing mitochondria to be dispersed (m3-m6, black and white arrowheads). Heavily or completely fragmented mitochondria formed the SDB (black arrowhead) and turned into dense particles (small red arrowheads), which had changed (DE1 and m3/2) or were changing into a nuclear appearance (opposite red arrowheads). (**D**)

Individually formed small nuclei (N1-N5, MN1 and MN2) joined together to build a large ring-shaped nucleus while concurrently partitioning the cytoplasm to form an INC, which was closed by partially (fm1 and fm2) and completely (opposite red arrowheads) fragmented mitochondria. Traces of nuclear fusions are shown (red arrows), and m18 changed to a nuclear appearance through the internal and external aggregation of dense particles, and the particles from m18 and its neighboring counterparts formed DE1, which could develop into the appearance of Nu1. The organelles (fm1 and fm2) assembled dense particles to diffuse into each other and nuclei (double red arrowheads) before totally turning into particles. Partially (such as m1-m6 and m14-m17) and completely (such as m7-m13) fragmented mitochondria were present in the INC. During mitochondrial fragmentation and before dispersing into dense particles, SDBs (black arrowheads) and SDIBs (white arrowheads) were intermediately formed. Between N4 and N5, mitochondria assembled dense particles (such as m13 and double red arrowheads) to merge the nuclei (**D1-D4**). (**D3** and **D4**): high magnification images of inset I and II in (**D1**), respectively.

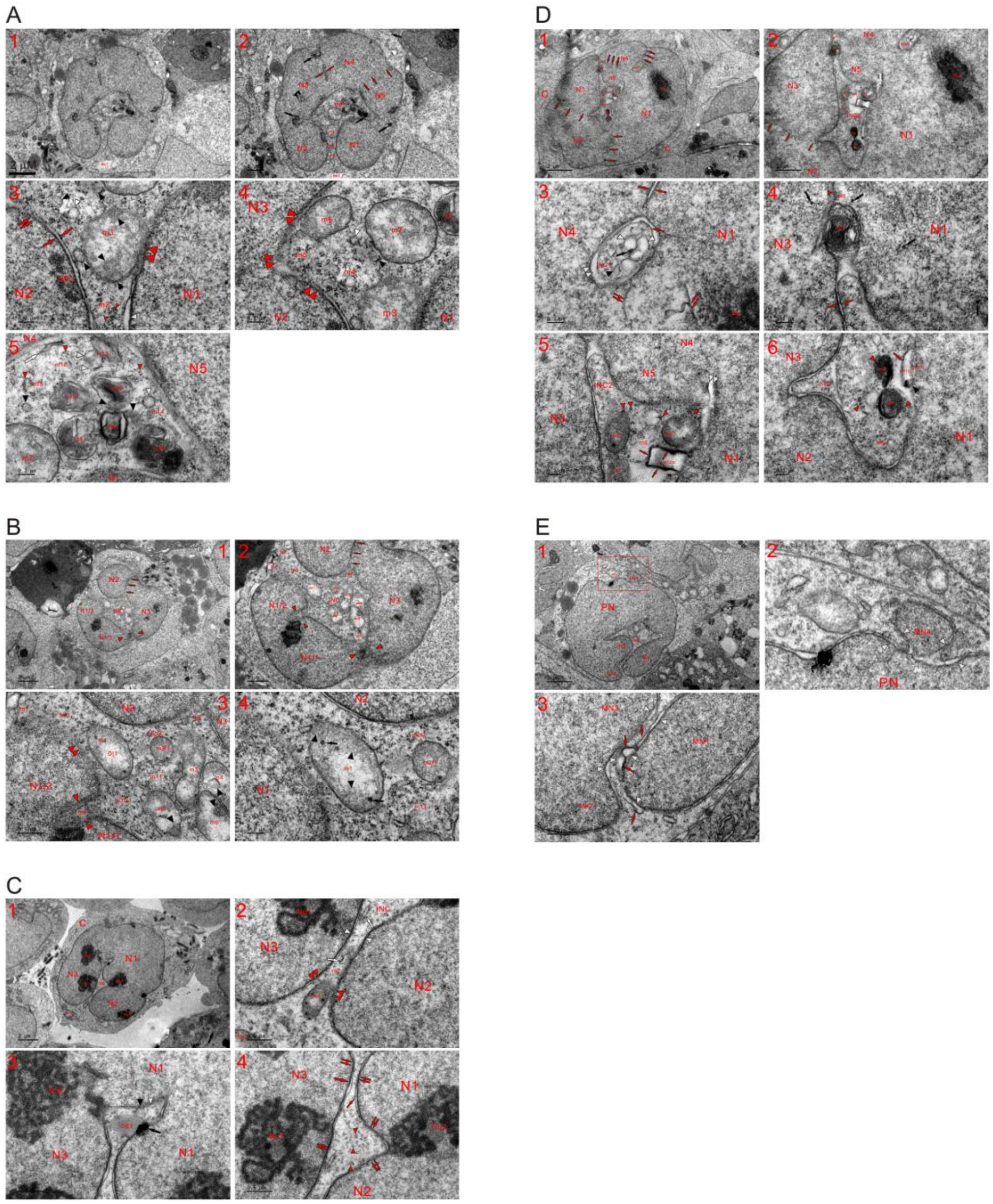

**Fig. S15. Mitochondria were present at the opening of a large INC and fragmented to close it.** TEM was performed on five K562 cells at the 48 h time point, and the micrographs show that mitochondria were present at the opening of a large INC and fragmented to seal it for nuclear development; due to the richness of dense particles at the opening, its closure usually occurred early than disappearance of the whole INC (**A-E**). (**A**) The same cell shown in **Figure 2B**, and traces of nuclear fusions representing the incomplete nuclear transition of mitochondria (black, three abreast and opposite red arrows; double black arrowheads). fm1 assembled dense particles to the opening, where completely (such as m1, m2, white and small red arrowheads) and partially (m3 and black arrowhead) fragmented mitochondria were present. m3 was dispersed (double red arrowheads) via the aggregation of dense particles (black arrowheads). The organelles (m4-m6) fragmented by assembling the particles to diffuse into nuclei (double red arrowheads) and the surroundings in the form of dense particles. The aggregation of the particles at periphery from neighboring mitochondria transiently formed tubule-NE fragments (opposite red arrows), which disappeared during further assembly. Internal and external aggregation of dense particles occurred in enclosed mitochondria (m7-m14), and completely fragmented mitochondria (such as m15 and m16) dispersed into the particles (small red arrowheads). During mitochondrial fissions, the ERL (white arrow), SDBs (black arrowheads) and SDIBs (white arrowheads) were intermediately formed (**A1-A5**). (**B**) Incomplete nuclear conversion of the organelles was observed (three abreast red arrows and opposite red arrowheads), and fragmented mitochondria (fm1-fm3 and m1) were present at the opening of the INC and dispersed into nuclei (double red arrowheads). Once completely fragmented, m1 diffused, resulting in the appearance of fm3. Part of m2 turned into dense particles (m2/2). Through the aggregation of dense particles (black arrows and arrowheads), the enclosed organelles (such as m2-m11) dispersed into each other and some of them diffused into nuclei. Due to the incomplete nuclear transition of mitochondria (such as m8 and m9), nuclear intervals appeared (between N1/1 and N1/2; between N2 and N3). As m8 changed into a nuclear appearance, m3 dispersed into m8 to provide dense particles for continuous nuclear growth and enlargement, and the external assembly of the particles at its periphery caused matrix of m1 become electron lucent (**B1-B4**). (**C**) Along with nuclear growth by mitochondrial assembling of dense particles, the partitioned cytoplasm (INC) narrowed and decreased, while partially fragmented mitochondria disappeared within the INC, in which only traces of mitochondrial fissions (DE1, black and white arrows, black and white arrowheads) were observed. The aggregation of the particles condensed m1, which thereafter further assembled dense particles (black arrows) to disperse into the nucleus (double red arrowheads), m2 and the surroundings. The aggregation of dense particles at the periphery caused the organelle become electron transparent and to lose its mitochondrial morphology (opposite white arrowheads). To make the INC disappear, former aggregates of dense particles, such as DE1, the ink mass (black arrow), the dark strand (large red arrow) and the thread (small red arrow), tended to disperse through their further assembly (double small red arrows). (**D**) Both INCs (INC1 and INC2) were sealed, and fragmented mitochondria (m1 and m2) were present at the closing site of INC2, which was larger and filled with fragmented organelles (such as m3-m11). Vestiges of nuclear fusions were observed (N1-N5 and three abreast red arrows) (**D1** and **D2**). (**D3**) INC1 was a group of fragmented mitochondria or one mitochondrion, in which internal and external assembly of dense particles occurred (red and black arrows; double white, black and white arrowheads). Either an SDB (black arrowhead) or SDIB (white arrowhead) could appear similar to a vesicle through the aggregation of dense particles at the periphery. Double white arrowheads: an electron-lucent interval, which formed by external and internal assembly of the particles; opposite red arrows: a nuclear tubule; double red arrows: undispersed aggregates of dense particles. (**D4**) External aggregation of dense particles at its periphery (small red arrowhead and black arrow) caused m1 to disperse into nuclei while concurrently forming electron-transparent spots (white arrowhead), and their internal assembly allowed m2 to become a structure of membrane whorls, which diffused into nuclei (N1 and N3) by the external aggregation of the particles. Black arrows: dark dots, which formed via the aggregation of mitochondria-derived dense particles; opposite red arrows: tubule-NE fragment, which was shaped via the external assembly of the particles from two neighboring mitochondria along their fragmentations. (**D5** and **D6**) Similar to m2, the organelles with membrane whorls (m3 and m4) were dispersed into dense particles (small red arrowheads), which were also derived from the condensed (m5-m7) and heavily or completely (such as m8-m10) fragmented mitochondria. Assembly of the particles separated m11 (m11/1 and m11/2; red arrow and black arrowhead) and allowed electron-opaque m11/1 to be dispersed. The aggregate of dense particles (red arrows), which surrounded the dilute m12/2 of three-sided, was diffused. Double white arrowhead: an electron-lucent spot, which formed via the external assembly of dense particles. (**E**) MN4 was merged with the PN by fragmented mitochondria (opposite white arrowheads), and the formation of MNs (MN1-MN3) reduced the partitioned cytoplasm (INC). Between MNs, the nuclear interval was sealed by dispersed mitochondria and dense particle reassembly (red arrows and white arrowheads).

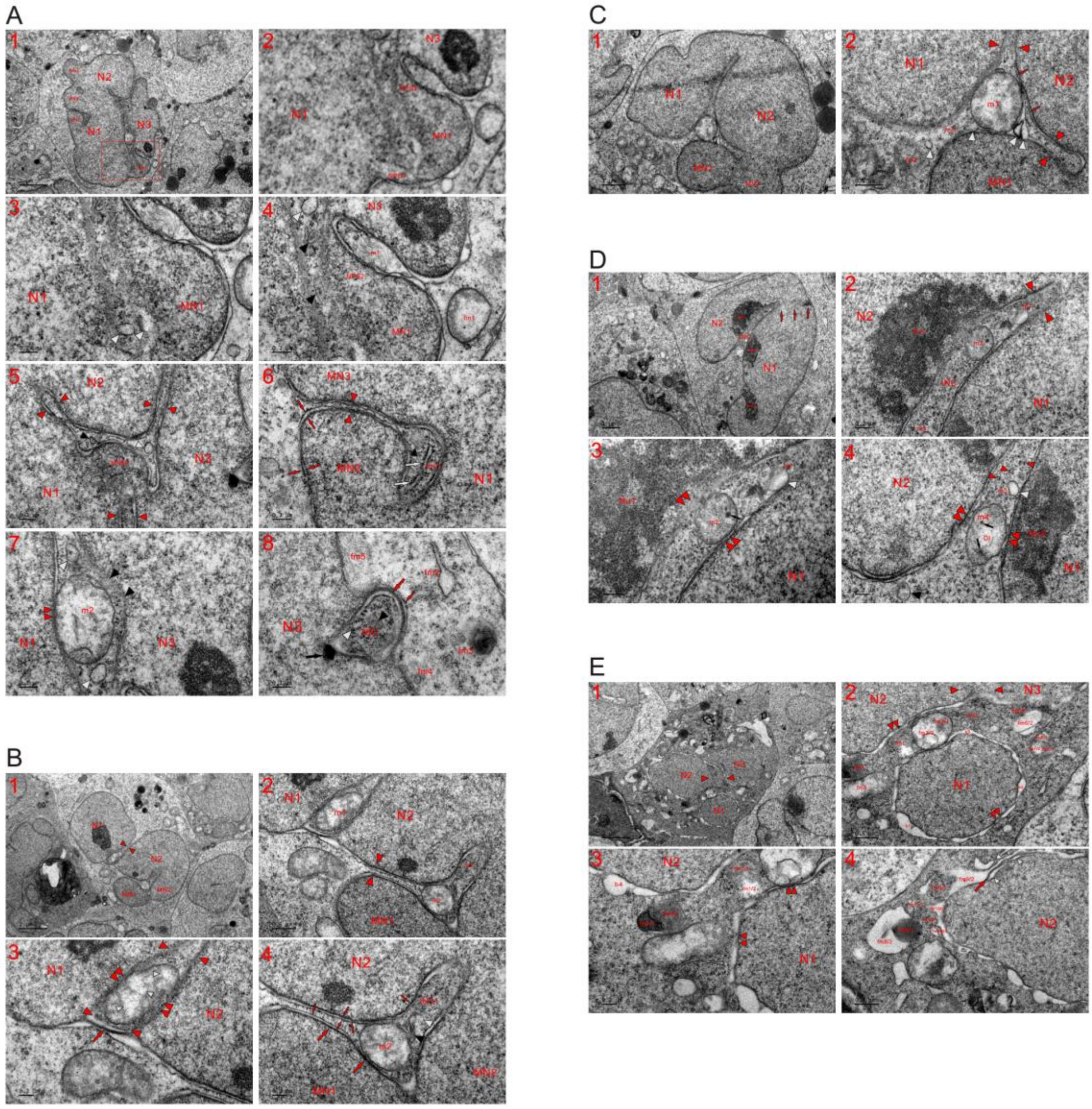


**Fig. S16. Mitochondria are included in the nucleus during the nuclear transition of neighboring counterparts.** TEM was performed on five K562 cells at the 48 h time point, and the micrographs show that a mitochondrion was included in the nucleus during the nuclear conversion of its neighboring counterparts. (**A**) Mitochondrial assembled dense particles (black and white arrowheads) to enlarge nuclear sizes (MN1 and N1) and consequently resulted in the combination of the separately formed MN1 and N1. The formation of NPBs (NPB1 and NPB2) first linked MN1 with the large nuclei (N1 and N3). The external aggregation of the particles dispersed m1 into nuclei (N3 and NPB2) and caused fm1 to become electron transparent, and the incomplete nuclear conversion of the organelle caused m1 look like the INC at the nuclear edge (**A1-A4**). (**A2**): a high magnification image of the inset in (**A1**). (**A5**) Separately formed nuclei (N1-N3) were merged together (opposite red arrowheads), and along with nuclear growth and NPB3 formation by mitochondrial fragmentation (black and white arrowheads), the space available in the partitioned cytoplasm reduced as less fragmented mitochondria disappeared. (**A6**) Along the growth of MN2 and MN3, the nuclear interval was almost sealed (opposite red arrowheads), and the space of the compartmentalized cytoplasm (INC1) decreased. Within INC1, the organelle fragmented to transiently form ERLs (white arrows) and an SDB (black arrowhead) before totally turning into dense particles. Opposite red arrows: tubule-NE fragment, which was shaped by the external assembly of the particles from neighboring mitochondria or by their internal assembly in a single mitochondrion. (**A7**) Internal and external aggregation of dense particles in m2 caused it to fuse with N1 (double red arrowheads) and become electron-lucent, and the mitochondria that neighbored it fragmented for nuclear development (black and white arrowheads). (**A8**) At the edge of N3, a mitochondrion had changed into a nuclear body (NB1) and included itself in N3 via the external aggregation of dense particles at its periphery (black and red arrows). Internal assembly also occurred (black and white arrowheads). Adjacent to N3, heavily or completely fragmented mitochondria (fm2-fm5) were observed, and the aggregation and linearization of the neighboring organelle formed a dark thread (small red arrow). (**B**) Partial nuclear fusion was achieved between MNs (MN1 and MN2), and pointed merging (opposite red arrowheads) occurred between large nuclei (N1 and N2) (opposite red arrowheads). During the nuclear transition of its neighboring counterparts via the assembly of dense particles (red arrow and opposite red arrowheads), m1 was included between or among nuclei and dispersed into them (double red arrowheads) via the assembly of particles, which led to the formation of lucent spots within the organelle (small white arrowheads). Partial merging was achieved between nuclei (MN1 and N2), which concurrently closed INC1, whose formation was due to the incomplete transition of mitochondria (such as m2, m3, black and white arrowheads) that existed in the partitioned cytoplasm (INC1). Fragmented mitochondria formed dark threads (small red arrows) via the further aggregation and linearization of dense particles and their external assembly at mitochondrial periphery shaped dark strands (large red arrows), which combined with the thread to create tubule-NE (opposite small red arrows) (**B1-B4**). (**C**) During nuclear enlargements (N1, N2, MN1 and MN2) caused by mitochondrial fissions and the consequent nuclear conversion (white and opposite red arrowheads; red arrows), both the partitioned cytoplasm and enclosed mitochondria decreased. Mitochondria fragmented (such as fm1, fm2 and white arrowhead) to accomplish partial nuclear fusion between nuclei (MN1 and N1), and once the neighboring organelles developed into a nuclear appearance, the less fragmented m1 became a nuclear mitochondrion as it still possessed the morphology at that time (**C1** and **C2**). (**D**) Partial merging (three abreast red arrows) between nuclei (N1 and N2) compartmentalized the cytoplasm to form an opened INC, which was reduced along with mitochondria-derived dense particles that reassembled for nuclear development (m1 and opposite red arrowheads). Within the INC, less fragmented mitochondria (m2 and m4) aggregated dense particles to disperse into nuclei (double red arrowheads) and the surroundings, and heavily or completely fragmented mitochondria had almost dispersed into dense particles (small red arrows). Before totally diffused into the cytoplasm, a SDB evolved into a vesicle through the assembly of dense particles at the periphery (black and white arrowheads) (**D1-D4**) (**E**) Three nuclei were formed in this cell (N1-N3), and partial nuclear fusion (opposite red arrowheads) was accomplished between two of them (N2 and N3). Mitochondria fragmented for nuclear development to join together N1 and N2 (fm1-fm5). Aggregation of dense particles separated the organelles into dense (fm1/1, fm2/1, fm6/1 and fm7/1) and electron-lucent parts (fm1/2, fm2/2, fm6/2 and fm7/2), and condensed fm4 further divided (fm4/1 and fm4/2). A dense spherical-shaped body (fm8/1), which looked like a lipid droplet, separated from lucent fm8/2, and either dense (such as fm8/1-fm11/1) or lucent (such as fm9/2, fm10/2 and fm10/3) parts of the organelles were connected. The connection of these electron-transparent structures formed the lunar halo. After occurring once at the nuclear edge, it looked like nuclear bubbles (such as b1-b4) following the aggregation of dense particles at the periphery. The particles from other mitochondria (such as fm1, fm2 and fm8) or from its own dense part reassembled into lucent structures (double red arrowheads). At the edge of N2, the internal assembly of dense particles in the organelle formed a dark strand (red arrow), and their internal and external aggregation created electron-lucent structures (white arrowheads).

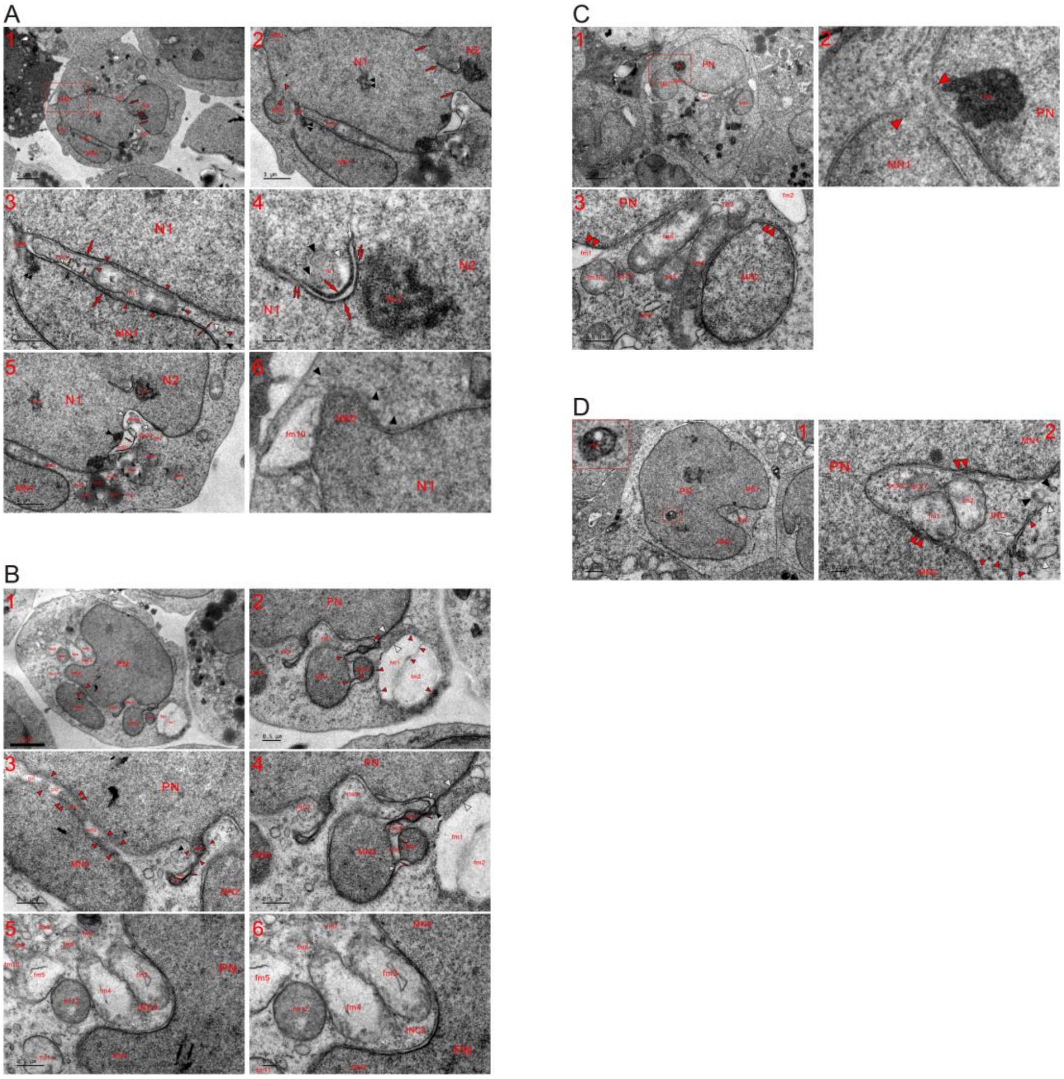


**Fig. S17. Attachment of an individually built MN to the PN partitioned the cytoplasm to shape the small and shallow INC.** TEM was performed on four K562 cells at the 48 h time point, and the micrographs demonstrate that either the attachment of an MN to the PN or the fusion of large nuclei partitioned the cytoplasm to shape a small and shallow INC at the nuclear edge. (**A**) The same cell shown in **Figure 2D**. MN2 was merged with N1 (opposite red arrowheads), which linked with MN1 through a newly formed NPB, and the attachment of MN1 and nuclear fusion between large nuclei (N1, N2 and three abreast red arrows) partitioned the cytoplasm to form INCs (INC1-INC3). The assembly of dense particles caused m1 to disperse (red arrowheads) and formed dark strands at the peripheries of fragmented mitochondria (large red arrows), and further aggregation and linearization of the particles in m2/1, which was an internal aggregate of m2, shaped dark threads (small red arrows). The combination of the strand and thread created a tubule-NE fragment (opposite small red arrows), and along with the aggregation of the particles, electron-lucent structures (white arrowheads) appeared. In INC1, both SDB (black arrowhead) and SDIB (large white arrowhead) were diffused into dense particles (small red arrowheads), and their internal assembly was observed in MN1 (double black arrowheads) (**A1**-**A3**). (**A4**) Adjacent to Nu1, internal (red arrow, black and white arrowheads) and external (red arrow) aggregation of dense particles occurred in m1, which was dispersed, and two dark strands combined to form nuclear tubules (white arrow). Through their further assembly, dark strands dispersed in the form of dense particles into Nu1 and the electron-lucent lumen of nuclear tubules (double small red arrows). (**A5**) At the opening of INC1, the congregation of dense particles separated (fm1/1 and fm1/2; fm2/1 and fm2/2; fm3/1 and fm3/2; fm5/2, red arrow and black arrowhead; fm6/1 and white arrowheads) and dispersed (fm1/2-fm3/2, fm4, fm7-fm9) mitochondria, and the incomplete nuclear transition of mitochondria (fm4-fm8) and nuclear fusion (N1 and N2) caused the partitioned cytoplasm to look like a small and shallow INC (INC3). (**A6**) A high magnification image of the inset in (**A1**) shows that fm10 became electron transparent via the assembly of dense particles at its periphery and cooperated with other fragmented mitochondria (black arrowheads) to form an NBD. (**B**) Aggregation of dense particles at mitochondrial peripheries (small red arrowheads) caused the organelles to become electron lucent (fm1 and fm2), the particles reassembled into nuclear pieces at the edge of the PN (opposite white arrowheads), and mitochondria fragmented to link nuclei (opposite red arrowheads). Between MN3 and PN, the assembly of dense particles in mitochondria (such as m1/1 and m1/2) dispersed the organelles to merge nuclei (double and opposite red arrowheads) concurrently forming electron-lucent spots (DI1 and DI2), and their aggregation (small red arrowheads) from the fragmented organelles (black and white arrowheads) formed a NBD at the edge of the PN. Their aggregation separated mitochondria (fm3/1, fm3/2 and red arrow; fm4/1, fm4/2, red arrow, black and white arrowhead; fm5/2, red arrow and white arrowhead) and caused the organelles to link with nuclei (fm3 and NBD; fm4, PN and MN2; fm5, MN1 and MN2), and the formation of INCs (INC1 and INC2) was due to mitochondrial aggregation of dense particles that led to nuclear formation (PN, MNs, NBD) and dark strand assembly. Along with the further assembly of dense particles, which existed in dark strands (red arrows) and condensed mitochondria (NBD, fm3/1, fm4/1 and black arrowhead), dilute nuclear intervals (between MN1 and MN2; between MN2 and NBD) and INCs (INC1 and INC2) eventually disappeared (**B1**-**B4**). (**B5** and **B6**) Three-sided nuclear growth (PN and MN4) or the attachment of MN4 to the PN (black arrows) compartmentalized the cytoplasm to form INC3, which was shallow and located at the nuclear edge. Within INC3 and adjacent it, mitochondrial fissions occurred through the aggregation of dense particles (fm3-fm11), and condensed fm12 tended to change into a small MN. (**C**) Individually formed MN1 was attached to the PN by fragmented mitochondria (opposite red arrowheads), the organelles (such as fm1-fm8) assembled dense particles to disperse into each other or/and nuclei (MN2 and PN), and the attachment of MN1 partitioned the cytoplasm to form an INC. The aggregation of dense particles at the periphery caused mitochondria (fm1 and fm2) to become electron transparent and to fuse with nuclei (double red arrowheads), and their dispersion into each other promoted nuclear development (**C1-C3**). (**D**) High magnification image of the inset reveals a dense nuclear body, which was derived from the internal assembly of dense particles in a mitochondrion (m1). Three-sided nuclear growth or the attachment of MNs (MN1 and MN2) to the PN partitioned the cytoplasm to form an opened INC at the nuclear edge, and within it, the aggregation of dense particles separated the organelles (fm3/1 and fm3/2) and dispersed them into each other or/and into the nucleus (double red arrowheads). At the edge of the INC on the cytoplasmic side, mitochondrial fissions transiently formed the ERL (white arrow), SDBs (black arrowheads) and SDIBs (white arrowheads), and all these structures eventually dispersed into dense particles (small red arrowheads) (**D1** and **D2**).

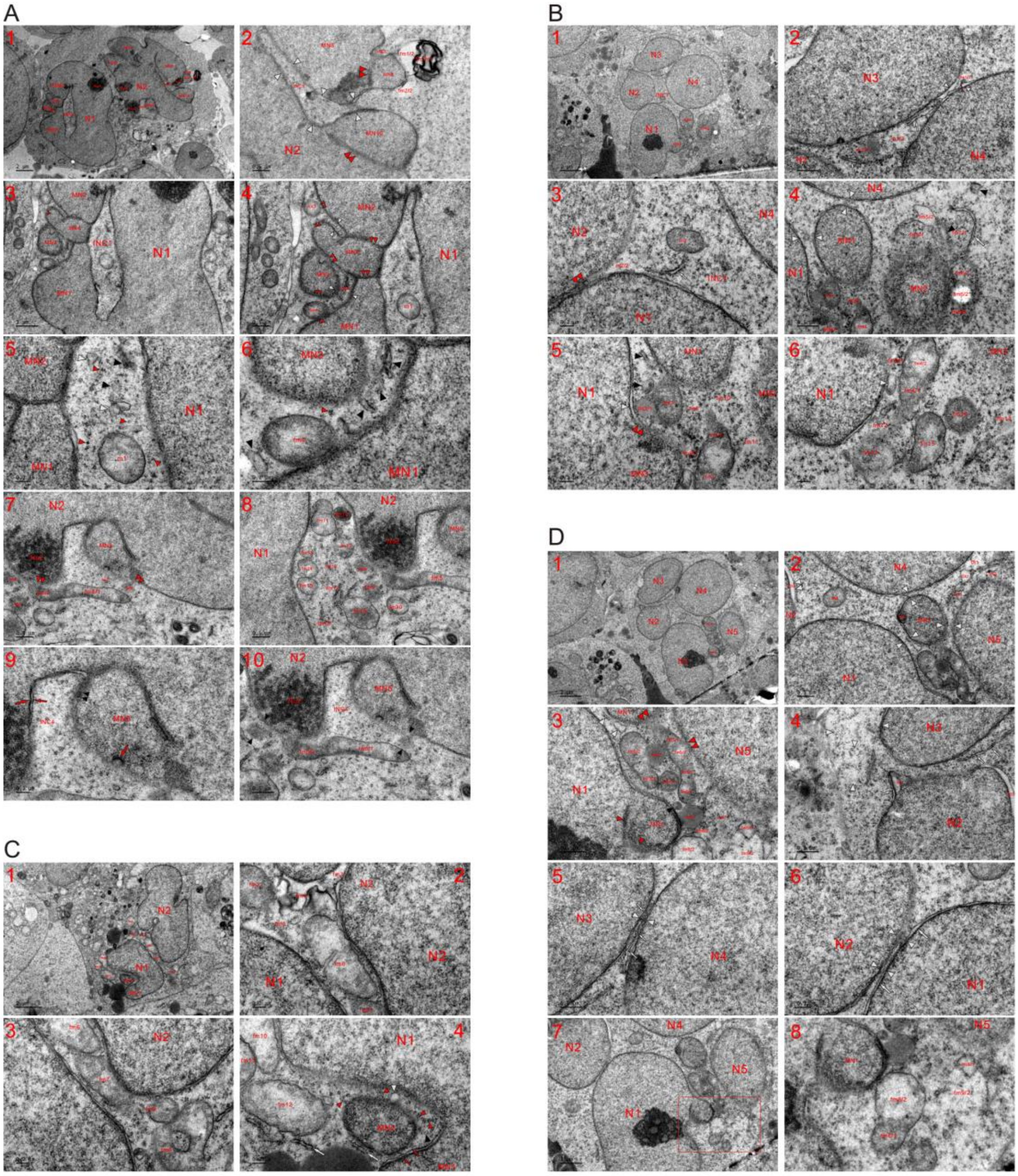


**Fig. S18. Individual formation of an MN and its attachment.** TEM was performed on four K562 cells at the 48 h time point, and the micrographs show that the attachment of individually formed MNs built the large nucleus and partitioned the cytoplasm to form INCs, which were large or small and closed or opened. (**A**) In addition to two large nuclei (N1 and N2), ten small nuclei appeared in the cell (MN1-MN10), and these MNs fused with each other or joined together with the large nuclei (**A1**). (**A2**) Nuclear merging (MN8 and MN9; MN10 and N2) was accomplished (double red arrowheads), and mitochondria-derived dense particles assembled for nuclear development to seal the intervals (INC3, INC5 and opposite white arrowheads). The internal and external aggregation of dense particles separated fm1 (fm1/1 and fm1/2) and allowed fm1/1 to become a structure of membrane whorls, and the external assembly of the particles at the periphery in fm1/2 expanded MN9 and promoted the disappearance of INC5 while concurrently changing it into an electron-lucent structure. A dense part of fm2 assembled into MN9, and fm2/2 became electron transparent via the external assembly of dense particles. (**A3-A6**) MN4 was merged with other MNs (MN1-MN3) while concomitantly partitioning the cytoplasm to shape either large and closed INC1 or small and shallow INC2, and within INCs, partially (m1) and heavily (black and white arrowheads) or completely (small red arrowheads) fragmented mitochondria were observed. The dispersion of condensed mitochondria (fm3 and fm4) into nuclei (double small red arrowheads) and the further assembly of the particles by fragmented mitochondria eventually caused dilute nuclear intervals (INC2 and opposite white arrowheads) to disappear and achieved the nuclear conversion of the organelles. (**A7-A10**) Mitochondria (such as fm5/2, fm6 and fm7) assembled dense particles to disperse into nuclei (double red arrowheads), and partial (such as fm10-15) and heavily or completely (such as fm8 and fm16-fm18) fragmented mitochondria were present between large nuclei. Their dispersion into each other promoted nuclear development, and the further assembly of dense particles from fm5 into dense structures (Nu2, large and small red arrows, and double black arrowheads) eventually caused INC4 to disappear. (**B**) Nuclei were joined together

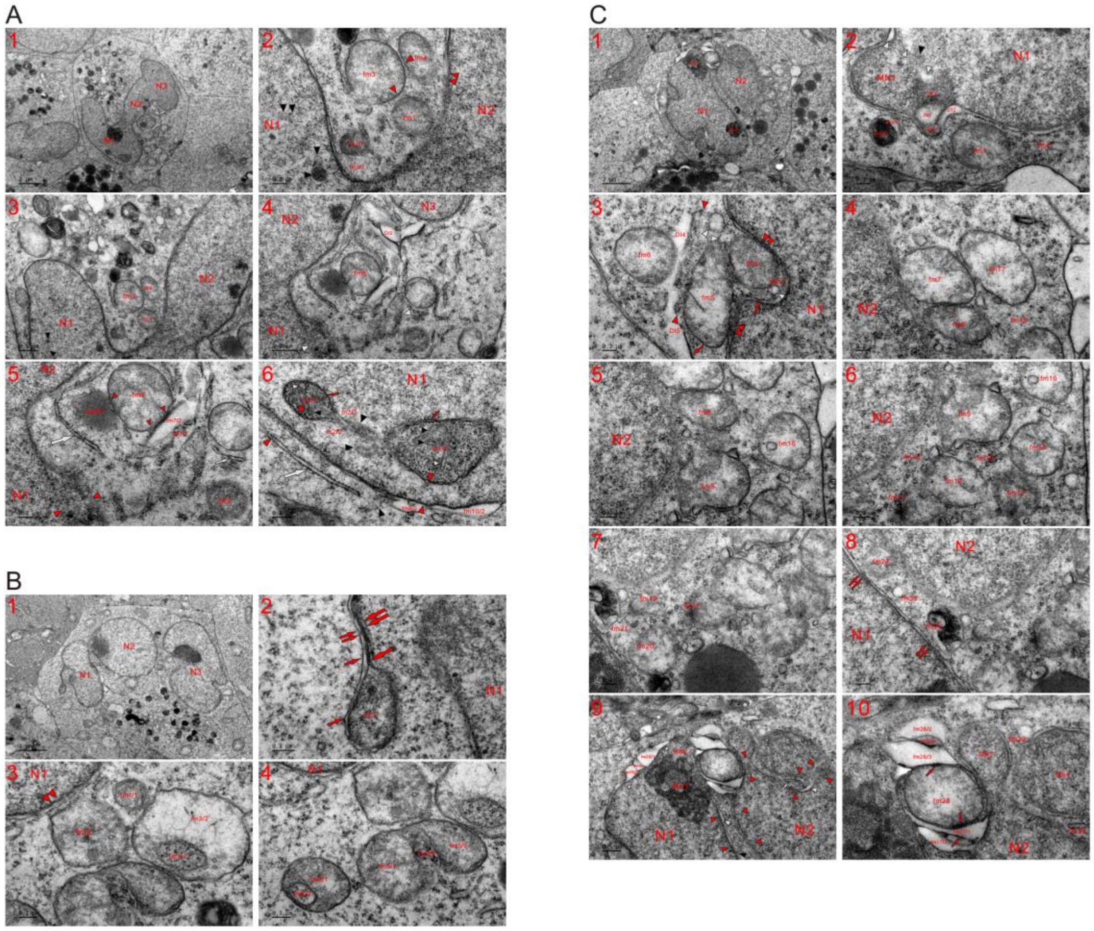


**Fig. S19. A whole mitochondrion developed into the appearance of a small MN through the assembly of dense particles.** TEM was performed on three K562 cells at the 48 h time point, and the micrographs reveal that one whole mitochondrion assembled dense particles resulting in the appearance of a small MN or with a similar density to that of nuclei (**A-C**). (**A**) Three separately formed nuclei (N1-N3) joined together, and within N1, aggregates of mitochondria-derived dense particles were observed (black arrowheads). Between N1 and N2, fm1 looked like a small MN and dispersed into N2. The aggregation of dense particles separated fm2 (fm2/1 and fm2/2) and caused the organelles to diffuse into each other (red arrowheads) or/and nuclei (fm2/2, fm4 and double red arrowheads) (**A1-A3**). (**A4** and **A5**) At the edge of N1, mitochondrial assembling of dense particles for nuclear development was observed (opposite white arrowheads). Adjacent to N2, the density of fm5 was similar to that of the nucleus. The internal aggregation of dense particles in the organelle formed a dense body (fm6/1), which looked like a LD, appeared in the cytoplasm along with dispersion of the whole mitochondrion and was diffused into either fm5 (small red arrowheads) or the surroundings. The internal aggregation of the particles separated fm7 (fm7/1 and fm7/2) and their external assembly caused dense particles to enter fm5 (small red arrowheads) while concurrently diluting fm7/2. fm8 lost the LMM, and the appearance of fm8 was similar to that of the nucleus at the edge of N1 (opposite red arrowheads). Based on mitochondrial fragmentation, the assembly of the particles at the periphery shaped ERLs (large and small white arrows). (**A6**) Internal aggregation of dense particles in mitochondria formed dense bodies (m1/1, m2/1 and black arrowheads) in N1, and further particle assembly occurred in both of them (red arrows, black and white arrowheads). The external assembly of particles from the organelles (m1 and m2/2) and neighboring counterparts (such as fm9/2, fm10/2, black arrowhead and large white arrow) achieved nuclear transition (opposite red arrowheads), during which the mitochondria became nuclear localized. The aggregation of dense particles at its periphery caused the separated organelles to become electron lucent and fuse with the nucleus to look like nuclear bubbles (fm9/2 and fm10/2). (**B**) Adjacent to N1, fm1 looked like a small MN and connected with N1 through a dark strand (large arrow) and thread (small arrow), both of which were dispersed (double large and small red arrows) via further assembly of dense particles. (**B1** and **B2**). (**B3** and **B4**) At the edge of N1, fm2 assembled dense particles to disperse into the nucleus (double red arrowhead) and the surroundings, and their internal assembly separated mitochondria (fm3/1 and fm3/2; fm5/1-fm5/3; fm6/1 and fm6/2). Once the whole fm3 diffused, fm3/1 appeared similar to fm4/1. (**C**) Mitochondria dispersed via the assembly of dense particles (such as DE1, DE2, DI1, DI2, black and white arrowheads) to promote nuclear growth while concurrently merging MN1 and N1. At the edge of N1, electron-opaque fm1 was dispersed into the appearance of fm2, and fm3 aggregated dense particles to separate itself (ink fm3/1 and lucent fm3/2) (**C1** and **C2**). (**C3**) While its neighboring counterparts accomplished the nuclear development, fm4 and part of a heavily fragmented mitochondria (red arrow and white arrowheads) formed a small and shallow INC (INC1), and mitochondria (fm4 and fm5) fused with the nucleus (double red arrowheads) and began to diffuse through the assembly of dense particles at its periphery (external assembly). During the aggregation of the particles (red arrow and opposite red arrowheads), connected electron-lucent structures (DI4 and DI5) appeared, whose formation promoted the nuclear development of the enclosed organelles (fm4, fm5, white and opposite red arrowheads). Through the external assembly of dense particles, fm6 was dispersed. (**C4-C7**) Aggregation of dense particles caused the organelles to diffuse into N2 and each other (such as fm7-fm21), and some of them had evolved into a nucleus-like appearance (such as fm8 and fm9). (**C8**) Between N1 and N2, fragmented mitochondria dispersed into N2 to promoted its expansion via the aggregation of the particles (such as fm22-fm24), and their further assembly caused tubule-NE to disappear (double red arrows). (**C9** and **C10**) External assembly of dense particles allowed mitochondria (fm25-fm27) to become electron transparent, their internal aggregation divided the organelles (fm25/1-fm25/3; fm26/1-fm6/3; fm27/1, fm27/2 and red arrow), and the aggregates of dense particles entered these lucent mitochondria through further assembly of the particles for nuclear development. During their further assembly, the enclosed fm28 could evolve into an MN. Nuclear formation (MN2 and opposite white arrowheads) by dense particles from the organelles (fm25-fm8) caused Nu1 to be away from the nuclear edge. When the nuclear fusions were almost finished (opposite red arrowheads), both MNs (MN3 and MN4) became attached, and traces of mitochondrial fragmentation were observed (fm29, fm30, white arrow, black and white arrowheads).

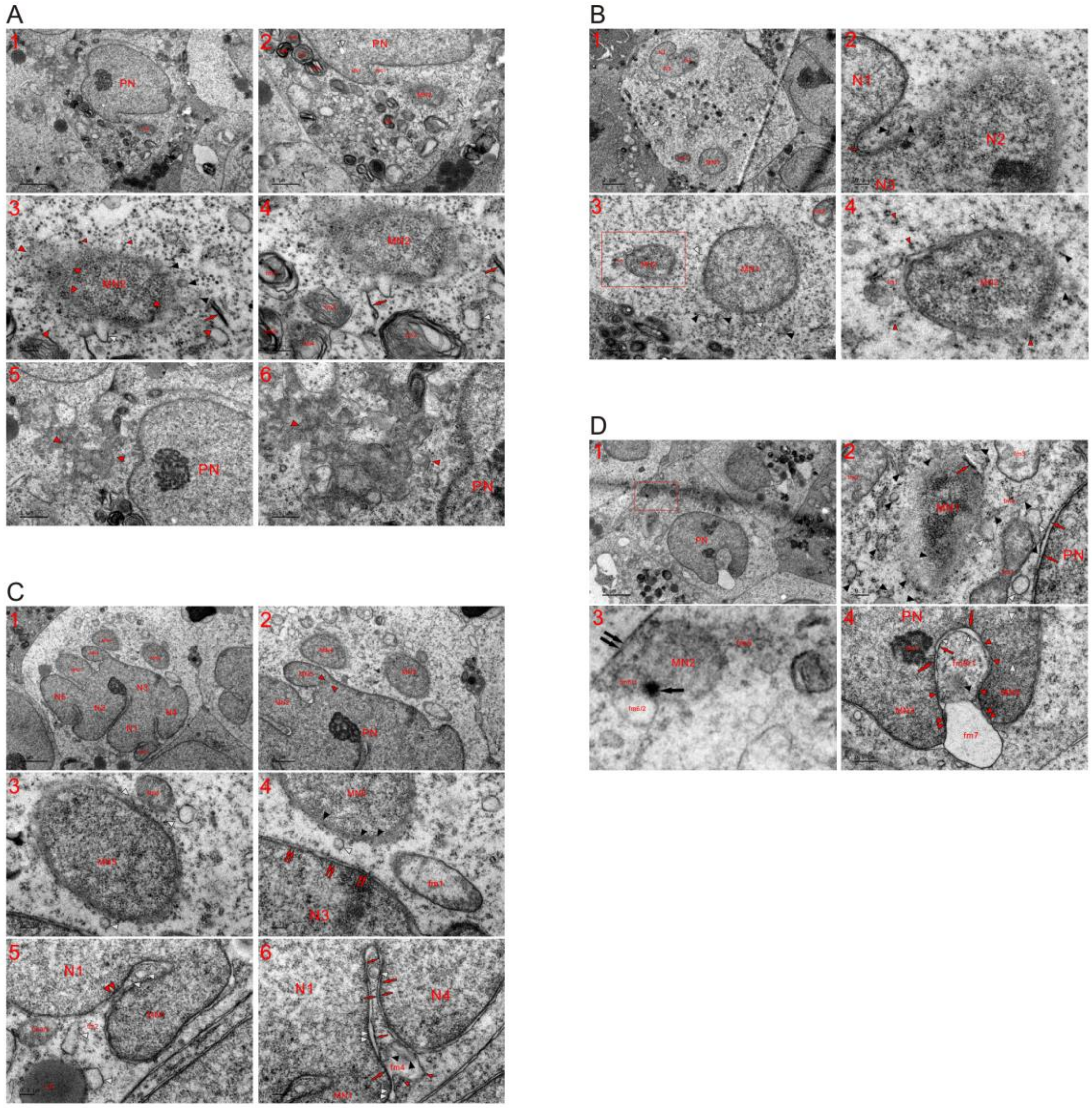


**Fig. S20. Formation of one or more MNs in a single cell that already possessed a PN.** TEM was performed on four K562 cells at the 48 h time point, and the micrographs show that one or more MNs separately formed in a single cell, which already possessed a PN. (**A**) MN1 had just attached to the PN as demonstrated by the vestiges of nuclear fusion (INC1 and double red arrowheads), and fragmented mitochondria (opposite red, black and white arrowheads) and mitochondria-derived dense particles (small red arrowheads) surrounded MN2 to promote its growth. At the edge of the PN and somewhat far away from MN2, the internal aggregation of dense particles caused the organelles to be structures of membrane whorl (such as fm1-fm9) , which dispersed to provide the particles (red arrows) (**A1-A4**). (**A5** and **A6**) A group of fragmented mitochondria appeared at the edge of the PN, and their grouping accelerated nuclear development (opposite red arrowheads). (**B**) A large nucleus was built (N1-N3), and a fragmented mitochondrion existed between nuclei (red arrow and black arrowheads). Adjacent to the plasma membrane, two small nuclei (MN1 and MN2) appeared, and all nearby mitochondria were heavily or completely fragmented (fm1, black and white arrowheads). After the dispersion of the organelles, they turned into dense particles (small red arrowheads), which reassembled into MNs for nuclear growth (**B1-B4**). (**B4**): a high magnification image of the inset in (**B3**). (**C**) In addition to the attached micronuclei (MN1-MN3 and opposite red arrowheads), three small nuclei were individually formed (MN4-MN6), and MN6, which was fused with MN5 by the fragmented organelle (opposite white arrowheads), looked like a dispersed mitochondrion. Through the further assembly of dense particles (double red arrows and black arrowheads) and the dispersion of the organelles (fm1 and white arrowhead), MN5 fused with N3 (**C1-C4**). (**C5** and **C6**) Aggregation of dense particles dispersed fm2 (white arrowhead), condensed fm3 (fm3/2), diluted fm4 (black and small red arrowheads) and led to the formation of a LD (white arrowhead). During particle assembly in the organelles, a tubule-NE fragments (opposite red arrows) and nuclear bubbles (double white arrowheads) appeared. Mitochondria-derived dense particles (fm2, fm4, double red and white arrowheads) in the combination with their further assembly in the aggregates (fm3/1, LD, small and large red arrows) eventually sealed the intervals between the nuclei (MN1 and N1; N1 and N4). (**D**) MN1 appeared to be at the initial stage of nuclear formation and looked like an aggregate of dense particles, which were derived from completely fragmented or dispersed mitochondria (red arrow, black and white arrowheads). Mitochondria that were somewhat far away from it were usually less fragmented (such as fm1-fm3). Their further assembly dispersed electron-opaque fm1 into either nucleus (MN1 and PN), allowing dark threads to disappear (red arrows) and diffusing the fragmented organelles (fm4, black and white arrowheads) into dense particles, which reassembled to increase the size of the nuclei and eventually joined together MN1 with the PN. Very close to the plasma membrane (double black arrows), MN2 appeared, which looked like a dispersed mitochondrion that had been condensed and changed into nuclear appearance via the further assembly of dense particles (black arrow). Electron-dense fm5 diffused into MN2, and fm6 separated to become part of MN2 (fm6/1) through the aggregation of the particles (**D1-D3**). (**D3**): a high magnification image of the inset in (**D1**). (**D4**) External aggregation of dense particles at the periphery in fm7 promoted nuclear growth (MN3 and MN4) while concurrently diluting itself and leading to its fusion with nuclei (double red arrowheads). Internal (fm8/1) and external (large red arrows) assembly occurred in fm8. Between the internal and external aggregation, electron-lucent intervals (white arrowheads) appeared. The intervals disappeared by the further assembly of dense particles in both fm8/1 and the aggregation of dense particles (large red arrows), which concomitantly caused fm8/1 to diffuse into nuclei (red arrowheads). fm8/1 diluted itself via consequently internal (black arrowhead) and external (small red arrow) aggregation of the particles, and the external assembly of dense particles at periphery of fm8/1 transiently formed a dark thread (small red arrow), which combined with the external aggregates of fm8 to look like tubule-NE fragment. Opposite white arrowheads: newly-formed nucleus between MN4 and the PN.

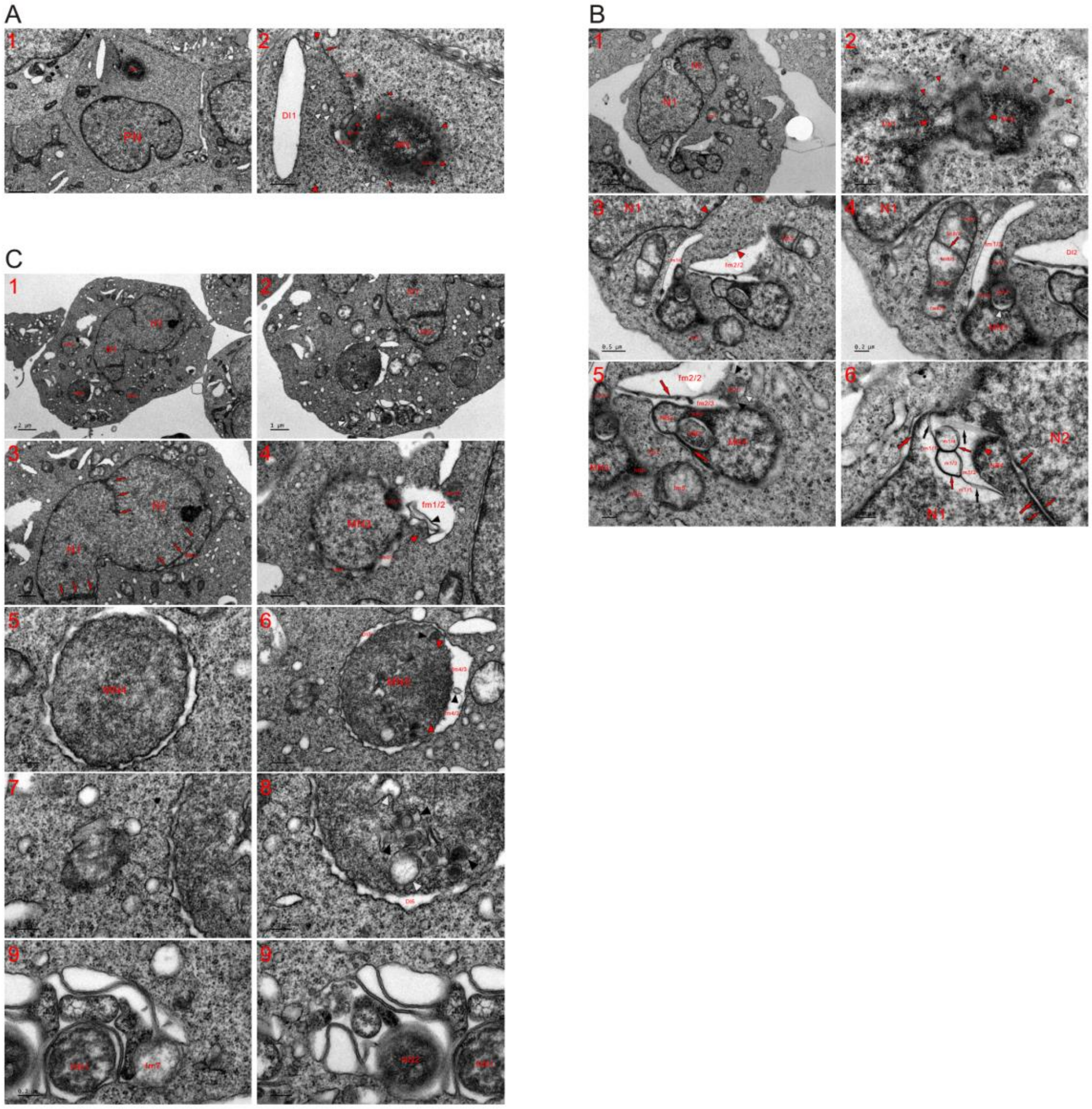

**Fig. S21. The formation of one or more micronuclei (MNs) in a single cell after a relatively short incubation time.** TEM was performed on three K562 cells at the 4 (**A**), 8 (**B**) and 12 (**C**) h time points, and the micrographs reveal that the formation of one or more MNs concurrently occurred with the growth of a large nucleus in a single cell. (**A**) Mitochondrial-derived dense particles (small red arrowheads) reassembled to form an MN, and the assembly of the particles separated fm1 (fm1/1, fm1/2 and white arrowhead) and caused the organelles to disperse into the MN (small red arrowheads). fm1 had diffused into it and become part of the nucleus, and dispersed mitochondria transiently formed virus-like granules (VLGs) at its edge (red arrowheads). Adjacent to the MN, dense particles from fragmented mitochondria (DI1, fm3, red arrow and white arrowheads) that subsequently promoted nuclear development (opposite red arrowheads), and the external aggregation of dense particles at the mitochondrial periphery formed DI1, which was electron transparent and had developed from a large mitochondrion (**A1** and **A2**). (**B**) MN1 merged with N2, and nuclear merging between MN1 and MN2 (opposite red arrowheads) was conducted by one or more dispersed mitochondria via the internal and external assembly of dense particles. Small red arrowheads: VLGs (**B1** and **B2**). (**B3-B5**) External assembly of dense particles in the organelles formed large electron-lucent structures (fm1/3 and fm2/2), which in turn enclosed an area of the cytoplasm with other mitochondria (such as fm3 and fm4) for nuclear development (opposite red arrowheads). The external aggregates of fm1 (fm1/1 and fm1/2) had become part of MN3, in which an internal aggregate of dense particles from other mitochondrion was observed (NB1 and white arrowheads); their internal assembly (red arrow) further divided the lucent part of fm2 (fm2/2 and fm2/3), and fm2/3 fused with MN4 by assembling the particles at its periphery while concurrently helping to form a NPB and promote the growth of the MN, in which mitochondria-derived nuclear bodies (NB2 and NB3) and a dark strand (large red arrow) were observed. Both NBs were localized in MN4 through their own aggregation of the particles, which further assembled to temporarily shape SDBs (black arrowheads) and SDIB (white arrowhead) within dense fm2/1 before totally dispersing into dense particles. At the edge of MN3, the particles from the organelles (such as fm5-fm7) reassembled to form a NBD, and less fragmented fm5 was going to disperse into MN4 as well as the surroundings through the aggregation of dense particles, which were going to separate (fm8/1-fm8/5). (**B6**) Along with the formation of NB4 and dark strand (large red arrow) via the assembly of dense particles at its periphery, the organelle (m1) became electron transparent and included the organelle itself into N1 simultaneously becoming nuclear localized. Their internal aggregation (red arrows) in m1 further divided its lucent part (m1/1-m1/4), while their further assembly led to the dispersion (red arrowhead and black arrows) of the dark aggregates and caused the nuclear tubules between nuclei (N1 and N2) to disappear. Two dark strands (large red arrows) combined to form nuclear tubules (opposite red arrows). (**C**) Traces of nuclear fusions were observed (three abreast red arrows), and in addition to the attached MNs (MN1 and MN2), three other MNs (MN3-MN5) were individually built in the cytoplasm (**C1-C3**). (**C4** and **C5**) External (fm1/1, fm1/3 and red arrowhead) and internal (black arrowheads) assembly of dense particles (red arrowheads) caused the organelle (fm1) to become electron transparent (fm1/2) and fuse with MN3, and at the nuclear edge, dispersed fm3/1 externally assembled dense particles resulting in the appearance of fm2/2. At the edge of MN4, the aggregation of the particles to their peripheries expanded the MN while concurrently causing the organelles to become electron lucent and allowing fragmented mitochondria to connect with each other to form a lunar halo structure. (**C6-C8**) Through their external assembly, the dense part of fm4 became part of MN5 (opposite red arrowheads), its lucent part was further divided (fm4/2 and fm4/3) by the internal aggregation of dense particles (black arrowheads), and their aggregation occurred at the nuclear edge (such as DI5 and DI6) as well as within the MN (black and white arrowheads). (**C9** and **C10**) Adjacent to the plasma membrane, fragmented mitochondria grouped together (opposite white arrowheads in **C2**) and tended to form nascent nuclei (NN1, NN2 and fm7) by assembling dense particles.

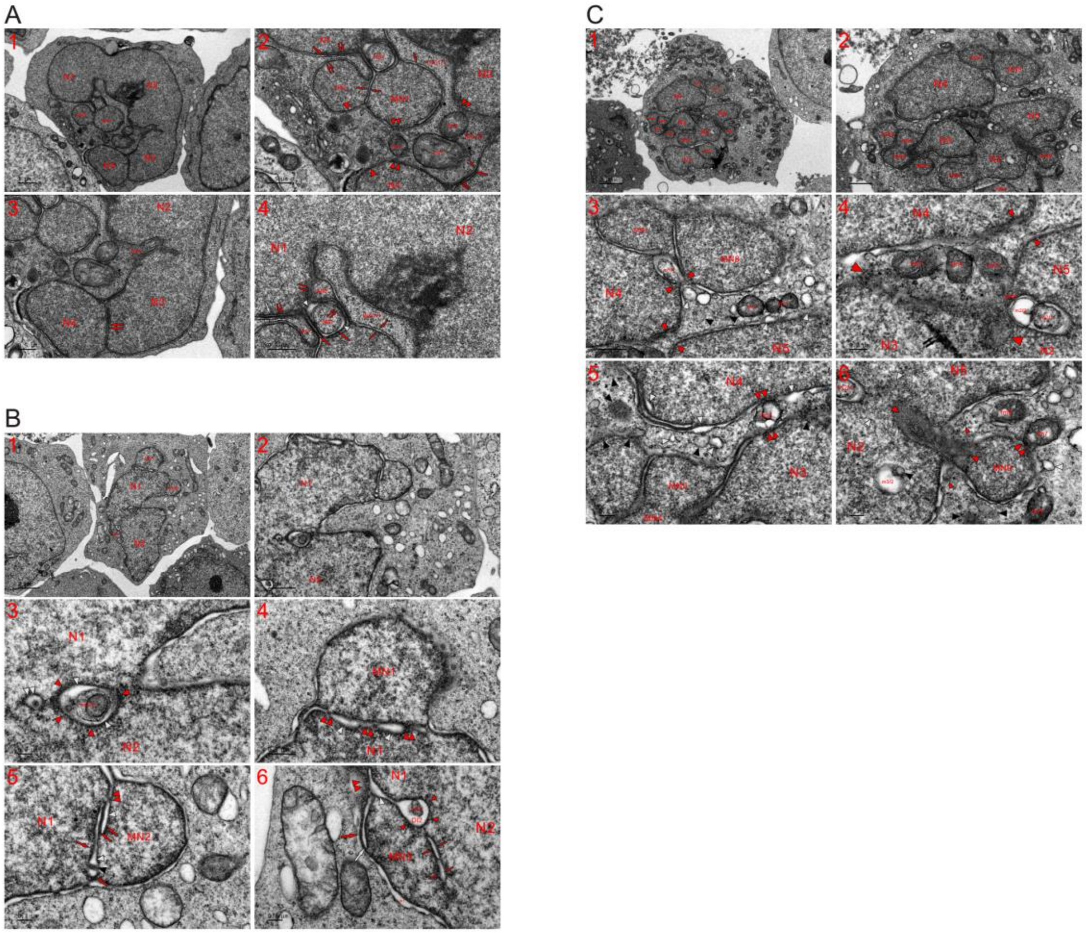

**Fig. S22. Separately formed nuclei joined together to build a large nucleus in a single cell at the 4, 8 and 12 h time points.** TEM was performed on three K562 cells at 4 (**A**), 8 (**B**) and 12 (**C**) h time points, and the micrographs demonstrate that the establishment of a large nucleus was a continuous process consisting of the assembly of dense particles in the organelles, mitochondrial fragmentation into the particles, their reassembly into small nuclei and nuclear fusion by fragmented mitochondria (**A-C**). (**A**) Nuclear fusions were mediated via the continuous nuclear transition of mitochondria to build a large nucleus in the cell, and the nuclear tubule disappeared (N3 and N4) or was going to disappeared (MN1 and MN2) between nuclei. Through the further assembly of dense particles, both dark strands (large red arrows) and threads (small red arrows) dispersed (double small red arrows) to provide particles for nuclear growth and nuclear merging (such as between MN1 and N1) while concurrently sealing the partitioned cytoplasm (INC2). Condensed mitochondria (fm1-fm4) assembled the particles to diffuse into nuclei (fm4 and double arrowheads) and into INC1/2, and the fragmented organelles were dispersed to become dense particles between MN2 and N4 (opposite large red arrowheads) (**A1-A3**). (**A4**) External and internal (NB1, NB2 and INC2) aggregation of dense particles occurred in the organelles, and along with their further assembly, tubule-NE fragments, nuclear tubule (white arrow), dark strands (large red arrows) and electron-lucent intervals disappeared (double red arrowheads) while the nuclear conversion of these internal aggregates occurred (NB1, NB2 and INC2). (**B**) Between N1 and N2, nuclear-localized m1 exhibited internal (m1/1) and external (red arrowheads) aggregation of dense particles, concomitantly creating electron-transparent intervals (white arrowheads), and adjacent to m1, the interval was disappearing (double white arrowheads) (**B1-B3**). (**B4** and **B5**) Along with the mitochondrial assembly of dense particles to promote nuclear growth (N1 and MN1), an electron-lucent gap (white arrowheads) appeared, which narrowed and was going to seal via the further assembly of the particles (double red arrowheads). The further aggregation of dense particles dispersed their former aggregates (red arrows and black arrowhead) into lucent lumens (white arrowheads) to merge N1 and MN2 (double red arrowheads). (**B6**) Aggregation of dense particles from both sides linked MN3 with N1 (double red arrowheads), and their internal assembly formed dark strands (large red arrow) and nuclear tubules (white arrow) at the edge of MN3. External assembly of dense particles at mitochondrial peripheries (the external assembly) promoted nuclear growth, concurrently creating different types of electron-transparent structures (b1, white arrowhead and opposite red arrows), and in addition to external aggregation, their internal congregation formed electron-opaque m2/1, concurrently shaping DI2 between m2/1 and the external assembling (small red arrowheads). (**C**) Five large nuclei and nine MNs were joined together to form a large nucleus, and fragmented mitochondria existed between or within nuclei (**C1** and **C2**). (**C3** and **C4**) The external assembly of dense particles from the organelles (such as m1 and white arrowheads) partially merged the nuclei (N4 and MN8) and was going to accomplish partial nuclear fusion between nuclei (N4 and N5) (opposite red arrowheads). Their internal aggregation occurred in both m1 (m1/1) and m2 (m2/1 and m2/2), part of which continuously conducted the external assembly of the particles to become electron transparent. Through further assembly of dense particles, condensed mitochondria (such as fm3-fm5) were dispersed to become particles, which reassembled to promote nuclear growth and expansion (opposite red arrowheads, NPB and double black arrows). (**C5**) Aggregation of dense particles caused the organelles to fuse with nuclei (fm6 and double red arrowheads; black and white arrowheads) and to turn into dense particles (black and white arrowheads), and the external assembly of dense particles occurred at its edge and within N4 (white arrowheads). (**C6**) Within N2, mitochondrial aggregation of dense particles was observed (electron-lucent m3/2 and black arrowhead), and the dispersion of condensed mitochondria (opposite red arrowheads) linked MN9 with large N2 and N5. Next to m2/1, mitochondrial assembling of dense particles for nuclear development displayed (opposite white arrowhead). Around MN9, less fragmented (such as fm7-fm9) and heavily or completely (black and white arrowheads) fragmented mitochondria turned into dense particles (red and double red arrowheads) via their assembly in the organelles.

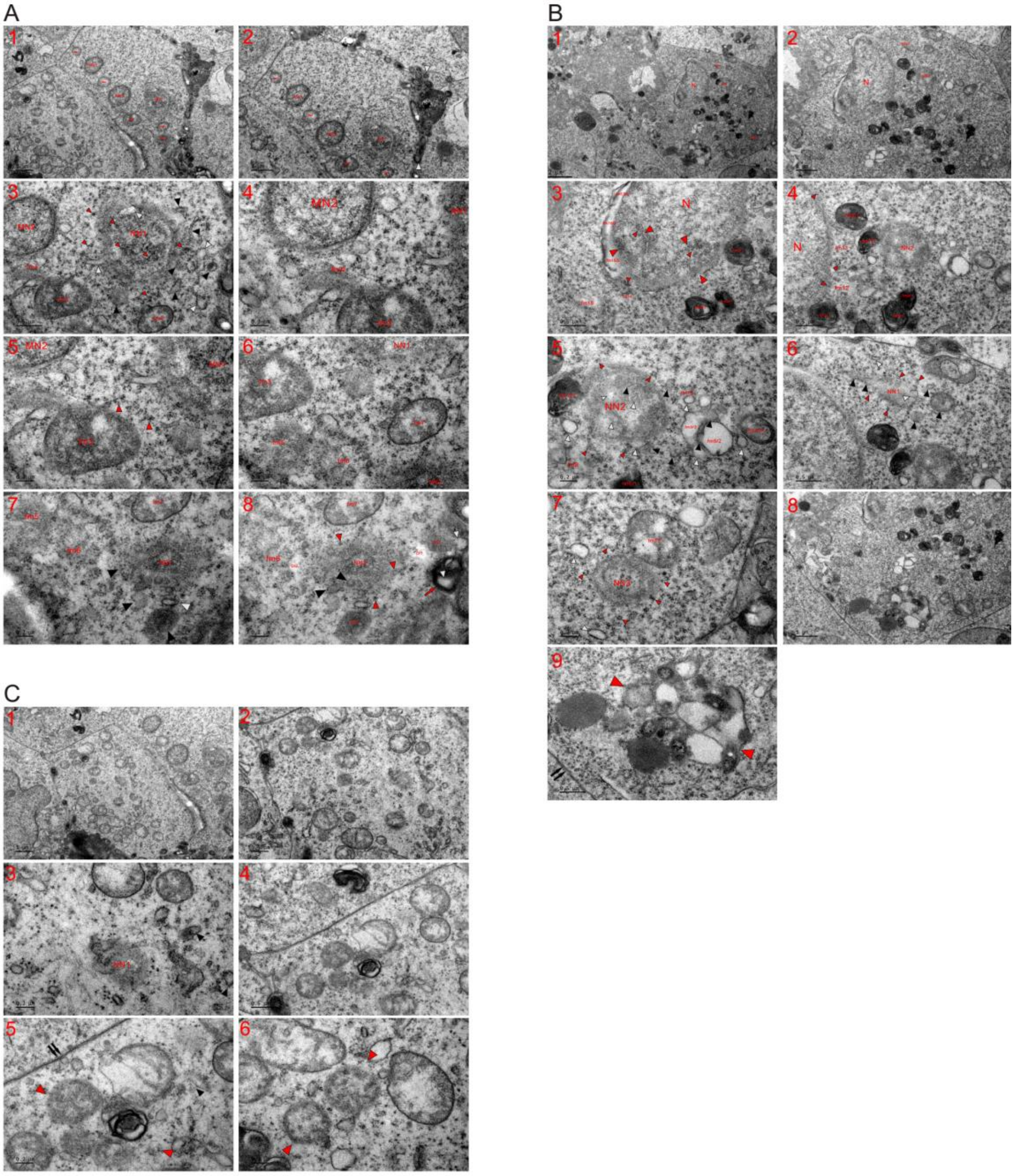

**Fig. S23. Initial formation of the nascent nucleus (NN) began with mitochondrial aggregation of dense particles at the 48 h time point.** TEM was performed on three K562 cells at the 48 h time point, and micrographs showed that the formation of an NN occurred in the cell with or without nuclei (**A-C**). (**A**) These cells possessed two small nuclei (MN1 and MN2), and two nascent nuclei were initiated (NN2) or enlarged (NN1) by fragmented mitochondria (**A1** and **A2**). (**A3-A6**) Heavily or completely fragmented mitochondria (black and white arrowheads) dispersed to become dense particles (small red arrowheads) at the edge of NN1 and within it, and their assembly in less fragmented fm3 and fm7 was going to diffuse these organelles into particles as observed with completely fragmented mitochondria (such as fm4-fm6). (**A7** and **A8**) Compared to NN1, NN2 was at a very initial stage of nuclear formation and looked like an aggregate of fragmented mitochondria (black, white and small red arrowheads), and traces of mitochondrial fragmentations via the aggregation of dense particles were demonstrated (DI1, DI2, white arrowhead, red arrow, DE1 and DE2). (**B**) In addition to a large nucleus (N), three NNs appeared in this cell (NN1-NN3), the internal assembly of dense particles condensed mitochondria (fm1/1-fm7/1), and their dilute parts dispersed in the cytoplasm as well as into nuclei (such as fm1/1 and fm7/1). Completely fragmented mitochondria had turned into dense particles (small red arrowheads) in the cytoplasm, at the nuclear edge (such as fm12-fm14) and within the large nucleus (opposite large red arrowheads). The separated mitochondria continued to fragment into dense particles (such as fm9/2, fm9/3, black and white arrowheads; fm11/1), and heavily fragmented organelles (such as fm8 and fm10) transiently formed SDBs (black arrowheads) and SDIBs (white arrowheads), both of which eventually dispersed into dense particles. The internal and external aggregation of dense particles in mitochondria (such as fm15-fm18) caused the organelles to appear at the edge of the large nucleus while concurrently becoming dilute or electron lucent (fm15/2-fm17/2) (**B1-B4**). (**B5** and **B6**) NN2 recently formed, and the aggregation of the particles appeared within (black, large and small white arrowheads) and surrounding (fm7/1, fm9/3 and fm10; black, white and small red arrowheads) NN2. NN1 looked like one fragmented mitochondrion, and the assembly of dense particles occurred within it and the neighboring fragmented organelles (small red, black and white arrowheads). (**B7**) NN3 looked like a dispersed organelle, and partial (fm19) or completely (small red and large white arrowheads) fragmented mitochondria diffused into it in the form of dense particles. (**B8** and **B9**) Adjacent to the plasma membrane (double black arrows), mitochondrial aggregation of dense particles diffused into each other to form a group, and the grouping tended to initiate nuclear formation (opposite red arrowheads). (**C**) No recognizable nucleus existed in this cell, which possessed a large number of mitochondria. Through the assembly of dense particles, mitochondria had changed into the appearance of an NN (NN1) or began to initiate nuclear formations (opposite red arrowheads). Double black arrows: plasma membrane (**C1-C6**).

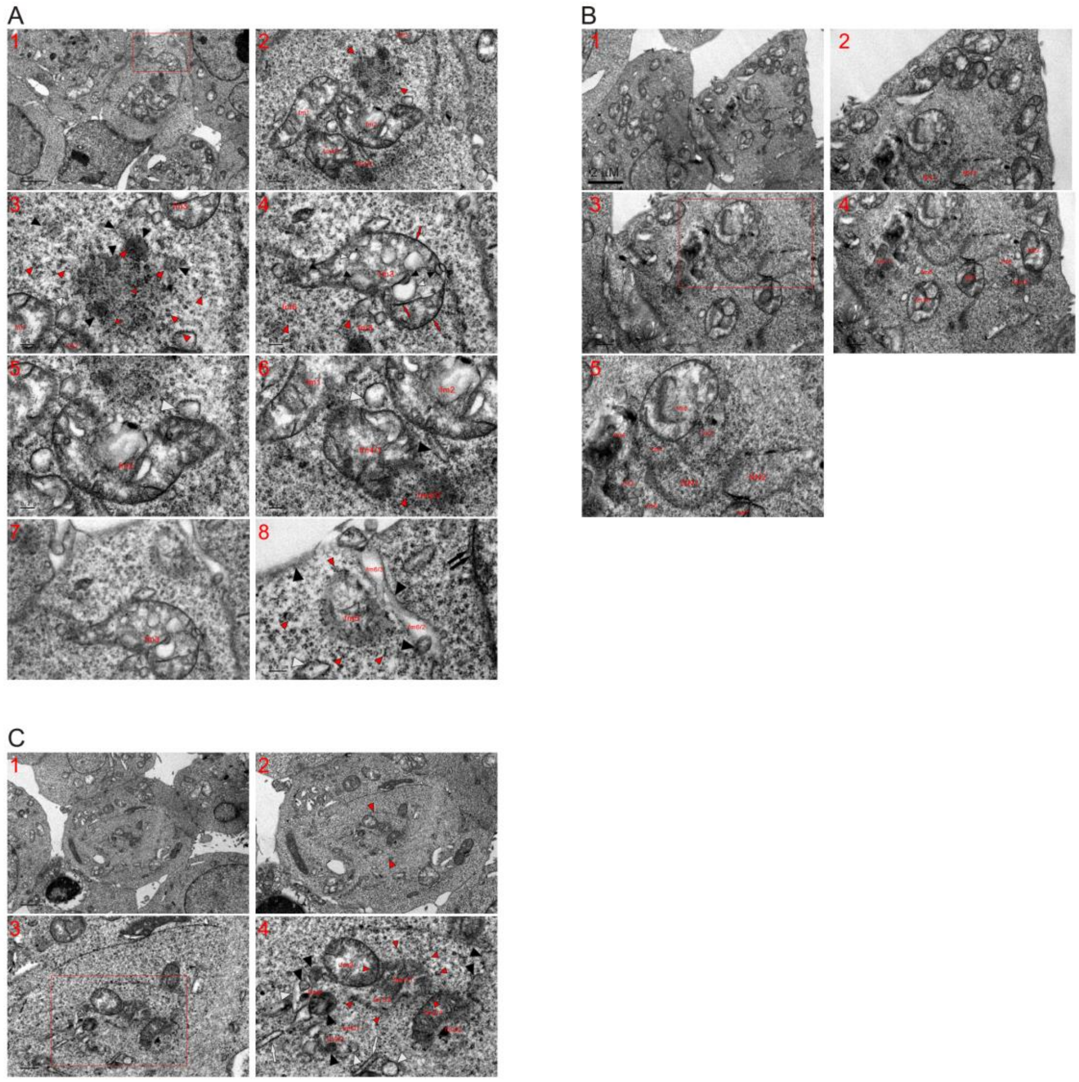

**Fig. S24. Mitochondria fragmented to initiate nuclear formation at the 2 h time point.** TEM was performed on three K562 cells at the 2 h time point, and the micrographs reveal that nuclear initiation began with the aggregation of fragmented mitochondria or mitochondrial assembling of dense particles (**A-C**). (**A**) The same cell shown in **Figure 3D**, and the micrograph of the whole cell shows that no recognizable nucleus it was present. Between less fragmented mitochondria (fm1-fm3), a group (opposite red arrowheads) of the fragmented organelles (black arrowheads) appeared, which had dispersed or were being dispersed into dense particles (large red arrowheads), and either SDBs (black arrowheads) or SDIBs (white arrowheads) were going to become the particles. Among the fragmented mitochondria, VLGs (small red arrowheads) were present. Completely fragmented mitochondria had dispersed into particles (red arrowheads) (**A1-A3**). (**A4-A6**) Through the internal (red arrow, black and white arrowheads) and external (red arrows) aggregation of dense particles (fm3), mitochondria were dispersed (fm1-fm3), and complete mitochondrial fragmentation (such as fm4/2, fm5 and fm6) caused the organelles to turn into dense particles (large red arrowheads). (**A7** and **A8**) Adjacent to the plasma membrane, mitochondria-derived dense particles (fm6/2, fm6/3, black arrowheads; white and small red arrowheads) were aggregated into fm5. (**A7**): a high magnification image of the inset in (**A1**). (**B**) Mitochondrial aggregation of dense particles resulted in the appearance of nascent nuclei (NN1 and NN2), and either NN1 or NN2 looked like a completely fragmented mitochondrion. Through the external and internal assembly of the particles, mitochondrial fragmentation occurred (such as fm1-fm12), and some of the organelles dispersed into the nascent nuclei (fm1-fm9) (**B1-B5**). (**B5**): a high magnification image of the inset in (**B3**). (**C**) Micrograph of whole cell shows that no recognizable nucleus was present in this cell and that a group of fragmented mitochondria appeared at a central part of its cytoplasm (opposite red arrowheads). The aggregation of dense particles in mitochondria caused the organelles to disperse in the form of dense particles (small red arrowheads) into the separated fm1 (fm1/1 and fm1/2). Before totally turning into particles, their aggregation divided the organelles (fm2/1 and fm2/2; fm4, black and white arrowheads), and due to mitochondrial fragmentations, intermediate ERLs (white arrows), SDBs (black arrowheads) and SDIBs (white arrowheads) appeared, all of which eventually diffused into dense particles.

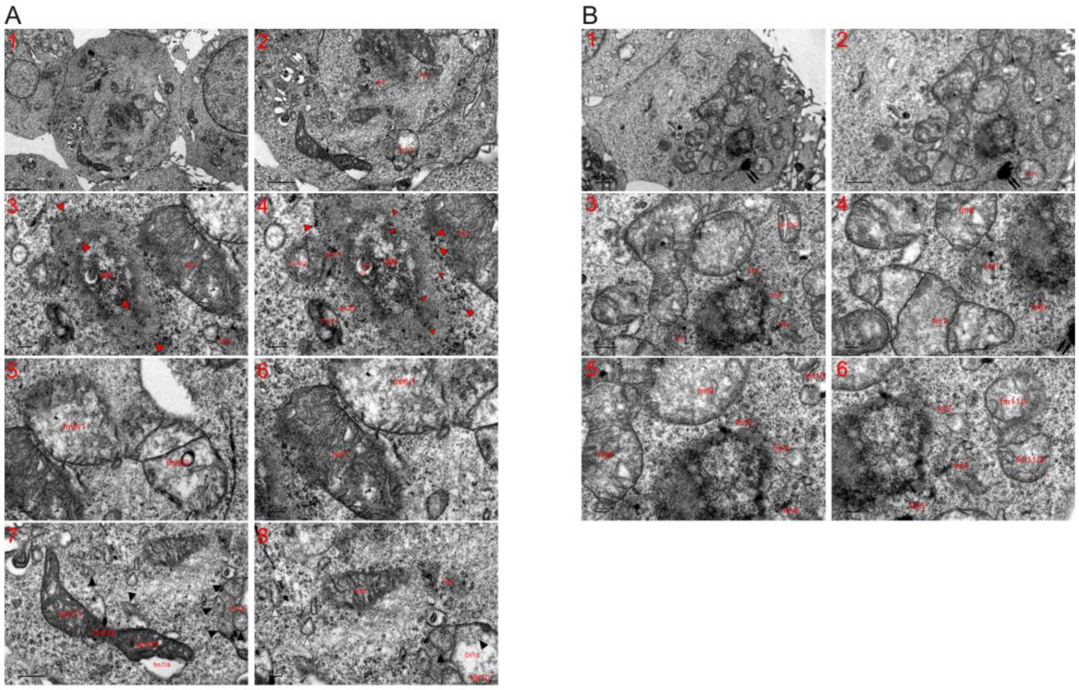

**Fig. S25. Mitochondria fragmented to promote the growth of the nascent nucleus at the 2 h time point.** TEM was performed on two K562 cells at the 2 h time point, and the micrographs reveal that fragmented mitochondria appeared within the NN and at its edge (**A** and **B**). (**A**) The same cell shown in **Figure 3D**. An NN appeared among fragmented mitochondria (such as fm1-fm5), which aggregated dense particles to disperse into it (opposite red arrowheads, fm1/1, fm2/2 and fm3). A mitochondrion included itself into the NN through the aggregation of the particles, whose internal assembly formed DE1 and lucent DI1 (**A1-A4**). (**A5-A8**) Through the assembly of dense particles, fm6 diluted and separated itself, concurrently dispersing into the particles, and condensed fm7 underwent further aggregation of dense particles (fm7/1-fm7/4). The assembly of dense particles began to diffuse electron-dense fm8, resulting in the appearance of complete fragmented fm9 and causing to the formation of SMLBs (black arrowheads) within fm10, concomitantly creating electron-transparent DI10. Either SDBs (black arrowheads) or SDIBs (white arrowheads) eventually turned into dense particles in the cytoplasm. (**B**) Partially fragmented (fm7-fm12) and heavily or completely (fm1-fm6 and double black arrows) fragmented mitochondria surrounded the NN to expand its size, and after the organelles that neighbored the NN had become part of the NN, mitochondria, which were somewhat far away from the nucleus and less fragmented, intensified their fissions to continuously provide dense particles for nuclear growth.

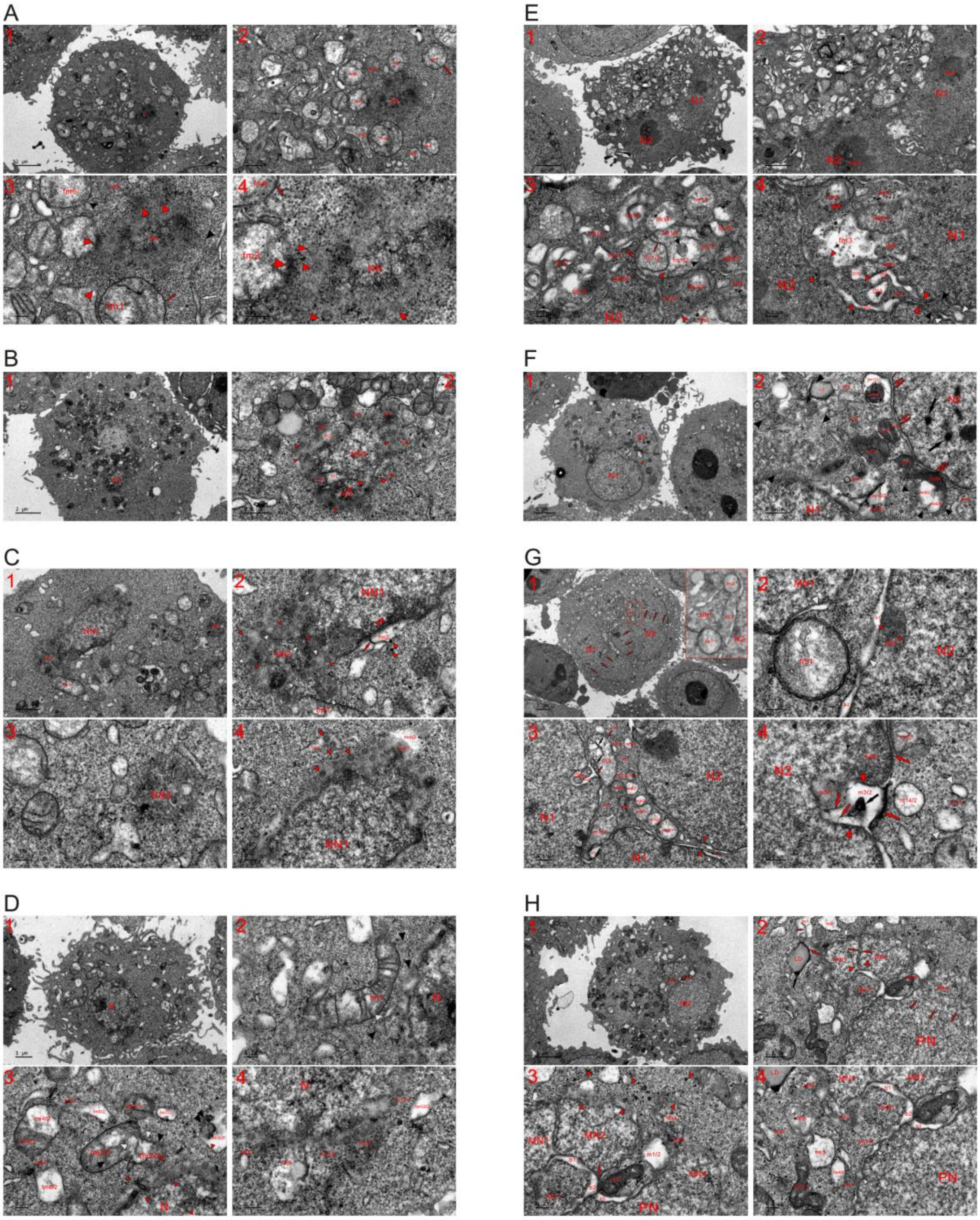

**Fig. S26. Mitochondria fragmented into dense particles to initiate nuclear formation and build nuclei in HeLa cells.** TEM was performed on eight HeLa cells at the 1 (**A-G**) and 3 (**H**) h time points, and the micrographs reveal that mitochondria not only fragmented to initiate and enlarge a nucleus but also joined together separately formed nuclei in a single cell. (**A**) The mitochondrial aggregation of dense resulted in the appearance of an NN, and dispersed mitochondria transiently formed

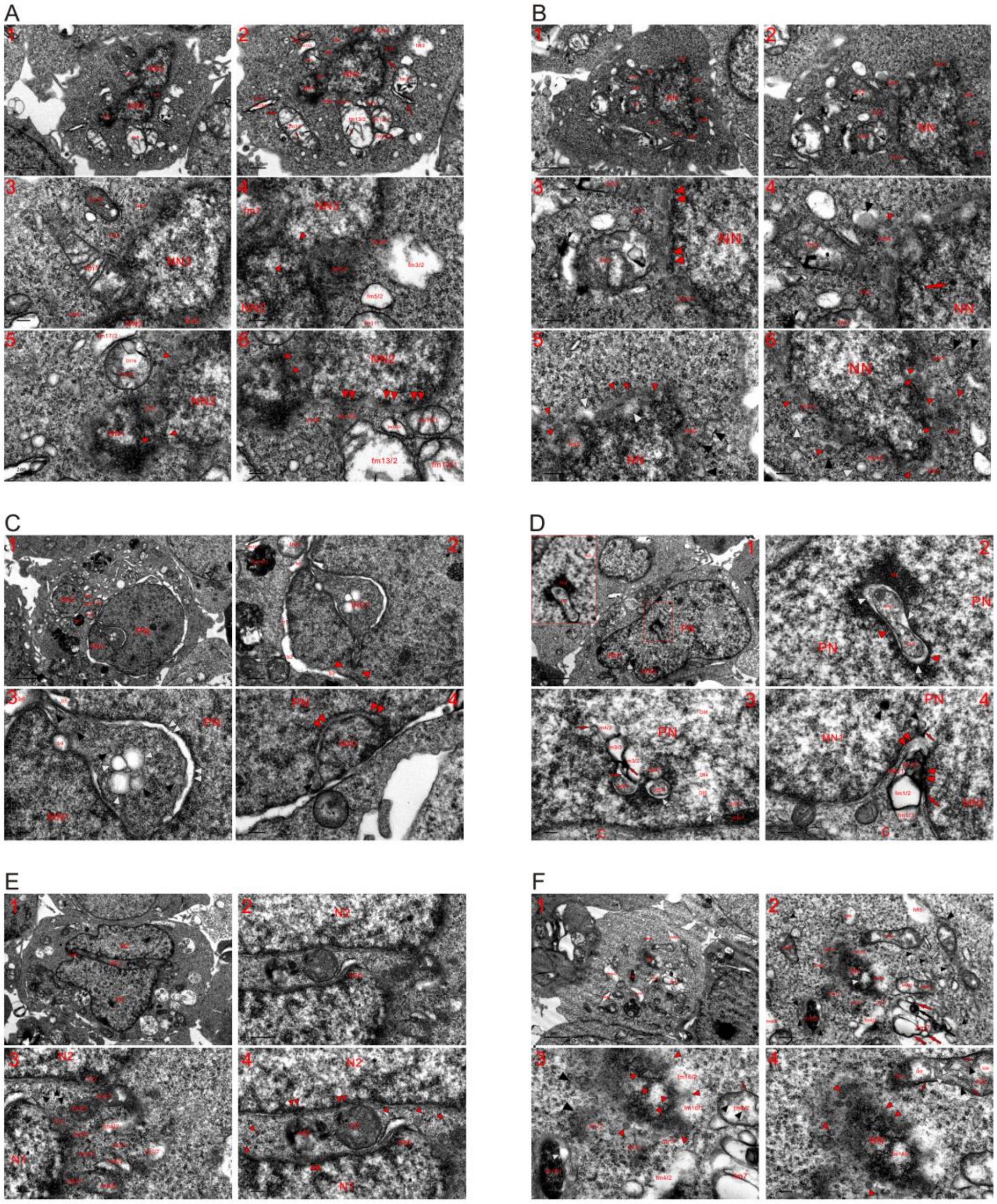


**Fig. S27. Mitochondria fragmented into dense particles to initiate nuclear formation and build nuclei in HEK293T cells.** TEM was performed on six HEK293T cells at the 2 (**A-E**) and 3 (**F**) h time points, and the micrographs reveal that the aggregation of dense particles in the organelles led to mitochondrial fragmentation into the particles, which in turn reassembled to initiate nuclear formation and build nuclei. (**A**) Three separately formed nascent nuclei (NN1-NN3)

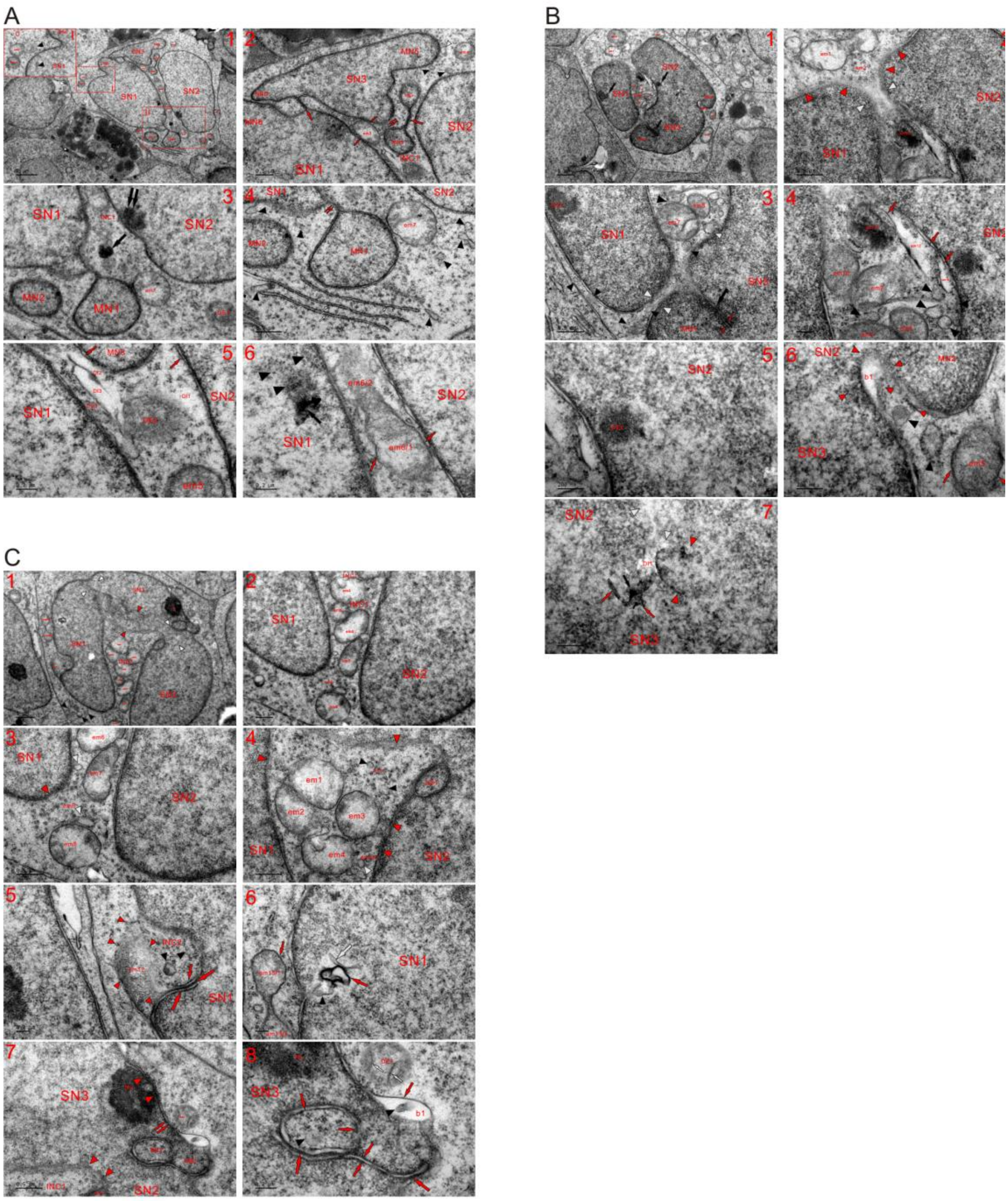


**Fig. S28. Mitogen stimulation from new medium obstructed nuclear fusion and growth.** TEM was performed on three K562 cells at the 5 min time point (switching the cells into new medium for 5 minutes after 48 h of continuous incubation), and the micrographs reveal that the new medium blocked nuclear fusion, prevented a large INC from sealing and led to mitochondrial renewal or re-establishment. (**A**) Gaps between nuclei widened, nuclear division occurred (SN1-SN3), and the attachment of micronuclei (MN1-MN4) was obstructed or micronuclei were detached. High magnification image of inset I reveals that mitochondria detached from the nucleus (em1 and em2) and that dense particles assembled into bodies with mitochondrial morphology (black arrowheads). The aggregation of dense particles separated NBD and MN6, let dark strands to disperse (large red arrows), detached em3 by forming its LMM (small red arrows), discontinued diffusion of em4 and was going to divide nuclei (MN3 and SN3) (**A1** and **A2**). (**A3** and **A4**) MN1 was detached from SN1 by dispersing the dark strand into dense particles (double red arrows), whose aggregation at nuclear edges prevented MN2 from joining together with SN1 and reversed diffusion of em7 to its renewal. Recovery of em7 completely connected INC1 with the cytoplasm outside of nuclei, and dispersion of dense aggregates hindered in either SN2 (double black arrows) or INC1 (black arrow), and dispersions of elongated ERLs (white arrows), SDBs (black arrowheads), SDIBs (white arrowheads) and DE1 obstructed in the cytoplasm. (**A5** and **A6**) New medium led to aggregation of dense particles for mitochondrial renewals (such as em5 and em6) and re-establishments (DE2, DE3 and DI1-DI3) concurrently forming the LMM (small red arrows). Rather than dispersing, the particles assembled in SN1 (black arrow and arrowheads). SN: separated nucleus; em: emerging or nascent mitochondria. (**B**) Fresh medium aggregated dense particles in nuclei (black arrows) to widen nuclear intervals (opposite white arrowheads) and to lead their return to the cytoplasm (red arrowheads), where mitochondrial reinstatement (such as em1, em2 and em11-em13; black and white arrowheads) and recovery (em4-em10) happened. The assembly of dense particles discontinued dispersion of em11, became more obvious in nuclei (DE1, DE2, large black and opposite red arrowheads) and let them return back to electron-lucent organelles (such as em12 and em13; red arrows) (**B1-B5**). (**B6** and **B7**) Diffusion of dense em5 blocked with reappearing of the LMM (red arrows) and aggregation of dense particles in nuclei (MN2 and SN3) widened the nuclear interval, concurrently returning of the particles (small red arrowheads) for mitochondrial re-establishment (b1 and black arrowheads). Between SN2 and SN3, dense particle aggregation occurred (black and red arrows; opposite arrowheads) and enlarged electron-transparent areas (such as DI1 and white arrowheads). (**C**) Upon stimulation with new medium, all large mitochondria detached from the nuclei within INC1 (em1-em7), nuclear divisions occurred (opposite white and red arrowheads), dense particles returned from nuclei (red arrowheads) and mitochondrial renewal (em8) or regeneration occurred (em9-em11), and for those completely fragmented, mitochondrial re-establishments began with the formation of SMLBs by assembling the returned dense particles (black and white arrowheads). At the edge of SN2, dense particles reassembled into a body of mitochondrial shape (NB1) (**C1-C4**). (**C5** and **C6**) Dispersion of em12 was discontinued, the return of dense particles (small red arrowheads) promoted its recovery into an orthodox mitochondrion, and the formation of SMLBs (black arrowheads) and a dark thread (small red arrow) promoted mitochondrial re-establishment in INC2, which became more obvious along with aggregation of the particles (em12; small and large red arrows; black and small red arrowheads). Within SN1, diffusion of the ERL (white arrows), a SMLB (black arrowhead) and dark strand discontinued, or all these structures reemerged by aggregation of dense particles in order for mitochondrial reinstatement. At the nuclear edge, fm13 detached from SN1 by forming LMM (red arrow) and recovered its dumbbell shape. (**C7** and **C8**) The return of dense particles from the nucleus, Nu and tubule-NE fragment to the cytoplasm (red arrowheads and double red arrows) was observed, and their aggregation enlarged INC1, reemerged nuclear bodies (NB2-NB3), discontinued the dispersion of dark strands (large and opposite small red arrows), formed the LMM (small red arrows), obstructed diffusion of DE1 and recombined electron-opaque NB3 with lucent b1, in which an SDB was being dispersed (black arrowhead). Within DE1, which lacked an LMM, dense particles assembled to be cristae-like structures (white arrows).

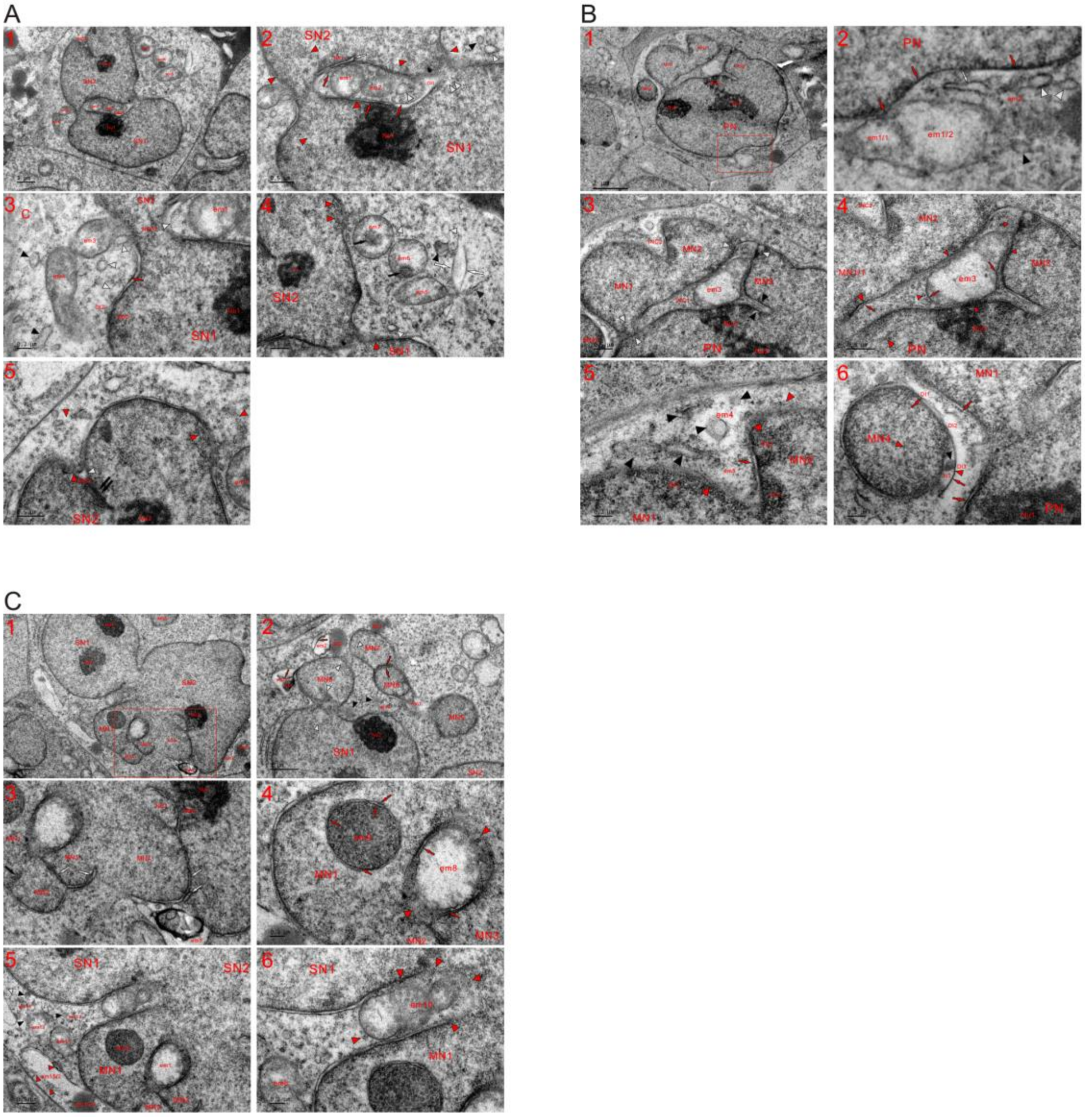

**Fig. S29. New medium reversed nuclear building to mitochondrial recoveries and re-establishments.** TEM was performed on three K562 cells at the 5 min time point, and micrographs demonstrated that new medium blocked nuclear conversion of both cytoplasmic mitochondria and those enclosed mitochondria, and aggregation of dense particles induced by the mitogen led to nuclear division, concurrently reversing nuclear building to allow for mitochondrial generation and regeneration. (**A**) Dispersion of mitochondria in INC1, which became more obvious and was opened by aggregation of dense particles (DI1, double white and opposite red arrowheads), discontinued (em1, em2, and white arrowheads). At the edge of INC1, assembled dense particles (NB1, white arrow, red arrows and opposite red arrowheads) were observed, whose aggregation was going to make NPB1 disappear. The detachment of mitochondria following their recoveries was observed (em3-em7), NB2 was going to recombine with DI2, and the formation of SMLBs (black and white arrowheads) and a dark thread (red arrow) promoted mitochondrial re-establishments in the cytoplasm. The assembly of dense particles in nuclei returned them back to the cytoplasm (red arrowheads) and spread dense bodies (black arrows) within either em6 or em7, and their return to the cytoplasm led to the formation of SMLBs (black and white arrowheads) and the large mitochondrion (white arrows) (**A1-A4**). (**A5**) The return of dense particles from the nucleus together with their aggregation in the cytoplasm benefited mitochondrial re-establishment (opposite red arrowheads), and their aggregation in SN2 (double black arrows and white arrowhead) enlarged INC2, concurrently leading to nuclear division. (**B**) Instead of dispersion into the PN, the re-establishment of the organelles occurred (em1 and em2) concurrently with the formation of a dark thread (red arrow), the ERL (white arrow) and SMLBs (black and white arrowheads) (**B1** and **B2**). (**B2**) A high magnification image of the inset in (**B1**). (**B3-B5**) Diffusion of em3 discontinued, and instead, it was recovered to be an orthodox mitochondrion by returning dense particles (small red arrowheads), which assembled to become LMM at its periphery (red arrows). Adjacent to em3, the formation of dark thread (red arrow) promoted mitochondrial re-establishment (opposite red arrowheads), and along with aggregation of dense particles, linkages between nuclei inclined to break up (opposite white arrowheads), concurrently connecting INC1 with the nonpartitioned cytoplasm (cytoplasmic area outside of the nucleus). The aggregation of dense particles widened nuclear intervals (INC2 and opposite black arrowheads), returned the particles back into the cytoplasm (red arrowheads), caused to their assembly at the nuclear edge (such as DE1-DE3) and was in favor of mitochondrial reinstatements (em4, em5, red arrow and black arrowheads). (**B6**) Fresh medium discontinued joining together of individually formed MN4 with the PN by enlarging electron-lucent areas (DI1-DI3 and b1) and reemerging dark threads (red arrows), and recombination of b1 with aggregates of dense particles demonstrated at the edge of the MN (black and opposite red arrowheads). (**C**) Nuclear fusions discontinued (MN5 and MN6; MN5 and SN1; MN7 and MN8) and instead, new medium led to their divisions through assembling dense particles (opposite white arrowheads and red arrows). The diffusion of these dense bodies into nuclei blocked (such as DE1-DE3) and mitochondrial re-establishments for those completely fragmented were being underwent (em1-em4, DE1-DE3 and black arrowheads), and aggregates of the particles (DE1, DE2 and red arrows) dispersed into electron-transparent parts of the organelles (such as em2 and em3) (**C1** and **C2**). (**C3**) A high magnification image of the inset in (**C1**) shows that nutrient supplementation increased aggregations of dense particles in nuclei (NB1, NB2, black and white arrows) and blocked attachments of micronuclei (MN1-MN4). At the nuclear edge, The organelles were regenerated (em5-em7) (**C1**). (**C4**) In the nucleus, diffusions of both em8 and em9 discontinued being blocked, and through assembling of their owns, dense particles returned to electron-transparent em8 to allow for its recovery (red arrows and arrowheads). Aggregation occurred within dense em9 and at its edge (red arrows), and internal assembly caused em9 to look like an MBV. (**C5** and **C6**) The return of dense particles to em10 occurred in both the cytoplasm and nuclei (small red arrowheads), and mitochondrial re-establishment (such as em11-em14, black and white arrowheads) occurred at the nuclear edge. The dispersion of its dense part (em15/1) into MN1 discontinued upon stimulation with new medium, and em15 was being re-established by the return of the particles (red arrowheads) and the recombination of em15/1 with lucent em15/2.

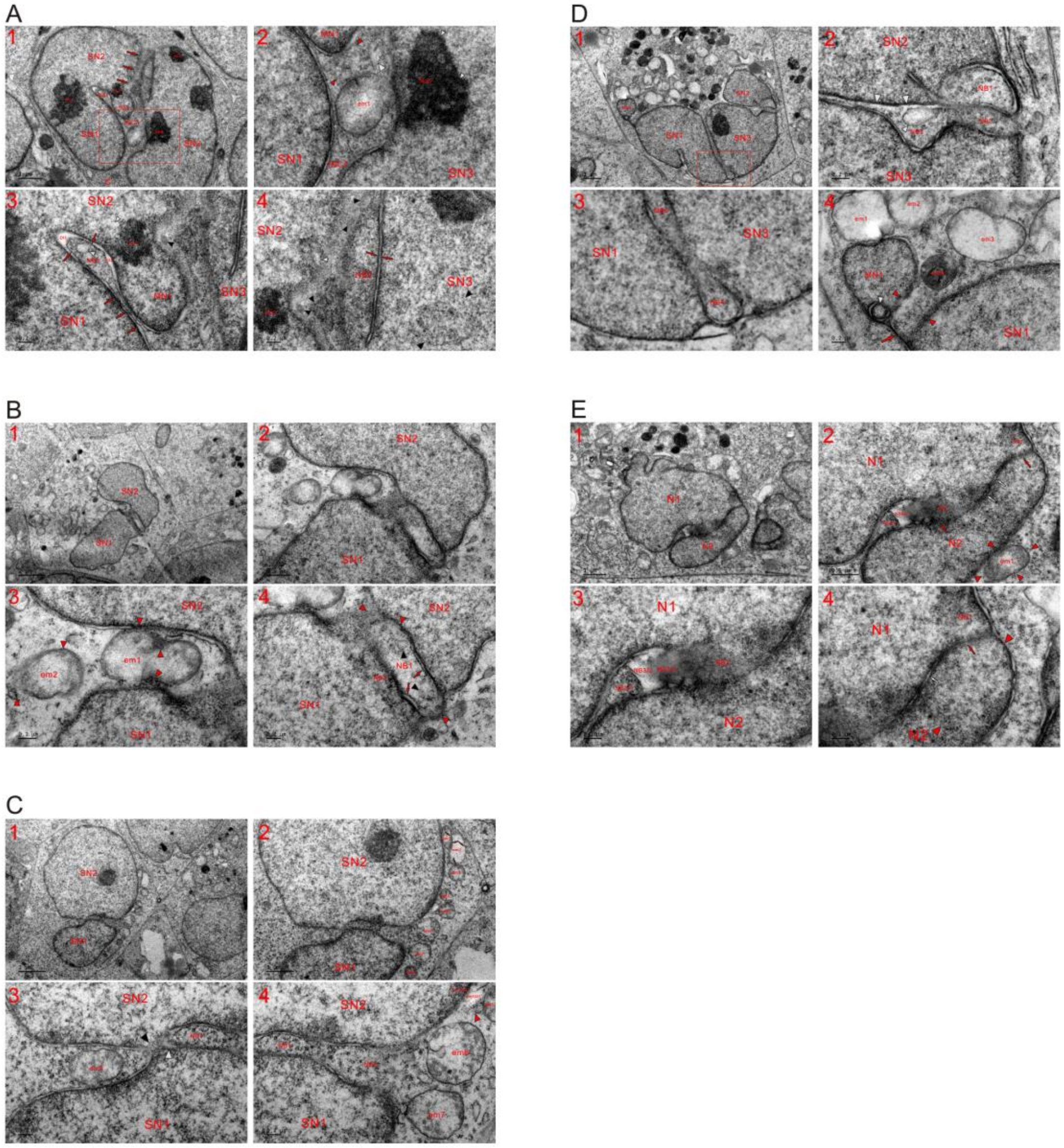


**Fig. S30. Nutrient supplementation isolated nuclei into bodies of mitochondrial morphology by assembling dense particles.** TEM was performed in five K562 cells at the 5 min time point, and micrographs showed that nutrient supplementation reversed nuclear formation of the organelles to mitochondrial renewal and re-establishment, and the assembly of dense particles led to the formation of nuclear bodies of mitochondrial morphology. (**A**) Aggregation of dense particles separated nuclei (SN2 and SN3; three abreast red arrows) and led to disappearance of the partitioned cytoplasm (INC1). New medium assembled the particles for mitochondrial recovery (em1) and re-establishment (opposite red and white arrowheads) (**A1** and **A2**). (**A2**) A high magnification image of the inset in (**A1**). (**A3**) Nuclear fusion between SN1 and MN1 was discontinued (three abreast red arrows), and aggregation of dense particles in the nucleus (Nu2, black and white arrowheads) could lead to detachment of MN1. Between nuclei (SN1 and SN2), the particles reassembled for mitochondrial regeneration (NB1, DI1, DI2, white arrowheads and red arrows). (**A4**) Congregation of dense particles (opposite red arrows, black and white arrowheads) instead of their intermixing occurred between nuclei (SN2 and SN3) and formation of nuclear tubules (opposite red arrows) isolated part of SN3 into a body of mitochondrial morphology (NB2). (**B**) Aggregated dense particles returned (red arrowheads) to recover mitochondria (such as em1 and em2) and reversed the nuclear transition of the organelle for mitochondrial re-establishment (NB1, NB2, red arrows, black and red arrowheads) (**B1-B4**). (**C**) Fresh medium detached mitochondria from nuclei (such as em1-em9), led to nuclear separation (SN1 and SN2) via aggregation of dense particles (black arrowheads) and caused the organelles to separate by assembling the particles into LMM (em1-em3 and small red arrows). At the edge of SN2 and adjacent to em5, dense particles were returned back into electron-transparent em10/2 from the surrounding (red arrowhead) and em10/1, which was dense and located within SN2. Between SN1 and SN2, nuclear bodies with a mitochondrial shape were formed (NB1 and NB2) (**C1-C4**). (**D**) Mitogen stimulation with new medium led to nuclear divisions (N1-N3 and MN1) by aggregating dense particles, whose assembly widened the electron-lucent interval (large white arrowheads) between nuclei (SN2 and SN3) and reversed nuclear formation of mitochondria to form the bodies of mitochondrial morphology in them (NB1-NB4 and small white arrowhead) (**D1-D3**). (**D4**) Dense particles returned back to electron-transparent mitochondria for their development into orthodox ones (em1-em3), dispersion of dense em4 into nuclei blocked, a SMLB reemerged within MN1 (white arrowhead) and dark strand was dispersed to provide dense particles for mitochondrial re-establishment (opposite red arrowheads). (**E**) The reassembly of dense particles occurred (NB1-NB3, red and white arrowheads) between nuclei (SN1 and SN2), and dense NBs (NB3/1 and NB3/3) were recombined with lucent NB3/2; within N1, the aggregation of the particles caused part of it to become a spherical body (opposite red arrowheads, red and white arrows) (**E1-E4**).

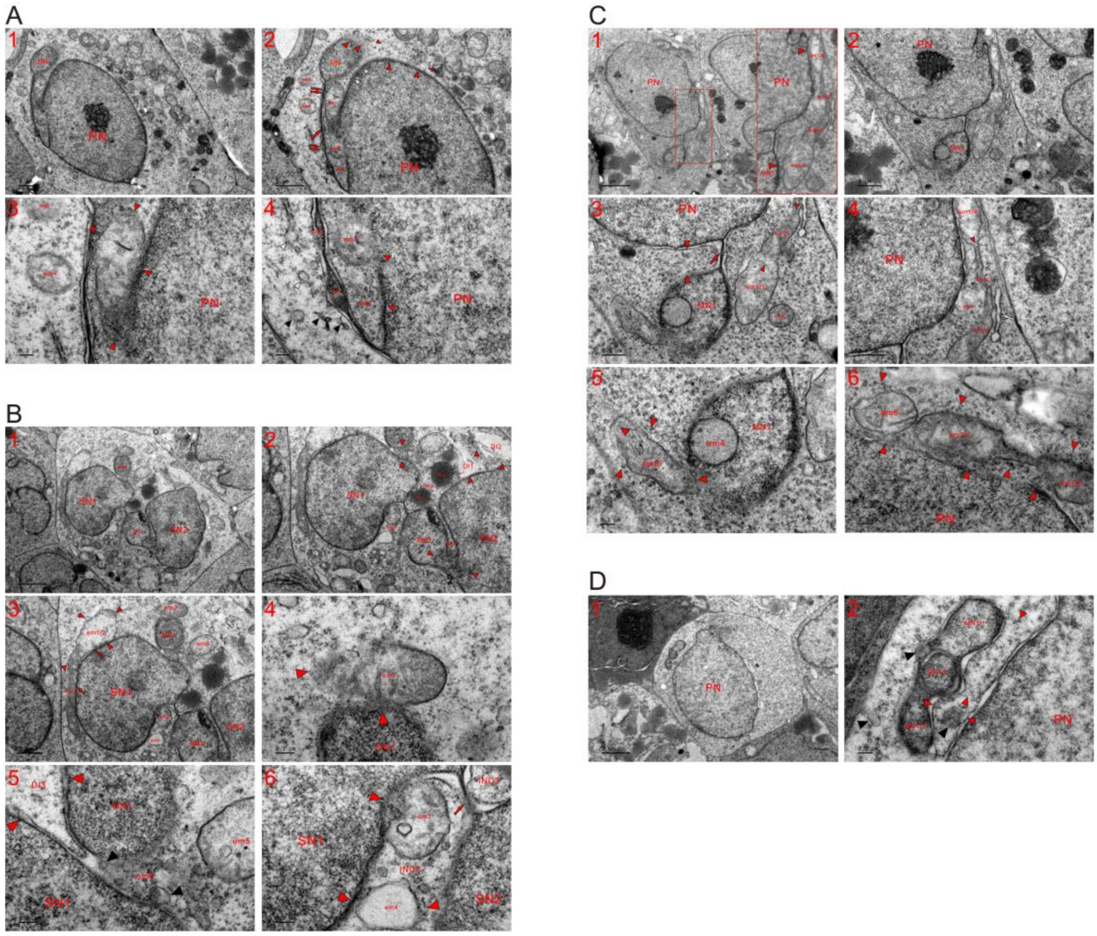

**Fig. S31. Fresh medium discontinued attachment of an MN to the large nucleus and blocked nuclear fusion.** TEM was performed on five K562 cells at the 5 min time point, and micrographs revealed that fresh medium discontinued the growth and attachment of an MN to the PN (or a large nucleus) by the return of dense particles for mitochondrial renewal and regeneration. (**A**) Instead of dispersion for nuclear growth, the dark strand (large red arrow), which linked MN1 with the PN, was dispersed (double small red arrows) to return dense particles back into the cytoplasm, and their return from the PN (red arrowheads) renewed em1 and re-established those completely fragmented (em2 and em3 and opposite red arrowheads) at the edge of the PN. The aggregation of dense particles widened the dilute interval between nuclei (white arrowheads), discontinued dispersion of the organelles in the cytoplasm (such as em4 and em5), formed SMLBs (black and white arrowheads) and diffused DEs (DE1 and DE2), and the formation of SMLBs and dispersion of aggregates of dense particles benefited mitochondrial reinstatements in the cytoplasm. (**B**) Aggregation of dense particles reversed nuclear conversion of the organelles to mitochondrial renewals (em1-em5), enlarged INCs (INC1-INC3) and blocked nuclear fusion (MN1 and SN1; opposite red arrowheads) or caused nuclear divisions (SN2 and MN2; opposite red arrowheads). The dispersion of lipid droplets (LD1 and LD2) into nuclei was obstructed, and the return of dense particles (red arrowheads) for re-establishment was observed (such as DI1-DI3). Along with the mitochondrial recovery induced by the return of the particles (red arrowheads), the elongated em1 was detached from SN1 (red arrows), and em2 was going to separate from MN1. Between SN1 and MN1, mitochondrial reinstatement was observed (fm6 and black arrowheads). Between INC2 and INC3, the dark strand was being dispersed (red arrow) to provide dense particles for mitochondrial formation and to break up the linkage between nuclei (SN1 and SN2), concurrently allowing the partitioned cytoplasm (INC2 and INC3) to disappear, and both em3 and em4 promoted the development of orthodox mitochondria with the return of the particles (**B1-B6**). (**C**) High magnification image of the inset reveals that elongated em1 detached from either the PN or MN1 by returning dense particles (red arrowheads). Their return led to the recovery of em1, and their internal aggregation separated em1 (em1/1-em1/4 and small red arrowheads), and adjacent to em1, the diffusion of an ERL discontinued or was restored (white arrow). In the cytoplasm, the aggregation of dense particles recovered em2, dispersed a dark strand to free MN1 (red arrow) and re-established the organelle between nuclei (opposite red arrowheads) or between the PN and em1 (such as em3) (**C1-C4**). (**C5** and **C6**) Within MN1 and at nuclear edges, the particles (red arrowheads) returned back for mitochondrial renewals (such as em4-em7), and em7 displayed a dumbbell-shaped morphology. (**D**) New medium discontinued nuclear fusion between N1 and MN1, which was separated into bodies of mitochondrial morphology by assembling dense particles within it, and adjacent it, dense particles aggregated for mitochondrial development (opposite red and black arrowheads) (**D1** and **D2**).

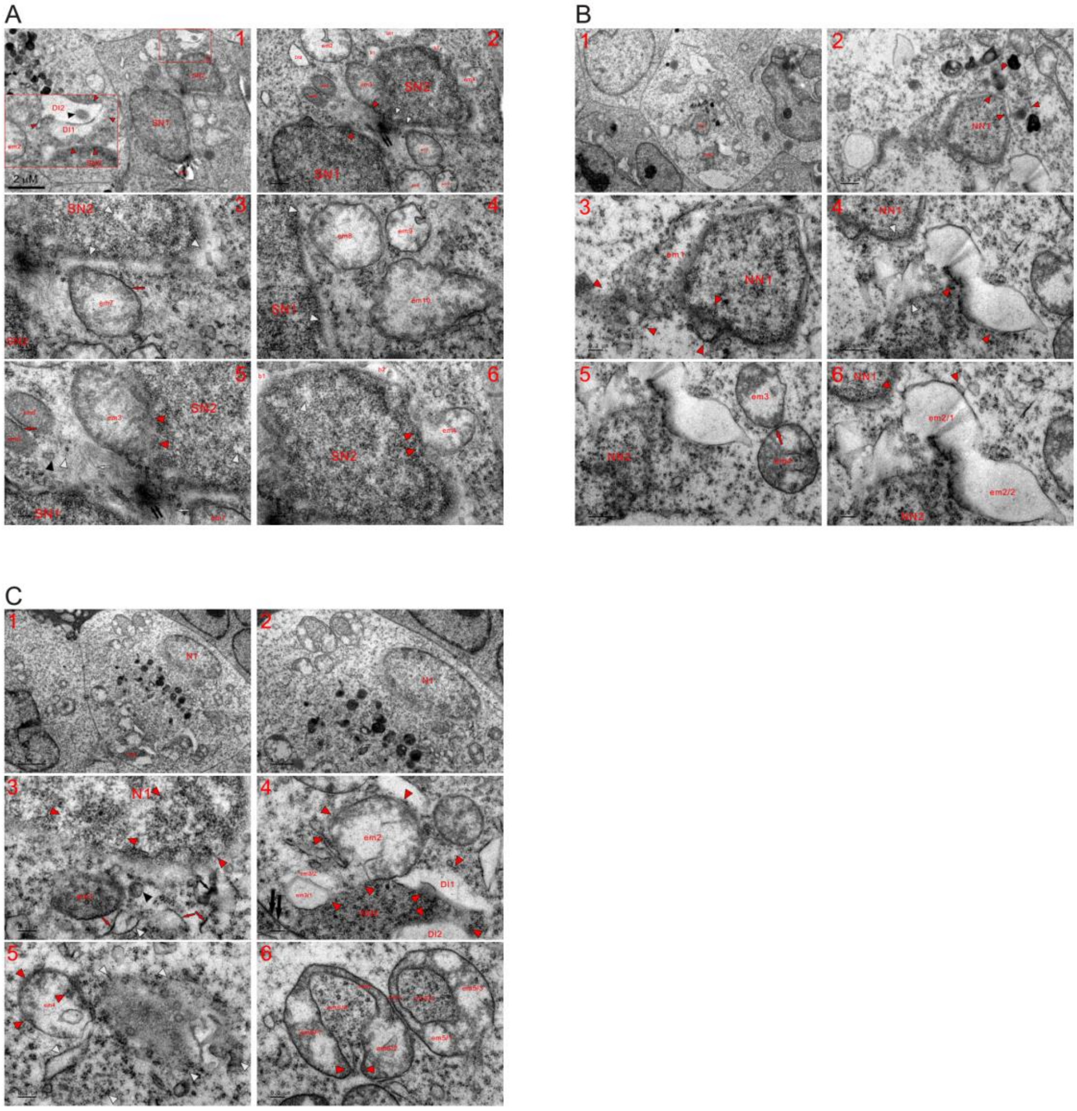

**Fig. S32. Nutrient supplementation discontinued nuclear initiation and blocked the building of a nucleus at its early stage.** TEM was performed on three K562 cells at the 5 min time point, and the micrographs demonstrate that both nuclear growth and initiation were blocked upon challenge with new medium. (**A**) At the edge of SN1, dense particles assembled to re-establish em1, and a high magnification image of the inset revealed that dense particles were returned back into DIs (DI1 and DI2) and DI2 contained a dispersed SDB (**A1**). (**A2-A6**) The return of dense particles benefited the renewal of electron-transparent organelles (such as em2 and em8-em10) and led to the re-establishment of completely fragmented mitochondria (DI3, b1 and b2). At the edge of SN2, the dispersion of both organelles (em3 and em4) was discontinued, and instead they changed into orthodox mitochondria with the return of dense particles (red arrowheads), whose aggregation (black arrow and white arrows; black and white arrowheads) widened the gap (opposite red arrowheads) between nuclei (SN1 and SN2) and created lucent (or dilute) structures either within nuclei or at nuclear edges (white arrowheads). Their aggregation reversed the diffusion of the dense organelles (em5 and em6) to promote mitochondrial recovery and separation by forming LMM (red arrow). Red arrows: microfilaments. (**B**) Two small nuclei appeared in the cell (NN1 and NN2), and they looked like at the early (NN1) or initial (NN2) stage of nuclear formation, and around NN1, mitochondrial fission or diffusion was blocked (opposite red arrowheads). At the edge of NN1, the aggregation of dense particles led to mitochondrial re-establishments (em1 and opposite red arrowheads). The assembly of dense particles widened the gap between nuclei, returned them back (red arrowheads) to the electron-lucent organelles (em2/1 and em2/2; opposite white arrowheads) for mitochondrial regeneration and separated mitochondria (em3 and em4) by forming LMM (red arrow) (**B1**-**B6**). (**C**) Dispersion of electron-opaque em1 blocked at the nuclear edge (N1), where dense particles were assembled into nuclear bodies of mitochondrial morphology (opposite red arrowheads), and adjacent to N1, their assembly (black and red arrows; black and white arrowheads) benefited mitochondrial reinstatements in the cytoplasm (**C1-C3**). (**C4** and **C5**) Very next to the plasma membrane (double black arrows), mitochondria-derived dense particles had evolved into the appearance of an NN (NN2) under nutrient-limited conditions, and new medium led to their return to electron-lucent mitochondria (em2 and em3) or phases (DI1 and DI2) from either NN2 or the cytoplasm (red arrowheads). Adjacent to em4, fragmented mitochondria should have developed to initiate nuclear formation under nutrient-poor conditions, and presently, they were re-established (opposite white arrowheads) or recovered (em4) by reassembling the particles. (**C6**) Internal (such as em5/4 and em6/4) and external (em5/1-em5/3 and em6/1-em6/3) aggregations of dense particles had occurred in the organelles under nutrient-poor conditions, and challenge of new medium led to their recoveries by reassembling of dense particles from both within (em5/4 and em6/4) and outside of mitochondria. Along with the aggregation of the particles, the gap between em6/1 and em6/2 could be sealed (opposite red arrowheads).

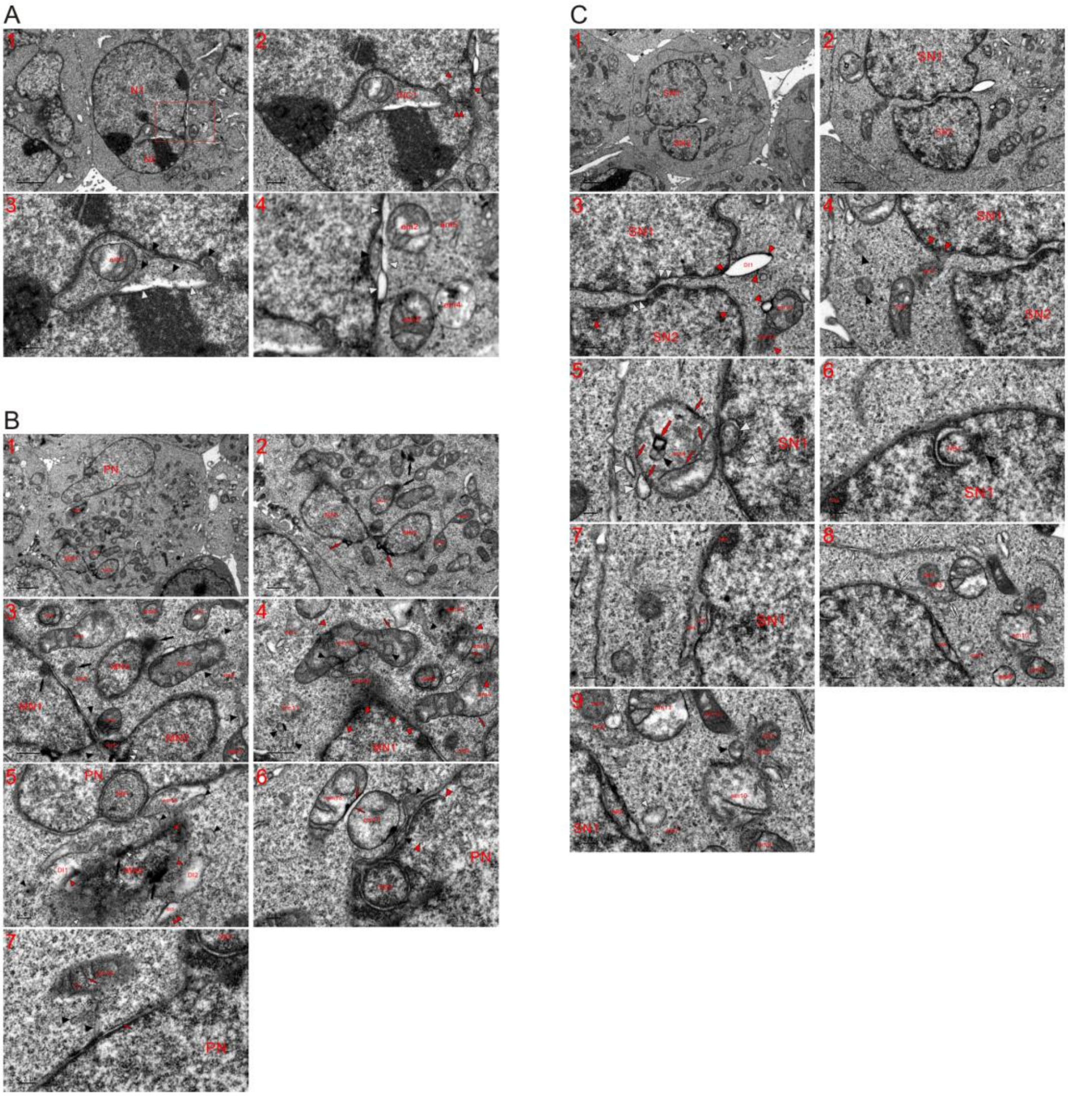


**Fig. S33. Dense particles continued to reassemble for mitochondrial development at the 15 min time point.** TEM was performed on three K562 cells at the 15 min time point (incubation for 15 minutes after switching the cells into new medium), and micrographs revealed that new medium discontinued the nuclear transition of mitochondria, obstructed nuclear fusion, and led to mitochondrial recoveries and re-establishments via the assembly of dense particles. (**A**) Aggregation of dense particles (opposite and double red arrowheads) was going to open INC1 and led to mitochondrial renewal (em1-em3) and re-establishment (em4, em5, black and white arrowheads) both in the cytoplasm and partitioned cytoplasm (INC1 (**A1-A4**). (**A4**): a high magnification image of the inset in (**A1**). (**B**) Aggregation of dense particles occurred at nuclear edges and in the cytoplasm (black and red arrows) and prevented nuclear fusion (MN1 and MN2) by blocking dispersion of the organelles (em1, black and double white arrowheads), which discontinued development of NBD1 into an NPB. Unlike nutrient-limited conditions, new medium assembled dense particles to prevent the organelles from dispersion, and instead, their assembly led to mitochondrial recoveries (such as em1-em7, em14 and em15) and re-establishments (such as em8-em13) depending on the degree of mitochondrial fragmentation. The external aggregation of the particles formed an LMM (small red arrows), and their internal assembly could shape cristae (small red arrows) and SMLBs (black arrowheads in em14). The return of dense particles occurred in MN1 and the cytoplasm (red arrowheads), where mitochondrial reinstatements began with the formation of SMLBs (black and white arrowheads) (**B1-B4**). (**B5** and **B6**) At the edge of the PN, dense particles assembled into bodies of mitochondrial shape (NB1, NB2, opposite red and black arrowheads) and aggregated to recover em17 or re-establish em16, and their aggregation (red arrows) caused separation of the organelles (em17 and em18). At the edge of MN4, which appeared to be in the initial stage of nuclear formation, the aggregation of dense particles and their return to the cytoplasm was observed (black arrows and red arrowheads), and the return of the particles benefited mitochondrial re-establishments (DI1, DI2 and opposite white arrowheads) and enabled DI3 to change into an ERL (double red arrowheads). For completely fragmented mitochondria, their reinstatements began with forming SMLBs (black arrowheads) and the ERL (DI3 and double white arrows). (**B7**) Internal assembly of dense particles formed cristae (red arrows) within em19, which lacked LMM, and formation of dark thread (red arrow) and SMLBs benefited mitochondrial regeneration. (**C**) Aggregation of dense particles divided nuclei (SN1 and SN2) (double white arrowheads) and returned (red arrowheads) to the organelles (DI1, em1-em3 and black arrowheads). Mitochondrial dispersions into nuclei were discontinued (such as em1/2 and em3), em4 was detached from SN1, and the internal aggregation of the particles occurred within either SN1 (white arrowheads) or em4 (black arrowhead, and large and small red arrows). Outside of em4, dense particles assembled into its own LMM (red arrows) and SMLBs (white arrowheads) (**C1-C5**). (**C6-C9**) At the edge of SN1, the particles assembled into nuclear bodies of mitochondrial morphology (NB1-NB5 and black arrowhead), and some of them (NB2-NB4) should have been less fragmented and had just been included in nuclei. In the cytoplasm, mitochondrial renewals (such as em8-em12) and re-establishments (such as em5-em7) occurred, and for those heavily or completely fragmented, their reinstatements began with forming ERLs (white arrows), DEs (DE1 and DE2), SMLBs (black and white arrowheads) and dense strips or sheets (double white arrows).

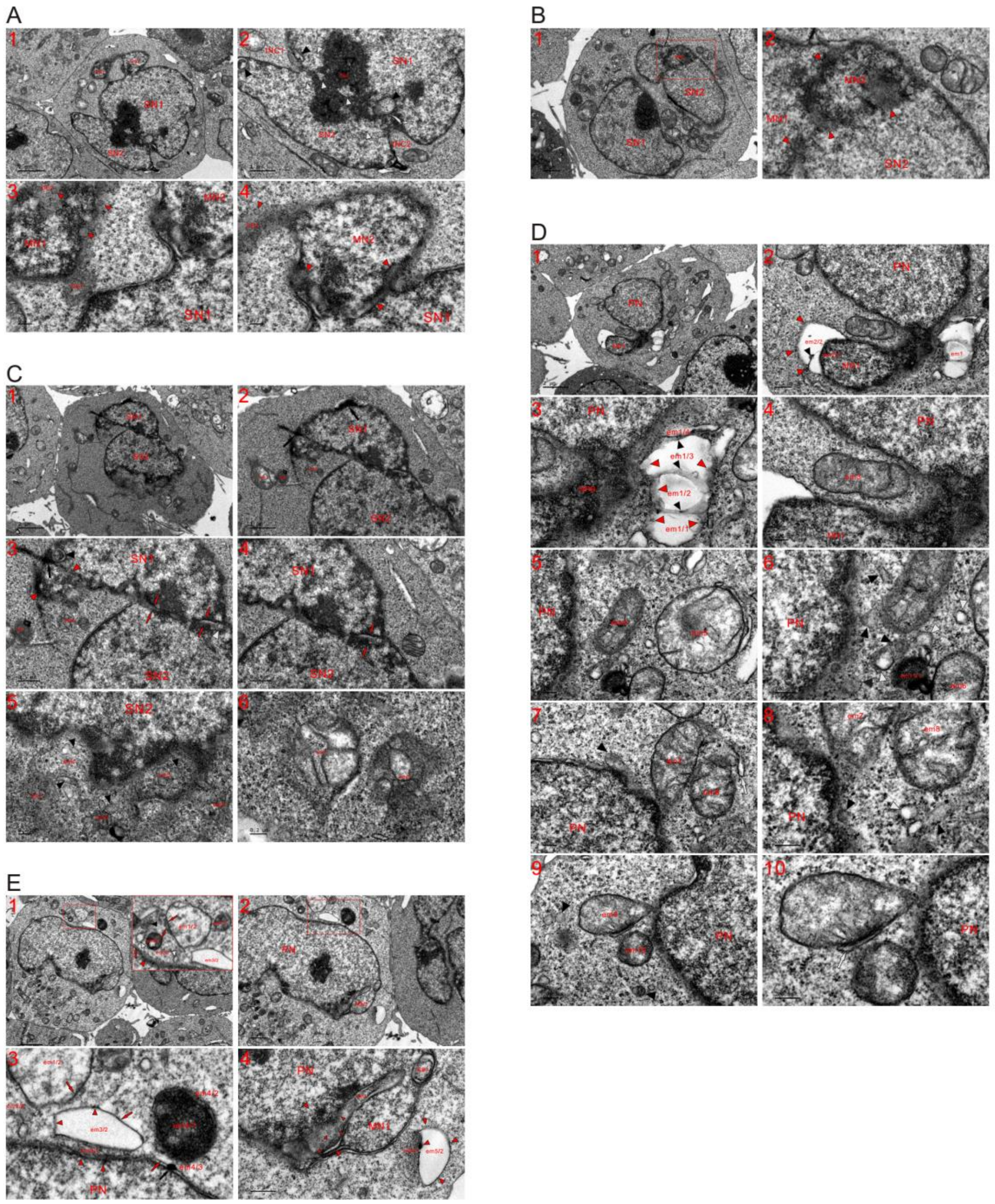


**Fig. S34. Aggregation of heterochromatin (in the form of dense particles) divided nuclei and detached an MN.** TEM was performed on five K562 cells at the 15 min time points, and micrographs revealed that nutrient supplementation stimulated the aggregation of heterochromatin (in the form of dense particles) to separate nuclei and detached an MN concomitantly, leading dense particles to return back into the cytoplasm, where they reassembled for mitochondrial renewal and regeneration. (**A**)

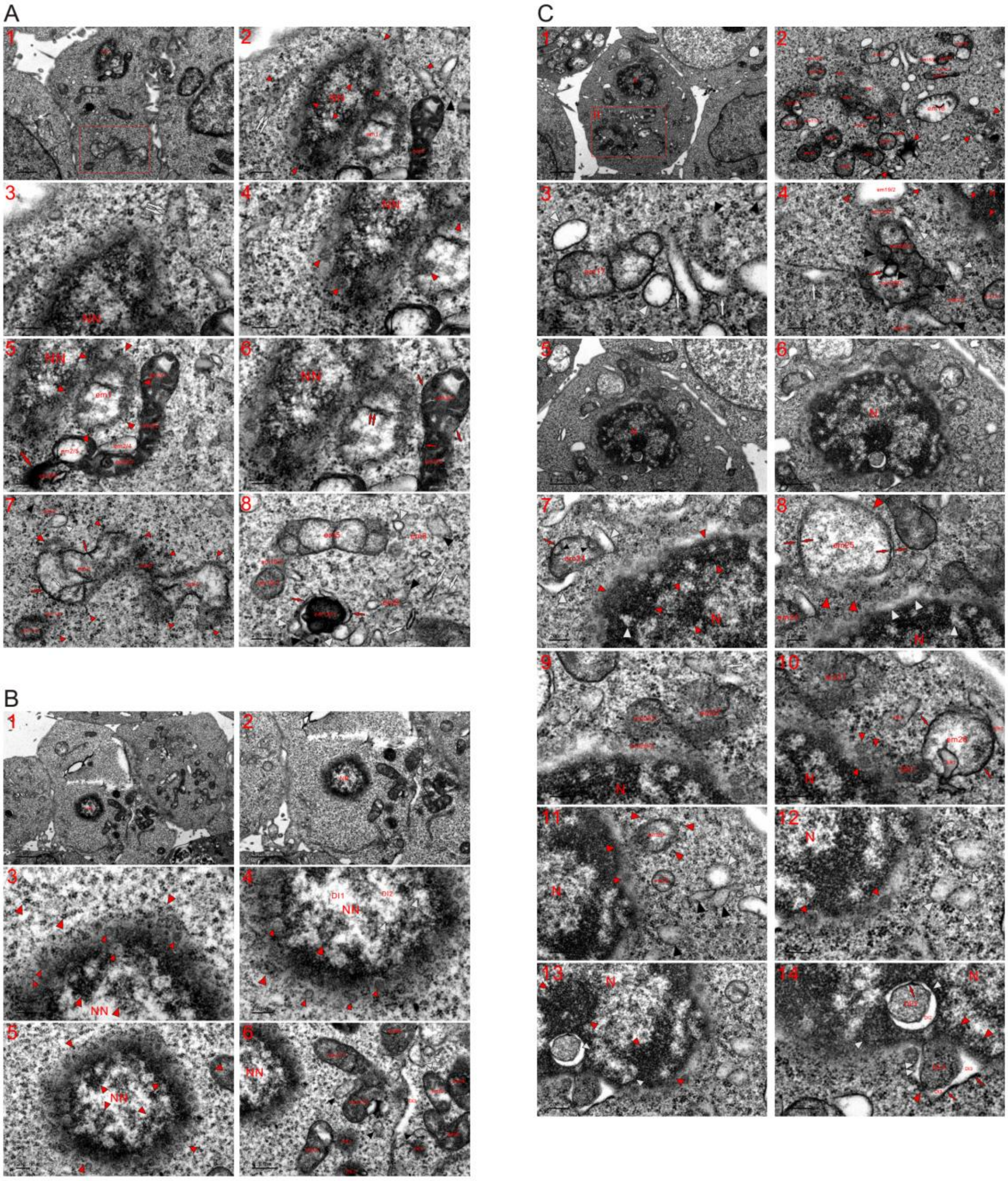


**Fig. S35. Orthodox mitochondria were not formed and were generated in the nucleus**. TEM was performed on three K562 cells at the 15 min time point, and micrographs revealed that nutrient supplementation returned heterochromatin (in the form of dense particles) back into the cytoplasm, where they assembled to restore and regenerate orthodox mitochondria (**A-C**). (**A**) The same cell as shown in **Figure 4E**. New medium discontinued growth of the nucleus, which was membrane-less, oval-shaped and at its initial stage of formation (NN). Instead of dispersion for nuclear growth, nutrient supplementation led to mitochondrial recovery (em1) and re-establishment (opposite red arrowheads) by forming the ERL (red arrow) or precursors of ER (double white arrows) at the nuclear edge (**A1-A3**). (**A4-A6**) Mitochondrial fragmentations reversed to their renewals by returning dense particles (red arrowheads and arrows), and VLGs appeared at the edge of NN and within em1 (small red arrowheads). The aggregation of the particles induced by new medium eventually detached em1 from either NN or em2, which had been condensed and elongated and was going to separate via internal or further assembly of dense particles (em2/1-em2/6 and large red arrow). Double red arrows in em1: the particles were being assembled into cristae. (**A7** and **A8**) The return of dense particles (red arrowheads) also occurred in the cytoplasm for mitochondrial recovery (such as em3-em5) and re-establishment (em6-em9, and black and white arrowheads), and their return caused mitochondrial recombination (em10/1 and em10/2; em11/1 and em11/2; em12/1, red arrows, black and white arrowheads). (**B**) The nucleus in this cell was small, membrane-less, spherical-shaped and at the early stage of its formation (NN), and a new medium caused heterochromatin (in the form of dense particles) to aggregate at its edge and the particles to return themselves to the cytoplasm, where they formed clusters (large red arrowheads). At its edge, a large amount of VLGs (small red arrowheads) appeared, which turned into clusters or dispersed when they were somewhat far away from the NN. Along with the aggregation of dense particles within the nucleus (red arrowheads), electron-transparent areas or spots (such as DI1, DI2 and white arrowheads) were created (**B1-B4**). (**B5** and **B6**) Aggregation of the particles induced by nutrient supplementation was going to separate the NN (opposite red arrowheads), discontinued mitochondrial fragmentation or dispersion and led to mitochondrial recovery (such as em1-em6) and re-establishment (DE1-DE3, DI3, black and white arrowheads). (**C**) Although it was relatively large (N), the nucleus was still membrane-less, and the aggregation of dense particles within it was observed. At another site of the cytoplasm, mitochondria should have been fragmented to initiate nuclear formation (em1-em5), which was discontinued by new medium supplement. With nutrient supplementation, condensed mitochondria separated (such as em7 and em8; em9 and em10) and recombined (such as em5/1 and em5/2; em6/1 and em6/2; em11/1-em15/1 and em11/2-em15/2) for mitochondrial recovery or re-establishment. Less fragmented mitochondria were being renewed (such as em16-em18), and mitochondrial reinstatements occurred in heavily or completely fragmented mitochondria (such as em1-em4 and opposite red arrowheads). The return of dense particles first formed vesicles (white arrowheads), ERLs (white arrows) and SMLBs (black arrowheads), some of the vesicles changed into small mitochondria (SMMs) or SDBs to participate in the generation of large mitochondria (**C1-C3**). (**C4**) Electron-lucent em19/2 evolved into an orthodox mitochondrion by recombining with em19/1 and the return of dense particles, which also happened at the nuclear edge (small red arrowheads). Their internal (em20/1, em20/2, red arrow and black arrowheads) and external aggregation led to the mitochondrial renewal of em20, and mitochondrial re-establishments began with forming SMLBs or SMMs (black and white arrowheads) and the ERL (white arrow) for those heavily or completed fragmented (such as em22 and em23). (**C5-C8**) Instead of dispersion, electron-opaque em24 was recovered via internal (black arrowheads) and external (red arrow and white arrowheads) assembly of the particles, and the return of dense particles (opposite red arrows and red arrowheads) benefited the mitochondrial renewal of lucent em25. At the nuclear edge and within it, their assembly (opposite red arrowheads) concurrently created electron-transparent areas or spots (white arrowheads). (**C9** and **C10**) Dispersion of the organelles was blocked, and the return of the particles led to mitochondrial reinstatements (em26/1 and em26/2; em27), and their internal (such as ER1) and external (red arrows, DE1 and ER2) aggregations occurred in em28. Rather than diffusion for nuclear growth, the formation of DE1, VLGs and the ER3 benefited mitochondrial re-establishments. (**C11** and **C12**) Aggregated particles returned from the nucleus and the surroundings to em29 (red arrowheads), which lacked its own LMM and was being recovered, and formation of VLGs (small red arrowheads) and SMLBs (em30, black and white arrowheads) promoted mitochondrial regeneration in the cytoplasm. (**C13** and **C14**) Within the nucleus and at its edge, aggregation of dense particles formed dense (opposite red arrowheads) and relatively dilute (opposite white arrowheads) structures or bodies. The diffusion of DE2 in the nucleus discontinued, and instead, the aggregation of the particles from all directions condensed it, concurrently enlarging the lunar halo-shaped DI2. During the further assembly of dense particles of its own and the surroundings (opposite white arrowheads) for mitochondrial development, both DE2 and DI2 appeared in the cytoplasm. The dispersion of DE3 into the nucleus discontinued, and the assembled dense particles returned to it and DI3 from either the nucleus or cytoplasm (red arrows and arrowheads), concurrently forming DI3 and electron-lucent intervals (white arrowheads). Once it separated from the nucleus, DE3 recombined with electron-lucent structures for mitochondrial re-establishment, and the formation of either dark threads via the aggregation and linearization of the particles or the lucent interval (double white arrowheads) promoted its separation from the regenerated organelles.

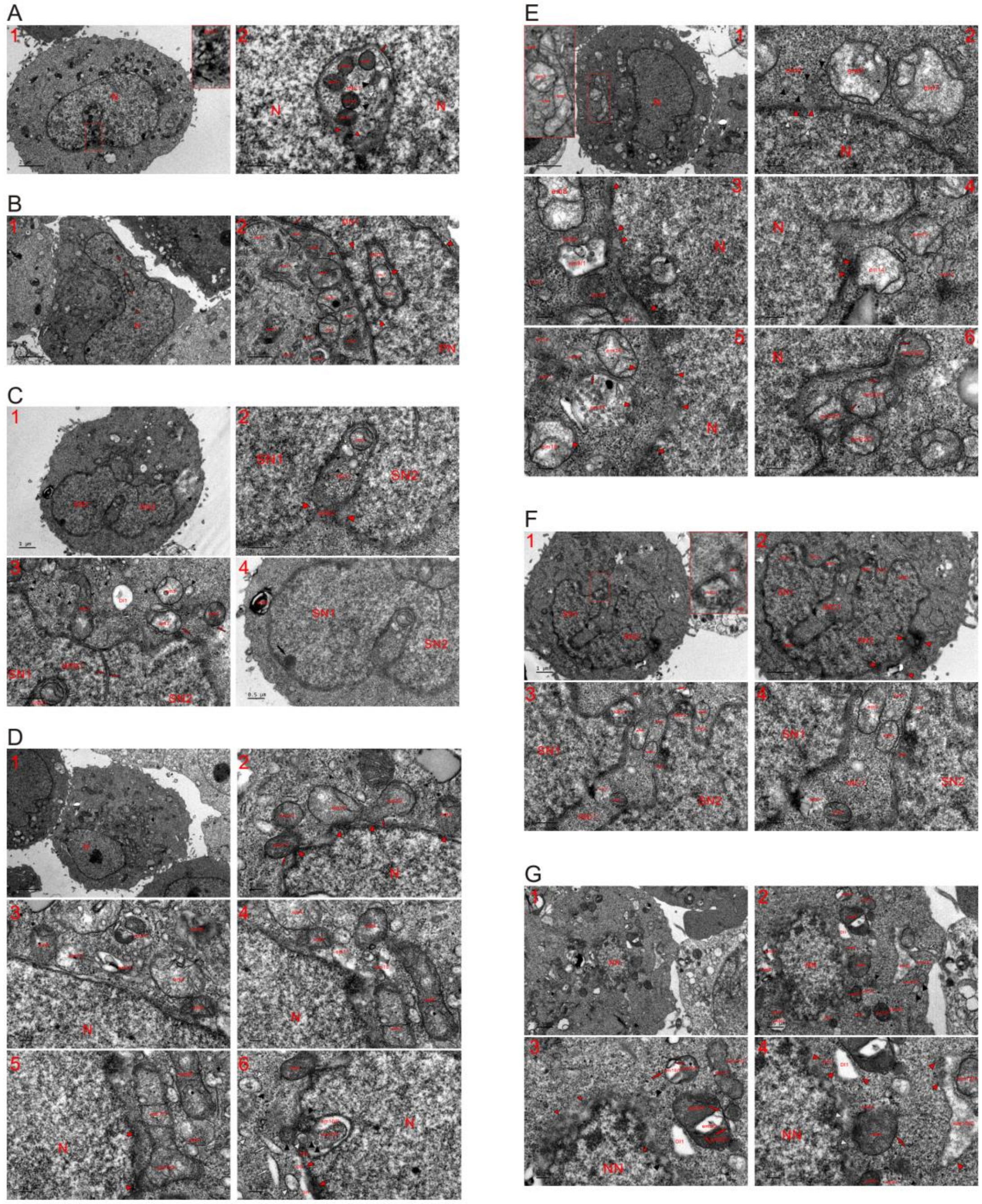

**Fig. S36. Nutrient supplementation reversed the nuclear transition of the organelles to mitochondrial recovery and re-establishment in HeLa cells**. TEM was performed on seven HeLa cells at the 15 min time point, and micrographs revealed that stimulation with new medium blocked nuclear growth by reversing dense particles to reassemble for mitochondrial recoveries and re-establishments. (**A**) High magnification image of the inset shows that INC1 was opened by aggregation of dense particles (black arrows) and dense particles reassembled for mitochondrial renewals (such as em1-em3) and reinstatements (red arrow, and black, white and small red arrowheads) (**A1** and **A2**). (**B**) Under nutrient-limited conditions, four-sided nuclear formation had partitioned the cytoplasm to form INC1, which became more obvious with challenge of new medium, and nutrient supplementation led to mitochondrial recoveries (em1-em7) and re-establishments (em8-em15). Red arrows: dark threads; white arrows: ERLs (**B1** and **B2**). (**C**) INC1 was opened through breaking up NPB2 by aggregation of dense particles (opposite red arrowheads), whose assembly formed nuclear tubules (opposite red arrow) and ERLs (white arrows). Within INC1, mitochondrial renewal (em1) and regeneration occurred (white arrows, black and white arrowheads), and formation of ERLs (white arrows) and SMLBs (black and white arrowheads) promoted mitochondrial re-establishment in the cytoplasm. For less fragmented mitochondria (such as em2-em5), the particles assembled to renew the organelles, and their return enabled DI1 to evolve into a mitochondrion. The dispersion of em6 into SN1 was blocked, em6 was restored into a membrane whorl structure via the external and internal assembly of dense particles, and their aggregation formed a dense spherical body (black arrow) at the edge of SN1 (**C1-C4**). (**D**) Rather than dispersing into the nucleus, nutrient supplementation led to mitochondrial renewal (such as em1-em8) and re-establishment (such as em9-em15), concurrently promoting the separation of the organelles (em1/1, em1/2; em2/1, em2/2; em7/1, em7/2). The return of dense particles from the nucleus detached the organelles by forming the LMM (red arrows), and their aggregation within the organelles led to mitochondrial separation (such as em8 and em9) (**D1-D5**). (**D6**) Incomplete nuclear transition of the organelle (em10) had evolved into the appearance of an INC at the nuclear edge under nutrient-limited conditions, and it was being re-established for mitochondrion via recombination (em10/1 and em10/2) and reassembly of dense particles (black arrowheads). The development of the NBD into an NPB discontinued, and instead, the assembly of dense particles promoted mitochondrial re-establishments (DI1, DI2, white arrows, and black and white arrowheads) demonstrated via their return (red arrowheads). (**E**) High magnification image of the inset revealed that mitochondria had been fragmented to group together under nutrient-limited conditions, and new medium was going to separate the organelles through mitochondrial recoveries (em1 and em2) and re-establishments (em3 and em4) (**E1**). (**E2**) Mitochondrial re-establishment of em5 began with the assembly of dense particles into SMLBs (black arrowheads), and both organelles (em6 and em7) were recovered. (**E3** and **E4**) Within the nucleus, the reassembly of dense particles for mitochondrial reinstatement was observed (black and white arrowheads), and their aggregation resulted in their return to the cytoplasm (red arrowheads), where either mitochondrial renewals (em8, em14 and em15) or regeneration occurred (such as em9/1, em9/2, and em10-em13). (**E5** and **E6**) Along with mitochondrial recoveries (em16-em18), mitochondrial re-establishments occurred between nucleus and the organelles (opposite red arrowheads), and a group of the fragmented organelles (em19-em21) began to separate along with the re-establishments. The external assembly of dense particles formed the LMM of em17 (red arrow) and their internal aggregation led it to look like an MVB. At the nuclear edge, new medium reversed the dispersion of the organelles into the nucleus to promote mitochondrial renewals and separations (em22/1, em22/2; em23/1, em23/2). (**F**) New medium led the NPB to the four-sided assembly of dense particles, which caused the disappearance of the partitioned cytoplasm (INC1), and a high magnification image of the inset in (**F1**) reveals that mitochondrial re-establishments occurred at the nuclear edge (em1, NBD1 and white arrow) (**F1** and **F2**). (**F3** and **F4**) At the openings and within INCs, either mitochondrial renewals (such as em5-em8) or re-establishments (such as NBD1, NBD2, NB1, NB2 and em1-em4) occurred. (**G**) The nucleus was at the early stage of formation, and its growth discontinued upon nutrient supplementation, which reversed the nuclear transition of the organelles to mitochondrial recoveries (such as em7-em11) and re-establishments (such as em1-em6). Reinstatements of the organelles occurred by the recombination of dense and lucent parts (such as em3/1 and em3/2; em1 and em2; DE1 and DI1) with the help of externally assembled dense particles (red arrowheads). At the nuclear edge, nutrient supplementation increased the formation of VLGs (small red arrowheads) and discontinued the dispersion of SMLBs (black arrowheads), both of which were further processed for mitochondrial regeneration. In order to change into orthodox mitochondria, dense mitochondria (such as em7/1-em13/1) usually and first recombined with lucent or dilute parts (such as em7/2-em13/2) of the organelles with the help of the external assembly of dense particles (red arrowheads and arrows), and this recombination promoted the further assembly of the particles within mitochondria. The formation of ERL (white arrow) and SMLBs (black arrow and white arrowheads) together with dilute em12/2 led to the generation of additional mitochondria (em14/2 and em15/2).

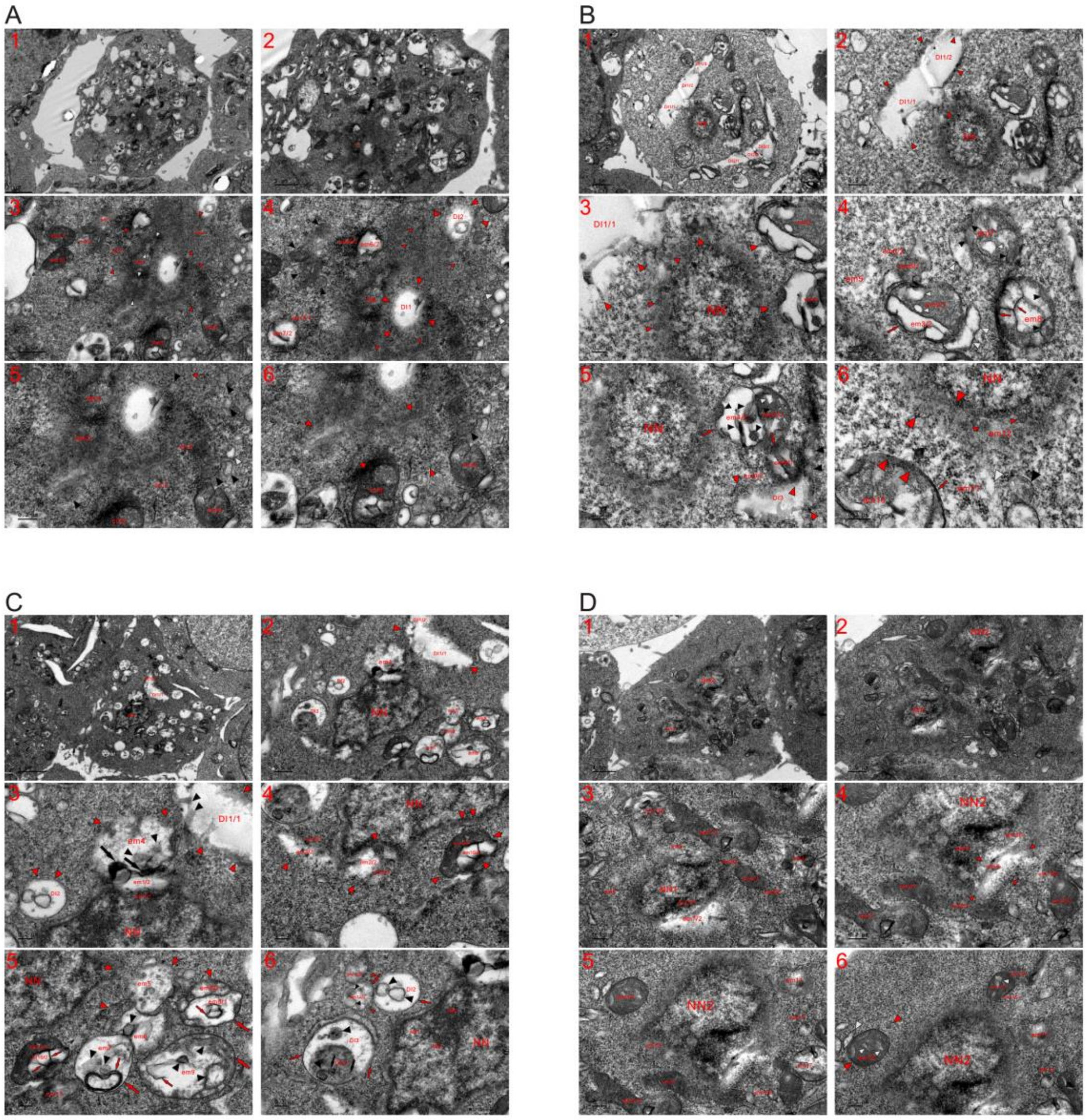

**Fig. S37. Nutrient supplementation reversed** the **nuclear transition of the organelles to mitochondrial** recovery **and re-establishment in HEK293T cells.** TEM was performed on four HEK293T cells at the 15 min time point, and micrographs revealed that fresh medium discontinued nuclear initiation and formation by reassembling dense particles for mitochondrial renewal and re-establishment. (**A**) It was difficult to discern nuclei at low magnification in the cell and cytoplasm filled with mitochondria, which had been fragmented or became electron transparent through aggregation of dense particles to the peripheries (**A1**). (**A2-A6**) Adjacent to DI1, there appeared a structure of nuclear appearance (NN) and was separated into bodies of mitochondrial morphology (NN1, NN2 and opposite red arrowheads). Fresh medium led to the aggregation of dense particles in both NN and cytoplasm, where they returned back into DIs (DI1 and DI2) and the organelles (such as em6/1, em6/2, em7/1, em7/2, and em8-em11) through their own assembly. For the organelles that had been completely fragmented and dispersed into particles, mitochondrial re-establishments (such as em1-em5 and opposite red arrowheads) began with the formation of VLGs (small red arrowheads) and SMLBs (black arrowheads) via reassembly. In the cytoplasm, the formation of SMLBs benefited mitochondrial regeneration (black and white arrowheads). (**B**) The large DIs (such as DI1 and DI2) should have been derived from more than one mitochondria (DI1/1-DI1/3 and DI2/1-DI2/3), and new medium caused dense particles (large red arrowheads) to return back into DIs and nascent or emerging mitochondria (such as em3 and em4). Small red arrowheads: VLGs (**B1-B3**). (**B4** and **B5**) New medium discontinued mitochondrial fissions or dispersions, recombined the separated mitochondria together (such as em3/1, em3/2 and large white arrowheads; em4/1, em4/2, red arrow, black and white arrowheads; em7, large white arrowhead, small black and white arrowheads; em8, large and small red arrows, black arrowheads; em9/1 and em9/2) and outlined the organelles via the external assembly of dense particles into LMMs (large red arrows). Mitochondrial re-establishment occurred in electron-dilute DI3 via the return of dense particles (large red arrowheads) and the formation of an LMM (red arrow) led to the separation of em9/1 from em4/1. (**B6**) The return of dense particles (red arrowheads) benefited mitochondrial reinstatement between em10 and nucleus (opposite red arrowheads), concomitantly shaping the LMM (red arrow). For those completely fragmented at the nuclear edge, mitochondrial regeneration (em11 and em12) began with the formation of SMLBs (black and white arrowheads) or VLGs (small red arrowheads). (**C**) Nutrient supplementation recombined the separated organelles (such as em1/1-em3/1 and em1/2-em3/2; em4, black arrows and arrowheads; em10/1, em10/2) for mitochondrial re-establishments with the help of externally assembled dense particles (red arrowheads), and unlike what had happened under nutrient-limited conditions, their presently internal (small red arrows and black arrowheads) and external (large red arrows and red arrowheads) assembly reversed the fragmentation of the organelles to promote mitochondrial recovery (such as em8 and em9) and reinstatement (em5-em7 and em11). The return of dense particles (red arrowheads) led to the separation of the large DI1 (DI1/1 and DI1/2), and the internal aggregates (black arrowheads) dispersed into electron-lucent parts with nutrient supplementation (**C1-C5**). (**C6**) External aggregation of dense particles outlined DIs (DI2 and DI3) by forming LMMs (red arrows), and a large DE3 in DI3 changed into SMLBs through the further assembly of the particles (black arrows and arrowhead), whose reassembly led to the intermixing of these SMLBs with the electron-transparent parts. Their internal (black arrowheads) and external (red arrow and arrowheads) aggregation occurred in em12/2, which was being recombined with em12/1. Within the NN, nuclear bodies (NB1-NB3) were being formed along with the aggregation of dense particles. (**D**) Between or around nuclei, dispersion of mitochondria reversed to mitochondrial recoveries (such em1/1-em3-1, em14, em15/1, em16 and em17) and re-establishments (em1/2-em3/2, em3/3, em4-em13 and em15/2) upon challenge with new medium. At the nuclear edge, a combination of the separated mitochondrial parts was demonstrated (such as em7/1 and em7/2), and the return of dense particles (red arrowheads) allowed for to the mitochondrial regeneration of the electron-transparent em8 with help of dispersion of the internal aggregate. Internal aggregation (small white arrows) occurred in dense mitochondria (such as em14/1 and em16), which changed into orthodox ones through recombination (em14/1-em14/3; em16, large white and opposite red arrowheads) (**D1-D6**).

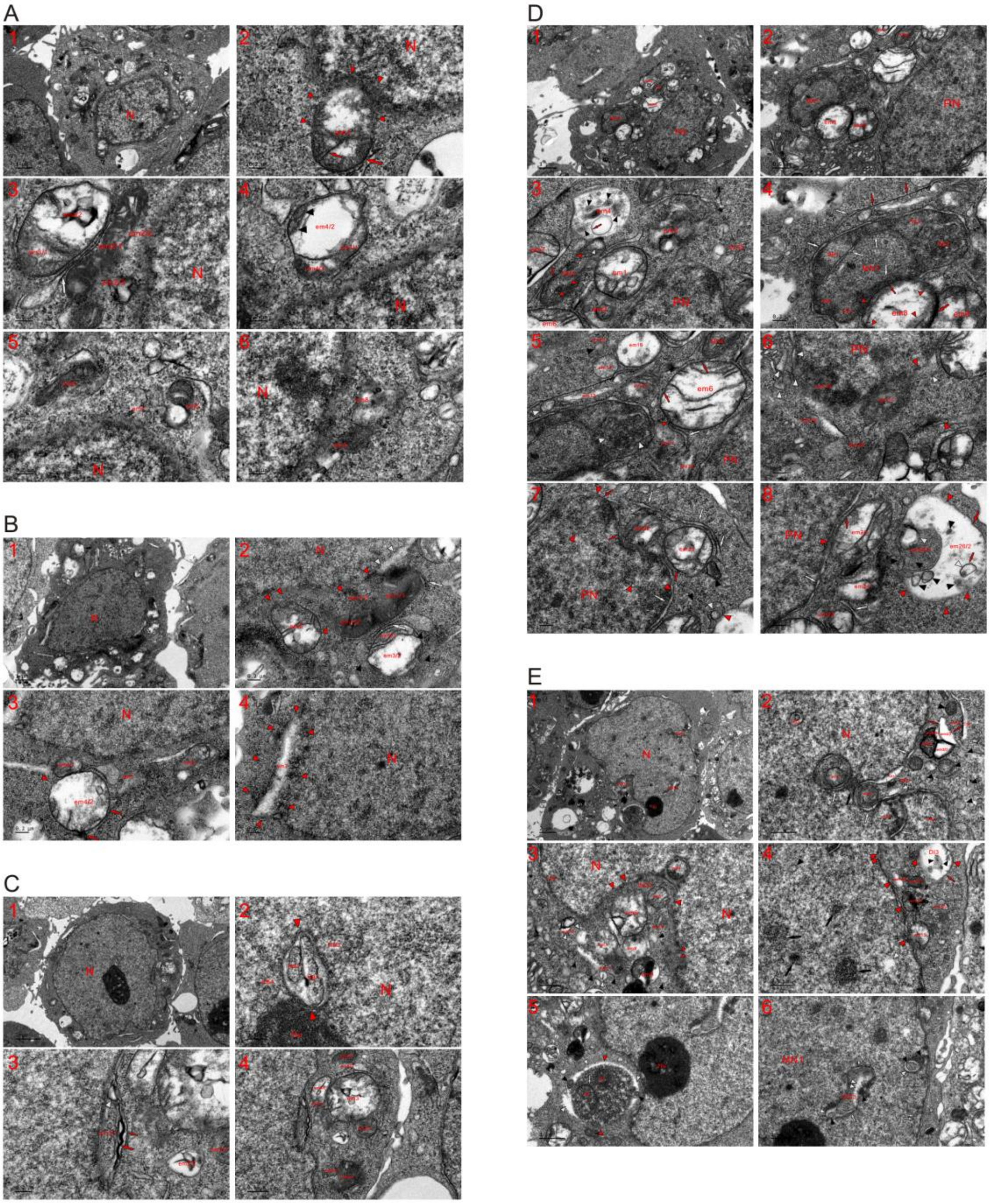


**Fig. S38. New medium converted nuclear formation to mitochondrial renewal and regeneration in HEK293T cells.** TEM was performed on five HEK293T cells at the 15 min time point, and stimulation with new medium discontinued nuclear growth by blocking mitochondrial fissions and promoted either recovery or regeneration of the organelles in the cytoplasm. (**A**) The nucleus lacked tubule-NE, and the dispersion of em1 reversed its recovery by returning dense particles from the nucleus and the surroundings (red arrowheads) upon challenge with new medium, and their assembly formed an LMM (large red arrow) and cristae (small red arrow) (**A1** and **A2**). (**A3-A6**) At the nuclear edge, mitochondrial renewals were performed by recombining the separated parts (em2/1-em2/3; em3/1 and em3/2; em4/1-em4/3 and black arrowheads), and formation of the ERL (white arrowhead) led to separation of em2 and em3. For heavily or completely fragmented mitochondria, their re-establishments were observed (such as em5-em9). (**B**) A large part of em1 had evolved to be a dense phase (em1/1 and em1/2) without LMM, and em1/3 already became nuclear under nutrient-limited conditions. Nutrient supplementation returned dense particles to it (white arrow and red arrowheads) for re-establishment with the help of internal assembly. The return of the particles created a dilute interval (white arrowheads) at the nuclear edge and caused em2 to separate from the nucleus (red arrowheads), and mitochondrial recombination occurred in em3 for renewal (em3/1, em3/2, white arrow and black arrowheads) (**B1** and **B2**). (**B3** and **B4**) The organelle was being recovered by recombining the separated parts (em4/1 and em4/2) and the return of dense particles (red arrows and arrowhead), and heavily or completely fragmented mitochondria was reinstated (such as em5 and em6). The organelle (em7) had changed into a DI through the aggregation of dense particles at the periphery under nutrient-poor conditions, and new medium assembled dense particles allowing their return (red arrowheads) into the electron-lucent em7 for its mitochondrial re-establishment. (**C**) Within the nucleus and very close to Nu, the aggregation of dense particles induced by nutrient supplementation formed rod-shaped nuclear bodies (NB1 and NB2), concurrently shaping an INC structure (opposite red arrowheads). Adjacent to the INC, bodies with mitochondrial morphology were being formed (NB3 and NB4) (**C1** and **C2**). (**C3** and **C4**) Mitochondrial fragmentation or dispersion was discontinued upon supplementation with new medium, which led to mitochondrial re-establishment and recovery (such as em1/1, red arrows and white arrowheads; em2/1 and em2/2; em3; em4/1-em6/1 and em4/2-em6/2). (**D**) Three nuclei (PN, MN1 and MN2) appeared in the cell, and MN2 looked like a mitochondrion and displayed similar appearance as that of either MN1 or the PN. New medium blocked its growth and returned dense particles (red arrowheads and arrow) to the cytoplasm along with their internal assembly (white arrowhead and arrows). Around it and at the edge of the PN, mitochondrial recoveries (em1, em6 and em7) and re-establishments (such as em2-em5) were observed. Electron-transparent em4 should have become a DI and dispersed in the cytoplasm under nutrient-limited conditions, and presently, the return of the particles outlined it and led to its mitochondrial regeneration along with their internal reassembly (red arrow and black arrowheads) (**D1-D3**). (**D4** and **D5**) Within MN1, aggregation of dense particles led to the formation of nuclear bodies of mitochondrial morphology (NB1-NB5) and ERLs (large white arrows), and in NB2, further assembly of the particles into SMLBs was demonstrated (white arrowheads). Adjacent to MN1, new medium reversed the diffusion of the organelles to mitochondrial renewals or re-establishments by the return of dense particles (em6, em8, red arrows and arrowheads), and their aggregation separated em8 from either MN1 (small red arrow) or em9 (large red arrow). The formation of a dark thread (small red arrows), ERLs and SMLBs (black and white arrowheads) benefited mitochondrial re-establishments (such as em11-em15), and the external return and internal reassembly of dense particles caused the electron-lucent em16 change back to a mitochondrion. (**D6**) At the edge of the PN, mitochondrial re-establishment and regeneration were evidenced due to the assembly of dense particles (such as em17-em20, white and opposite red arrowheads), and compared to em18, em17 was more advanced in mitochondrial development. (**D7** and **D8**) Nutrient supplementation discontinued dispersions of the organelles into the PN and led to mitochondrial recoveries (such as em21 and em25) or reinstatements (such as em22-em24 and opposite red arrowheads), and for those heavily or completely fragmented organelles, mitochondrial re-establishments began with forming ERLs (white arrows), dark threads (red arrows) and SMLBs (black and white arrowheads). The return of dense particles from the PN formed the LMM (red arrows), which detached mitochondria from the nucleus (such as em21 and em25). Previous aggregation of dense particles had separated em26 into electron-opaque em26/1 and electron-lucent em26/2, concomitantly forming the internal aggregates (small red arrow, black and white arrowheads) and leading to its dispersion under nutrient-limited conditions; nutrient supplementation recombined the separated counterparts (em26/1 and em26/2) and returned dense particles back to the organelle (red arrowheads). The return of the particles outlined it by forming the LMM, and their reassembly formed SMLBs within dense em26/1 and caused the internal aggregates to disperse into electron-transparent em26/2. (**E**) Nutrient supplementation caused INCs (INC1-INC3) to become more obvious, and within the nucleus and cytoplasm, mitochondrial recoveries (such as em1 and em2) and re-establishments (NB1-NB3, em3, em4, DI1, DI2, white arrows, red arrows, black and white arrowheads). Between em1 and em2, the aggregation of dense particles (black arrow) enlarged INC1, concurrently causing em1 to enter the cytoplasm. Mitochondrial recombination occurred to re-establish em3 (em3/1-em3/3) and em4 (em4/1 and em4/2) (**E1** and **E2**). (**E3**) At the nuclear edge, nuclear bodies with a mitochondrial shape (NB4 and NB5) appeared, spherical NB4 entered INC2, and the return of dense particles from the nucleus (large red arrowheads) led to mitochondrial recoveries and detachments (such as em5 and em6). For heavily or completely fragmented mitochondria, mitochondrial re-establishment (em7-em12) began with the formation of VLGs (small red arrowheads) and SMLBs (black and white arrowheads). (**E4**) Within the nucleus, the aggregation of dense particles was observed (black arrows, black and white arrowheads), and at the nuclear edge, mitochondrial recombination (em13/1 and em13/2), regeneration (em14 and em16) and renewal (em15) were exhibited by the return of dense particles from the nucleus (large red arrowheads) and the surroundings. The return of the particles (red arrowheads) outlined DI3 (red arrow) and reversed it to change into a mitochondrion with help of the internal aggregates (small black arrowheads). (**E5** and **E6**) Adjacent to Nu and in the cytoplasm, a large globular body (opposite red arrowheads) appeared, which should have been derived from the fragmented organelles and changed into an MN to fuse with the nucleus. New medium caused dense particles to assemble both inside and outside of it, concomitantly blocking its nuclear development. External aggregation of the particles formed SMLBs (black arrowheads) and a lunar halo-like structure (white arrowheads), and their internal assembly created bodies with a mitochondrial shape (B1 and B2) and shaped the ERL (white arrow). Within the nucleus, the aggregation of dense particles formed electron-opaque and electron-lucent structures or bodies (black and white arrowheads; double white arrowheads and white arrows), concurrently causing them to look like a closed INC (INC3).

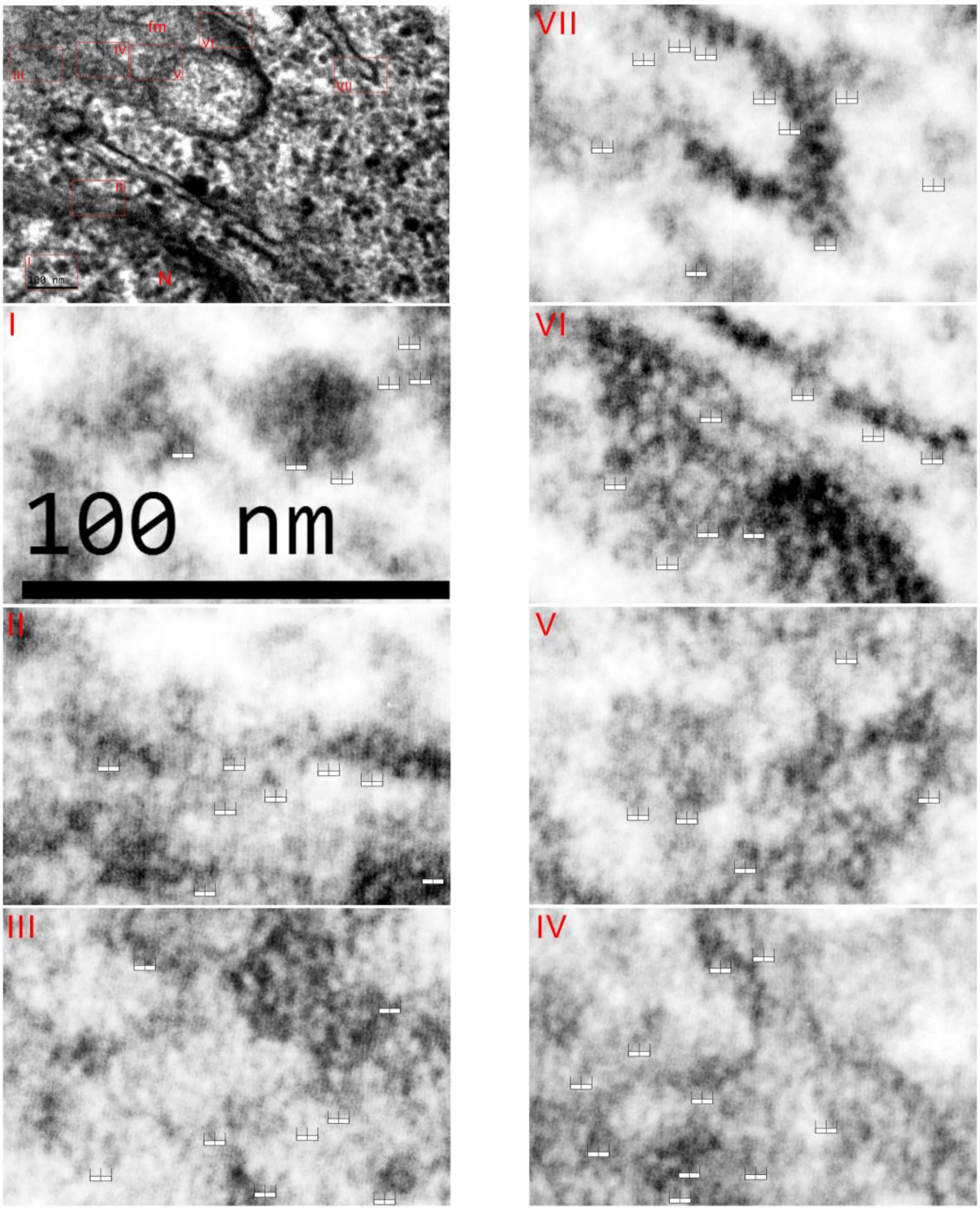


**Fig. S39. Dense particles consisted of microvesicles (MIVs) in K562 cells.** Electron micrographs showing a section of a K562 cell, and high magnification images (I-VII) reveal that MIVs existed in dense particles, which formed the nucleus (N), mitochondrion (fm), ERL (white arrowheads) and cytoplasmic network; the comb-like bar indicates the MIVs.

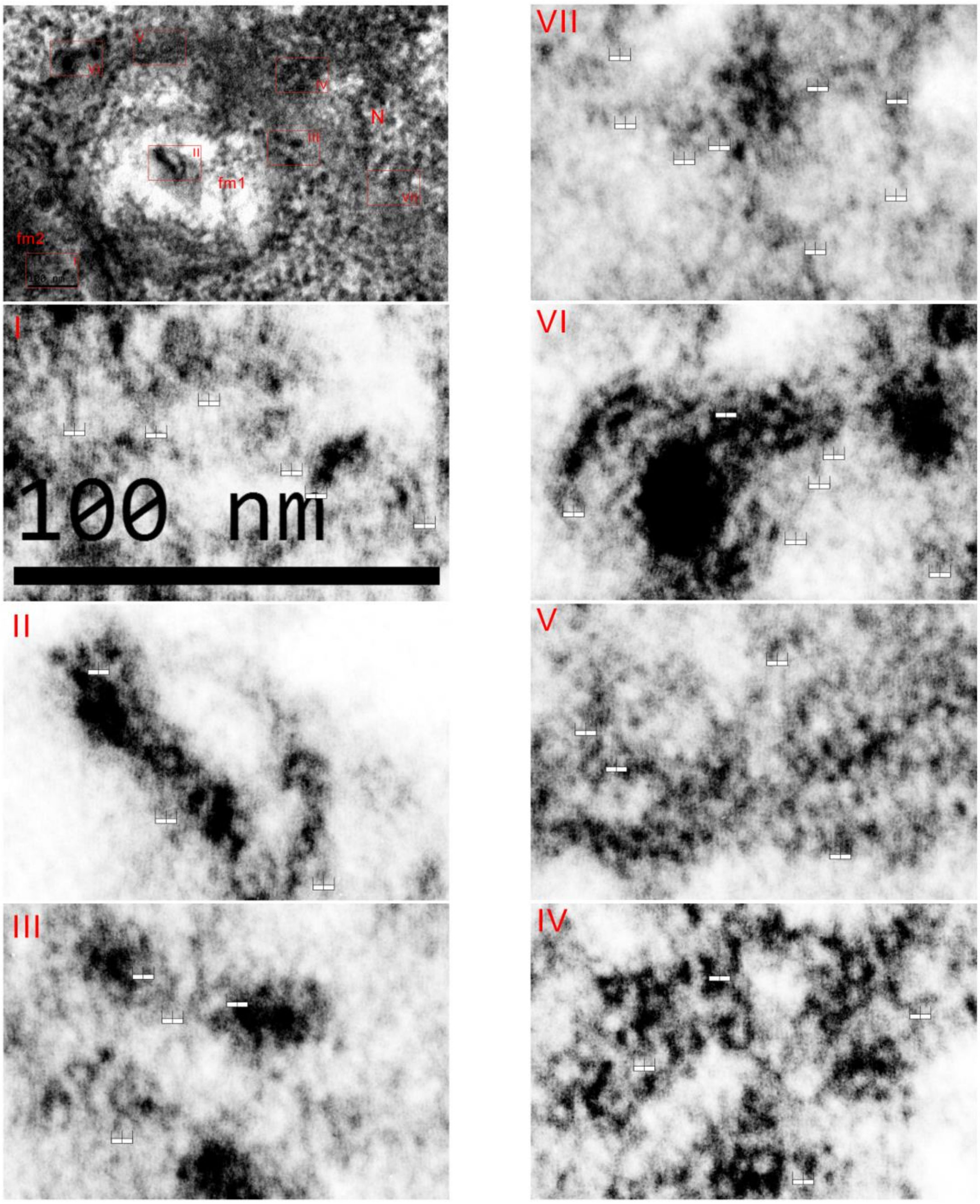


**Fig. S40. Dense particles consisted of MIVs in HeLa cells.** Electron micrographs showing a section of a HeLa cell, and high magnification images (I-VII) reveal that MIVs existed in dense particles, which formed the nucleus (N), mitochondria (fm1 and fm2) and cytoplasmic network; comb-like bar (5 nm) indicates the MIVs.

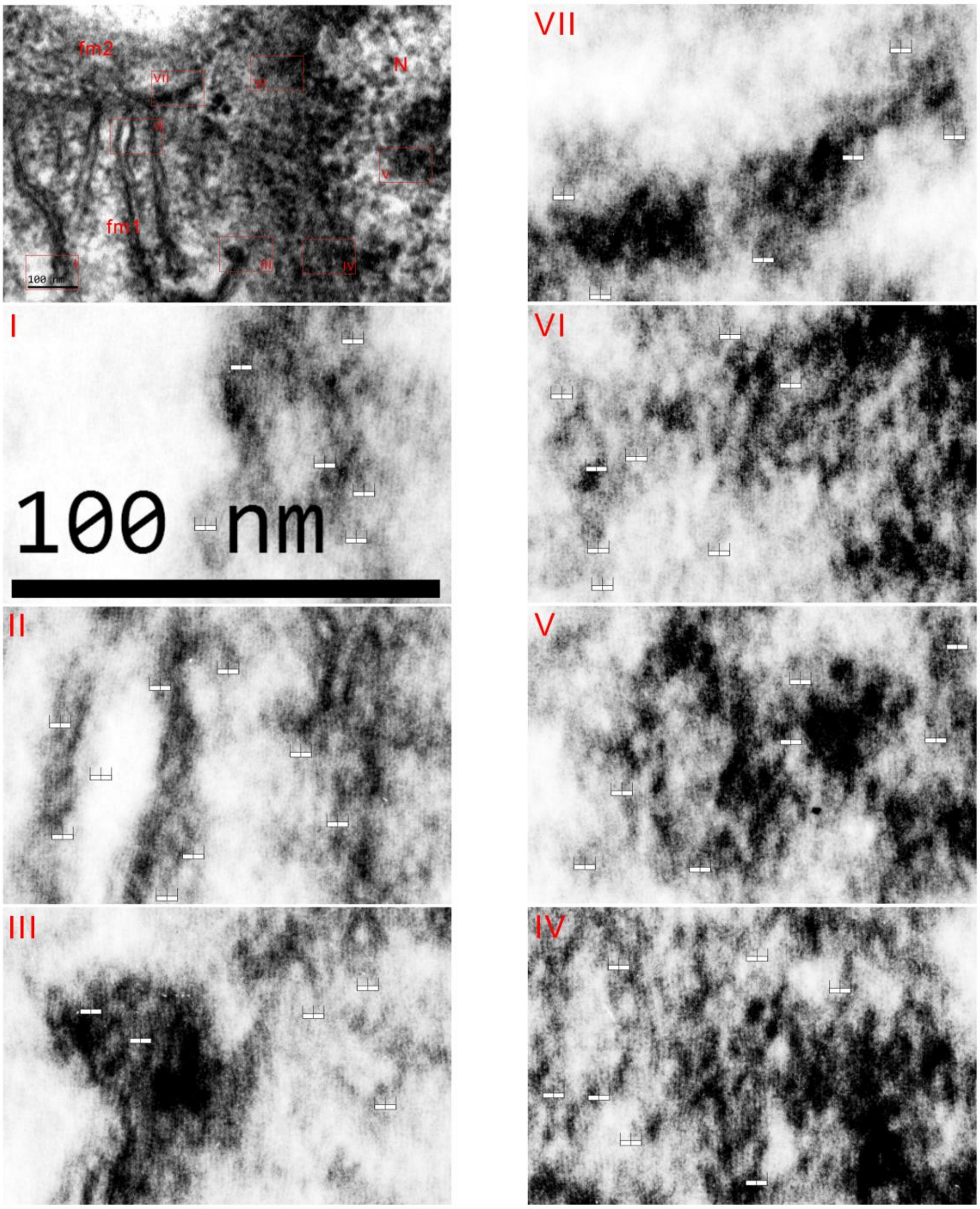


**Fig. S41. Dense particles consisted of MIVs in HEK293T cells.** Electron micrograph showing a section of a HEK293T cell, and high magnification images (I-VII) reveal that MIVs existed in dense particles, which formed the nucleus (N), mitochondria (fm1 and fm2), cristae and cytoplasmic network; the comb-like bar (5 nm) indicates the MIVs.

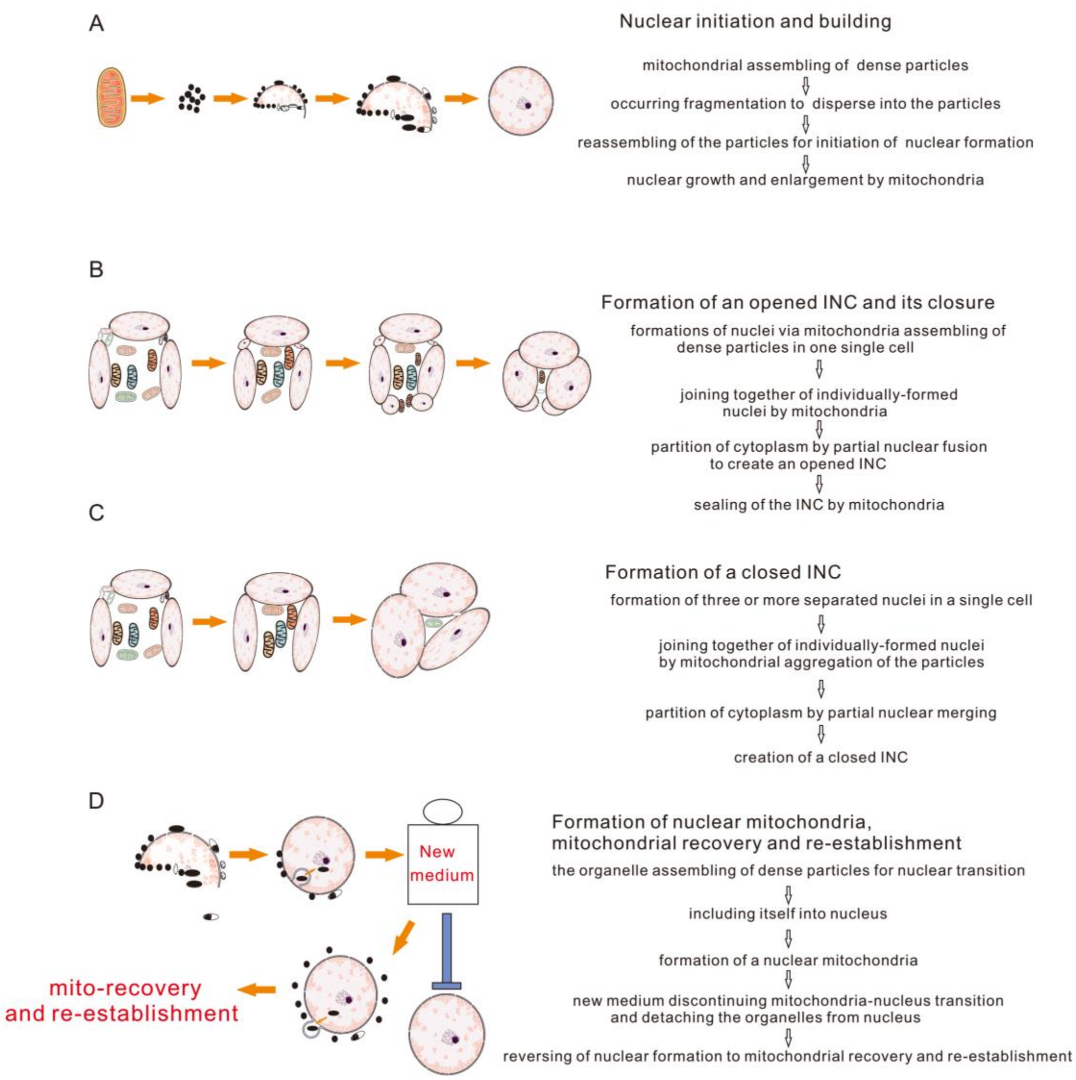


**Fig. S42. Mitochondria fragment and reassemble to initiate the formation and development of the nucleus.** (**A**) Nuclear initiation and building. (**B**) Formation of an opened INC and its closure. (**C**) Formation of a closed INC. (**D**) Formation of nuclear mitochondria, mitochondrial recovery and re-establishment.
